# Matrix stiffening toolbox: dynamic hydrogels for three-dimensional cell culture with real-time cell response

**DOI:** 10.64898/2026.03.25.714233

**Authors:** Eden M. Ford, Samantha E. Cassel, Bryan P. Sutherland, Samantha L. Swedzinski, April M. Kloxin

## Abstract

Extracellular matrix (ECM) mechanical properties regulate tissue homeostasis and disease progression, with persistent ECM stiffening serving as a hallmark of fibrosis; yet, the early transition from healthy to diseased tissue remains poorly understood. Dynamic three-dimensional (3D) tissue models that capture early-stage stiffening are needed to investigate cellular responses during disease initiation. This work presents an innovative platform for studying cell responses in 3D environments undergoing active matrix stiffening. A bioinspired synthetic ECM incorporates collagen-mimetic peptides and employs sequential, non-terminal strain-promoted azide–alkyne cycloaddition (SPAAC) reactions to enable controlled increases in matrix stiffness over physiologically relevant timescales. Alternating polymer incubations produce a 2.5-fold increase in storage modulus over 72 hours, modeling the mechanical transition from healthy to early-stage fibrotic lung tissue. Live-cell reporter fibroblasts enable real-time monitoring of alpha-smooth muscle actin (αSMA) expression, revealing significant upregulation during matrix stiffening that remains transient and difficult to detect via traditional endpoint assays. Active stiffening also modulates fibroblast motility, transiently increasing migration speed while persistently enhancing directional persistence. Complementary computational reaction-diffusion modeling provides mechanistic insight into modulus gradient formation and reaction kinetics. This versatile toolbox enables investigation of early mechanobiological responses to matrix stiffening and may aid identification of markers of fibrotic disease onset.

## 1. Introduction

Extracellular matrix (ECM) properties are fundamental to tissue development and homeostasis, where physical and biochemical properties dictate tissue structure and function.^[1]^ In particular, ECM stiffness plays a critical role in cell differentiation, tissue development, and long-term homeostasis.^[2-3]^ Indeed, mechanical properties vary widely throughout the body, with elastic moduli spanning several orders of magnitude (Pa to GPa), reflecting the function of each tissue, where cells sense and respond to tissue-specific ECM stiffness.^[4]^ Through mechanotransduction, ECM modulus directs cell response, including differentiation and phenotype; proliferative and apoptotic pathways; migration; and ECM remodeling.^[5-6]^ Under homeostatic conditions, cells continually remodel the ECM to maintain tissue integrity over time. However, how cells respond to early ECM modifications associated with injury and maladaptive wound healing remains unclear.

Although tissue stiffness is generally stable in healthy adults, there are many circumstances that disrupt homeostasis and result in localized modulus increases. These changes arise from a cascade of matrix-remodeling processes, including enhanced ECM deposition, crosslinking, and fiber alignment; growth factor-driven cell stimulation (e.g., with TGFβ); and reduced ECM degradation due to changes in matrix metalloproteinase (MMP) and other ECM protease activity.^[7]^ Tissue stiffening, for instance, is essential to organogenesis and normal embryonic development.^[8-10]^ Wound healing similarly relies on matrix stiffening, from fibrin clot contraction during hemostasis to tissue stiffening at later stages, where the increase in modulus promotes a wound-healing macrophage phenotype.^[11-13]^ Matrix stiffening also accompanies the aging process, though this is associated with reduced mobility and increased risk of injury and disease; for example, advanced glycation end products (AGEs) accumulate over time and contribute to covalent crosslinking of ECM proteins.^[7, 14]^ While matrix stiffening can be part of normal physiological processes, aberrant or excessive matrix stiffening is characteristic of various pathological conditions, from maladaptive healing to chronic disease progression. For example, a controlled and temporary increase in matrix stiffness during acute inflammation promotes tissue regeneration, whereas prolonged inflammation can lead to significant matrix modifications and permanent tissue stiffening (e.g., in cancer and fibrosis)—though the ‘tipping point’ between these processes is not well understood.^[14-15]^ Consequently, improved model systems are needed to study the early transition from acute inflammation to disease toward providing mechanistic insights and establishing improved and relevant therapies.

Fibroblasts, mesenchymal cells highly sensitive to changes in matrix stiffness, are key regulators of local ECM composition and biological mechanisms.^[6, 16]^ For example, fibroblasts respond to tissue stiffness gradients by migrating from low to high modulus areas (durotaxis), where the increased stiffness indicates the need for fibroblast-mediated ECM regulation.^[17]^ Following an injury, fibroblasts are essential throughout all stages of tissue repair, migrating to the damaged tissue, secreting soluble factors to recruit various wound healing cells, and depositing new ECM proteins.^[18]^ In homeostatic environments, fibroblasts largely remain quiescent; however, matrix stiffness plays a critical role in regulating the fibroblast-to-myofibroblast transition through various mechanosensitive signaling pathways.^[19]^ Activated fibroblasts often exhibit alpha-smooth muscle actin (αSMA) within the cytosol (proto-myofibroblasts) and then αSMA rich stress fibers (mature myofibroblasts), a cytoskeletal structure characteristic of the myofibroblast phenotype and a critical contributor to the contractile forces required for wound closure and long-term tissue repair.^[20-22]^ While fibroblast activation is integral to normal healing, myofibroblast persistence in response to pathologically stiff ECM drives the cells to deposit additional matrix proteins, resulting in an uncontrolled feedback loop observed in fibrotic diseases such as idiopathic pulmonary fibrosis (IPF).^[15]^ Ongoing diagnostic research aims to elucidate the mechanisms governing these complex fibrotic diseases, which typically are detected only after significant disease progression, to develop more effective therapeutic interventions.^[23-24]^ Well-defined, three-dimensional (3D) culture models that capture aspects of matrix stiffening like that observed in actively fibrosing tissue will provide a valuable tool for probing fibrosis-induced modifications at the molecular and cell level.^[5]^

A variety of strategies have been established for triggering stiffening within hydrogel-based ECMs in two-dimensional (2D) and 3D cell culture with relevance for the design of developmental, injury, and disease models.^[25-28]^ After the formation of an initial hydrogel, stiffening has been achieved by modulation of physical interactions within the hydrogel polymer network. In one approach, responsive particles have been encapsulated within hydrogels to promote triggered physical interactions. For example, hydrogels containing acoustic-responsive particles^[29]^ or magnetic nanoparticles^[30-31]^ have undergone rapid, orders-of-magnitude increases in modulus over short timescales (order of minutes) when exposed to the appropriate stimulus. Other approaches have leveraged physical interactions within hydrogels to achieve reversible stiffening in response to salt^[32]^ or temperature^[33-34]^ with moderate modulus changes. These responsive materials typically require continued application of the stimulus to maintain the desired modulus (e.g., applied acoustic or magnetic field, maintained salt or temperature). Physical interactions have also been exploited by incorporating self-assembling motifs into hydrogels, where matrix stiffening is pre-programmed over the course of weeks upon the spontaneous formation of β-sheet domains, an approach with physiological relevance but limited user control.^[35-36]^

Complementary to these physical interaction approaches, hydrogel-based ECMs with unreacted functional handles have permitted later matrix stiffening through secondary covalent crosslinking events, where light is often a desirable stimulus for user-directed *in situ* control. For example, functional handles that undergo photodimerization upon the application of light without (e.g., anthracene^[37]^) or with photoinitiator (e.g., methacrylate,^[38-40]^ dibenzocyclooctyne^[41]^) have been used to achieve light-triggered stiffening of engineered ECMs. Similarly, harvested ECMs have been incubated with ruthenium and sodium persulfate for light-triggered formation of dityrosine bonds, crosslinking proteins within the matrix.^[42]^ Other photostiffening approaches have implemented heteromolecular reactions, where hydrogels are incubated with a combination of photoinitiator and monomer(s) followed by irradiation.^[43-44]^ Although these systems frequently achieve modulus increases representative of tissue stiffening from healthy to late-stage fibrotic tissue, many *in situ* photo-reactions and related stiffness changes take place on the order of seconds or minutes, while the initiation and progression of fibrosis takes place over days and weeks to years. To better match physiological timescales, spontaneous click chemistries have been successfully employed to induce matrix stiffening (e.g., tetrazine–TCO ligation, hydrazone click chemistry),^[45-46]^ where diffusion and reaction rates can be balanced to achieve significant increases in modulus over a more relevant timescale (order of days). Guided by mathematical modeling of diffusion and reaction rates, such bio-orthogonal covalent chemistries combined with physically assembled peptides provide an opportunity to address the need for 3D culture models that enable sequential, tunable stiffening over time for probing cellular responses from healthy tissue to early stages of fibrosis in collagen-rich microenvironments.

Here, we present a toolbox for dynamically studying cell behavior in actively stiffening environments. We employed a fully synthetic hydrogel-based ECM inspired by collagen-rich microenvironments that includes an available reactive handle for crosslinking at user-defined time points. Leveraging these orthogonal reactive handles, we established a unique stiffening approach utilizing a sequence of multistage, non-terminal reactions that achieve moderate modulus increases over physiologically relevant timescales. Specifically, the base synthetic ECM underwent a series of alternating incubations with two homo-functionalized multi-arm polymers, leveraging reaction kinetics, stoichiometry, and mass transfer to create a tunable stiffening system and decoupling the maximum achievable modulus from the initial hydrogel composition. As fibroblast activation and dysregulation are central to fibrotic disease progression, we evaluated this system in 3D fibroblast culture and examined how an actively stiffening matrix representative of the initiation of fibrosis impacts fibroblast behavior using a combination of endpoint assays and real-time measurements. While most studies of fibroblast mechanobiology have been conducted in 2D, recent works highlight important differences in how fibroblasts respond in 3D microenvironments, like those found in many tissues.^[19]^ As the full onset of fibrosis can be preceded by tissue stiffening,^[7]^ the system established here recapitulates the relatively small but significant modulus increases that occur in the early stages of fibrotic diseases and, with integration of reporter fibroblasts for dynamically probing cell activation, provides a platform for studying fibroblast responses during the initial phases of disease. To further understand how reaction-diffusion kinetics and the corresponding timescales influence cell response, we also developed a computational model to visualize the stiffening process. Together, this hydrogel system and its accompanying analytical tools are designed to support future efforts to identify key cellular markers associated with fibrotic disease onset, enabling improved diagnostic strategies and earlier therapeutic intervention.

## 2. Results and Discussion

### 2.1 Stiffening approach enables significant increases in hydrogel stiffness

We previously have designed and utilized multifunctional collagen-mimetic peptides (mfCMPs) that assemble into fibrils for covalent incorporation into a poly(ethylene glycol) (PEG) hydrogel network.^[34, 47-49^] This hybrid hydrogel system allows the incorporation of collagen type-I-like assembled structure via physical assembly of the mfCMP [e.g., K(azide)(PKG)_4_PK(alloc)G(POG)_6_(DOG)_4_] while imparting control over the initial mechanical and biochemical properties of the covalently crosslinked PEG-peptide synthetic matrix [tetrathiol PEG concentration, mono- and di-functional K(alloc) bioactive peptides], enabling independent tunability of initial matrix structure, modulus, and bioactivity.^[34]^ The specific sequence of the mfCMP facilitates triple helix formation through interpeptide hydrogen bonding and end-to-end peptide interactions for the formation of elongated fibrillar structures.^[50-51]^ The allyloxycarbonyl-protected lysine residue [K(alloc)] provides an alkene functional group for participation in a photoinitiated thiol–ene reaction for covalent integration into a larger hydrogel network,^[47]^ while the K(azide) has been used for visualization of the hierarchical structures within the resulting synthetic matrix.^[49]^ Given the success of this base material in 3D fibroblast cultures, with observations of motility and foci formation over days,^[34]^ in this new contribution we sought to leverage this azide functional handle for controlled hydrogel modification—specifically, hydrogel stiffening—and to dynamically probe fibroblast responses to the initiation of fibrotic conditions, leveraging fibroblasts that report on activation.^[52]^

For this work, base synthetic ECMs were formed using four-arm PEG tetrathiol (PEG-SH, 10 wt%; **Figure S1** and **S2**), and three peptides (**Figure 1a**): non-assembling linker peptide, integrin-binding pendent peptide, and pre-assembled mfCMP-Azide (**Figure S3-S5**). Hydrogel crosslinking was achieved through exposure to cytocompatible doses of long wavelength UV light in the presence of lithium phenyl-2,4,6-trimethylbenzoylphosphinate photoinitiator (LAP, 2.2 mM; **Figure S6**). To enable cell-mediated degradation and remodeling of the synthetic matrix, an MMP-degradable peptide was chosen for the non-assembling linker peptide; specifically, the linker peptide selected is inspired by collagen I and known to be sensitive to MMP-2, -9, and -14, all produced by fibroblasts.^[53-54]^ Further, an RGDS pendent peptide was included to promote cell-matrix interactions, as the RGDS peptide sequence is presented on various ECM proteins including vitronectin, fibronectin, and damaged or thermally unfolded collagen type-I, interacting with a number of integrin types displayed by human cells.^[55-58]^ RGDS was incorporated at 2 mM, a concentration found to consistently promote cell-matrix interactions within 2D and 3D synthetic hydrogels.^[48, 59-60]^ Importantly, the mfCMP-Azide peptide exhibited a melting temperature (*T*_m_, where 50% of the peptide is associated into triple helices and the remainder are individual peptide strands) of *T*_m_ = 37.5 ± 0.008 °C and consistent fibril formation (**Figure S5c**,**d**). Pre-assembled mfCMP-Azide was incorporated into the hydrogels at 5 mM (accounting for 28% of the peptide linkages in the formulation, with the remainder provided by the non-assembling peptide linker), where mfCMP concentrations as low as 2.5 mM (without azide) have been shown to promote elongated morphologies for human mesenchymal cell types.^[47]^ Consequently and importantly, the resulting initial hydrogel included 5 mM of free azide groups for later hydrogel modification.

**Figure 1.**
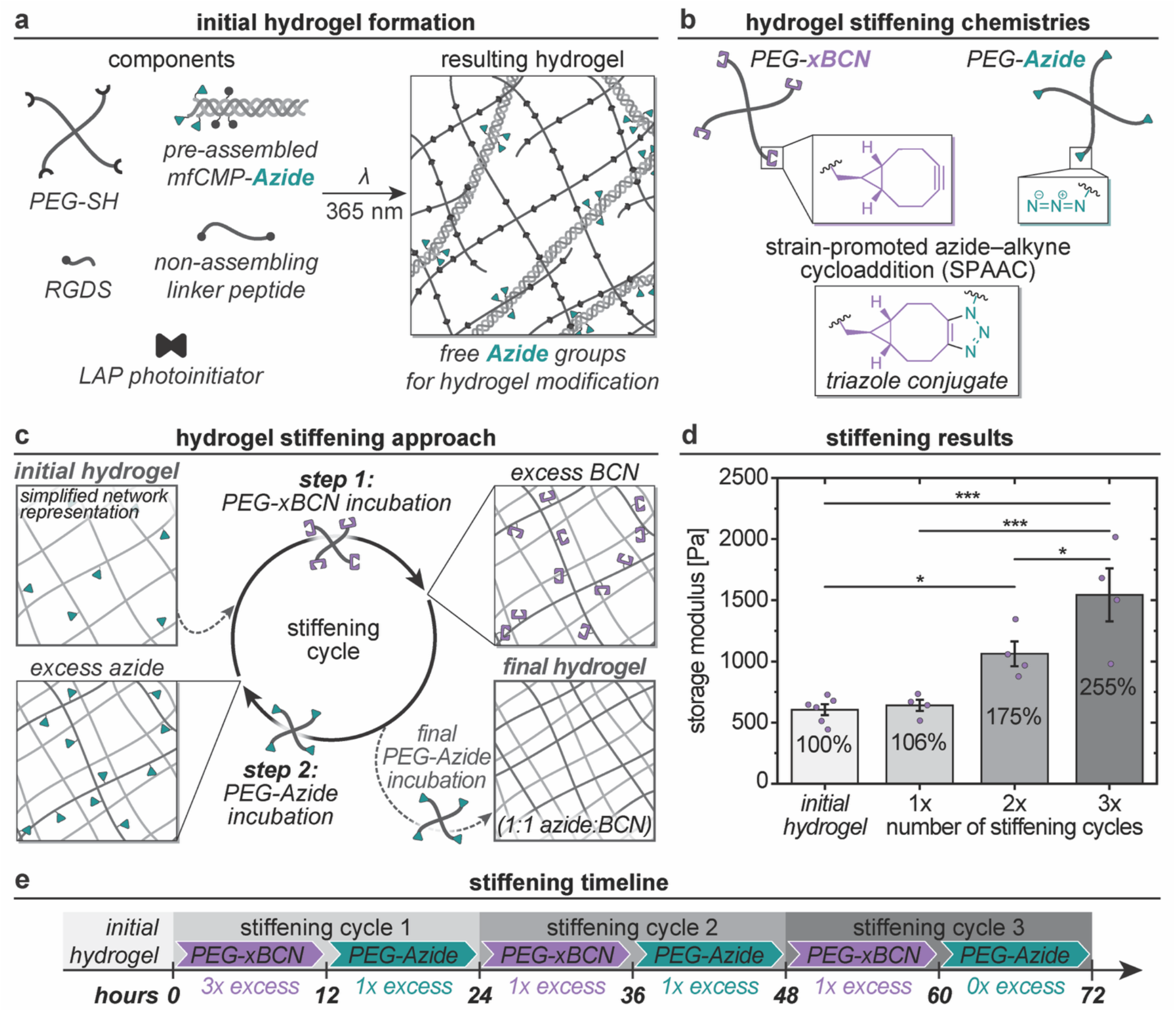
Hydrogel formation and stiffening approach. a) Schematic of initial hydrogel formation, where inclusion of pre-assembled mfCMP-Azide (along with PEG-SH, non-assembling linker peptide, RGDS pendent peptide, and LAP photoinitiator) results in a hydrogel with a 1:1 ratio of thiol:alloc with free azide groups available for later modification. (Schematics not to scale; hydrogel component concentrations available in **Table S1**.) b) Macromolecular components used for hydrogel stiffening: a 4-arm PEG-xBCN and a 4-arm PEG-Azide allowed for the implementation of SPAAC chemistry (spontaneous and cytocompatible). c) Simplified schematic demonstrating the hydrogel stiffening approach, beginning with the initial hydrogel (top left). In step 1, the hydrogel (free azide groups) is incubated in a solution of excess PEG-xBCN such that it reacts with the free azides in the network as it diffuses in, resulting in a hydrogel with dangling free BCN groups. Next, in step 2, the hydrogel (now with free BCN groups) is incubated in a solution of excess PEG-Azide for reaction with the free BCN as it diffuses in, resulting once again in a hydrogel with free azide groups. This cycle may be repeated multiple times, ending with a final PEG-Azide incubation (in place of step 2) to bring the total azide:BCN to a 1:1 ratio and eliminate any remaining functional groups. d) Oscillatory rheometry was performed on initial hydrogels after equilibrium swelling and hydrogels after 1, 2, or 3 cycles of stiffening, where the percentages represent the storage modulus after each cycle relative to the initial hydrogel. All incubations and measurements were conducted at 37 °C. (Mean ± SE with individual datapoints; n ≥ 4 independent samples for each condition; Significance determined via one-way ANOVA followed by Tukey’s post-hoc test: * p < 0.05, ** p < 0.01, *** p < 0.001.) e) Timeline of the stiffening cycles used throughout this study, including the excess amounts of stiffening polymer used for each step.

Toward inducing hydrogel stiffening, the free azide groups available throughout the initial hydrogel were leveraged for secondary click chemistry reactions. While we have successfully implemented copper-catalyzed azide–alkyne cycloaddition (CuAAC) click chemistry conjugations with the free azide groups using a similar hydrogel composition,^[49]^ the CuAAC reaction is inherently cytotoxic, and thus not amenable to application in cell culture conditions.^[61-62]^ As a spontaneous and cytocompatible alternative, we utilized the strain-promoted azide–alkyne cycloaddition (SPAAC) conjugation reaction to induce stiffening.^[62-64]^ Specifically, a four-arm PEG tetra-bicyclononyne (*exo*) (PEG-xBCN) and a four-arm PEG tetra-azide (PEG-Azide) were used, where the BCN functional group was elected over one of the bulkier alkyne groups available (**Figure 1b, Equation S1-S4**).^[62]^ Stiffening then was achieved by alternately diffusing PEG-xBCN and PEG-Azide into the hydrogel for reaction with the free functional groups (**Figure 1c, Figure S7** and **S8**). Specifically, each stiffening cycle consisted of a PEG-xBCN incubation followed by a PEG-Azide incubation, each for approximately 12 hours. Importantly, the stiffening solution concentrations (specifically, PEG functional group concentration) were calibrated according to the hydrogel equilibrium swollen volume and resulting change in available functional groups (i.e., 20 μL, 5 mM azide at preparation; 130 μL, 0.77 mM azide after swelling), as well as the incubation solution volume (250 μL) compared to the total volume (including the 130 μL hydrogel = 380 μL) (**Table S2**). The stiffening process was commenced with an initial ‘high concentration’ PEG-xBCN incubation (3× molar excess BCN relative to 0.77 mM azide), and all subsequent stiffening incubations were conducted at the ‘standard concentration’ (PEG-xBCN or PEG-Azide; 1× molar excess relative to the free azide or BCN functional groups, respectively). A final PEG-Azide incubation (1× molar equivalent relative to free BCN groups) was implemented, resulting in a 1:1 azide:BCN ratio, thus terminating the stiffening process. This process was designed such that each stiffening cycle (PEG-xBCN incubation followed by PEG-Azide incubation), may be repeated any number of times, according to the desired change in modulus, where each cycle results in an increased crosslink density within the hydrogel.

In this work, as a demonstration of this stiffening technique, we explored how three stiffening cycles over the course of 72 hours impacts hydrogel mechanical properties (**Figure 1d**,**e**). Toward understanding the changes in mechanical properties experienced by cells *in vitro* throughout the stiffening process, all swelling and stiffening incubations were conducted under cell culture conditions (37 °C, 5% CO_2_, with humidity), with rheometric measurements performed at 37 °C (ambient CO_2_, ambient humidity). The initial hydrogel (measured immediately before the first stiffening cycle) was found to have a shear storage modulus of *G′* = 0.605 ± 0.043 kPa, which equates to a Young’s modulus of *E* = 1.82 ± 0.131 kPa based on rubber elasticity theory, within the range of normal lung tissue (0.5-3 kPa).^[16, 65]^ Hydrogels underwent 1, 2, or 3 cycles of stiffening prior to measurement, and over the course of 3 stiffening cycles, hydrogels reached an average storage modulus greater than 2.5× that of the initial hydrogel modulus, demonstrating the efficacy of this stiffening approach. Specifically, after completing the full 3 stiffening cycles, the modulus increased to *G′* = 1.55 ± 0.217 kPa (*E* = 4.64 ± 0.650 kPa, higher than healthy lung tissue and below the 10-to 50-fold increase observed in late-stage fibrotic tissue^[65]^), providing a model system to study relatively small but significant increases in tissue stiffness during early stages of fibrotic lung disease. Further, conducting thorough washes to remove any remaining monomers after stiffening cycle 3 did not result in any significant changes to the modulus (**Figure S9**).

The significant impact that this cyclical SPAAC stiffening technique has on modulus over time establishes this approach as an effective method for cytocompatible stiffening of synthetic matrices. While the initial hydrogel-based synthetic ECM mechanical properties and stiffening characteristics explored here are relevant for modeling early stages of fibrotic lung disease, the general approach allows for a high degree of tunability independent from the initial hydrogel composition. The rate of stiffening within each cycle and overall stiffening timescale may be easily adapted for different applications by modifying polymer concentration or functionality, adjusting the number of cycles, or by spreading the stiffening cycles over a longer period (i.e., include incubations between stiffening cycles rather than conducting all cycles back-to-back, given the bio-orthogonal chemistry). With the tunability afforded by this stiffening approach, this system has the potential to be tailored to different tissues and model applications. For example, the initial hydrogel components may be adjusted to capture the modulus of various tissues: wound healing (including healthy and aberrant) could be modeled with a softer initial hydrogel (to capture properties of the fibrin-rich ECM that immediately develops at the injury site) using a short stiffening timescale with large modulus increases, and various disease progression models could be developed using longer stiffening timescales and the relevant degree of stiffening.^[16, 66]^

### 2.2 Macrophage-conditioned media influences fibroblast metabolic activity, while hydrogel stiffening does not

Importantly, changes in ECM stiffness direct cell response, including alterations in gene expression and protein function, determination of cell fate (e.g., activation of fibroblasts into myofibroblasts, stem cell differentiation lineage), variations in metabolic activity, and increased apoptosis resistance.^[65, 67-68]^ Therefore, studying if, and how, the stiffening model developed here directs cell response is critical. Further, while ECM mechanical properties certainly impact cell behavior and play a role in tissue remodeling, biochemical factors, too, significantly influence cell response. In the context of matrix remodeling, signaling between macrophages and fibroblasts is particularly consequential in activating myofibroblasts, from healthy wound healing to fibrosis associated with aberrant healing and chronic disease progression.^[16, 68]^ Specifically, macrophages secrete a variety of proteins involved in tissue inflammation (e.g., TGF-β, IL-6, PDGF) and promote collagen deposition and fibrotic remodeling.^[66]^ We therefore proceeded to study the effect of matrix stiffening or a biochemical stimulus, specifically, macrophage-conditioned media (CM), on cell response.

Toward assessing the relative influence of early-stage stiffening or biochemical signaling on cell response, human lung fibroblasts (CCL-151) were encapsulated within the initial hydrogel composition at a density of 5 × 10^6^ cells mL^-1^. Fibroblasts were cultured in hydrogels for 7 days to allow the cells to become established in the matrix (days -9 – -2 of the experiment), at which point the fibroblasts were weaned from 10% fetal bovine serum (FBS) media to 1% FBS media over the course of 48 hours (days -2–0) (**Figure 2a**). The stimuli were then introduced for each condition (0 hours) and maintained over the course of the experiment: (*i*) control (cultured in 1% FBS media), (*ii*) CM, or (*iii*) stiffened. Hydrogels treated with CM were cultured with 100% CM for the first ∼24 hours and 50% CM (with 1% FBS media) for the remainder of the experiment. Separately, hydrogels undergoing stiffening were subjected to 3 stiffening cycles over the course of ∼3 days then maintained for the duration of the experiment (Cycle 1: 0-26 hours; Cycle 2: 26-51 hours; Cycle 3: 51-76 hours; continued culture in 1% FBS media: 76-120 hours).

**Figure 2.**
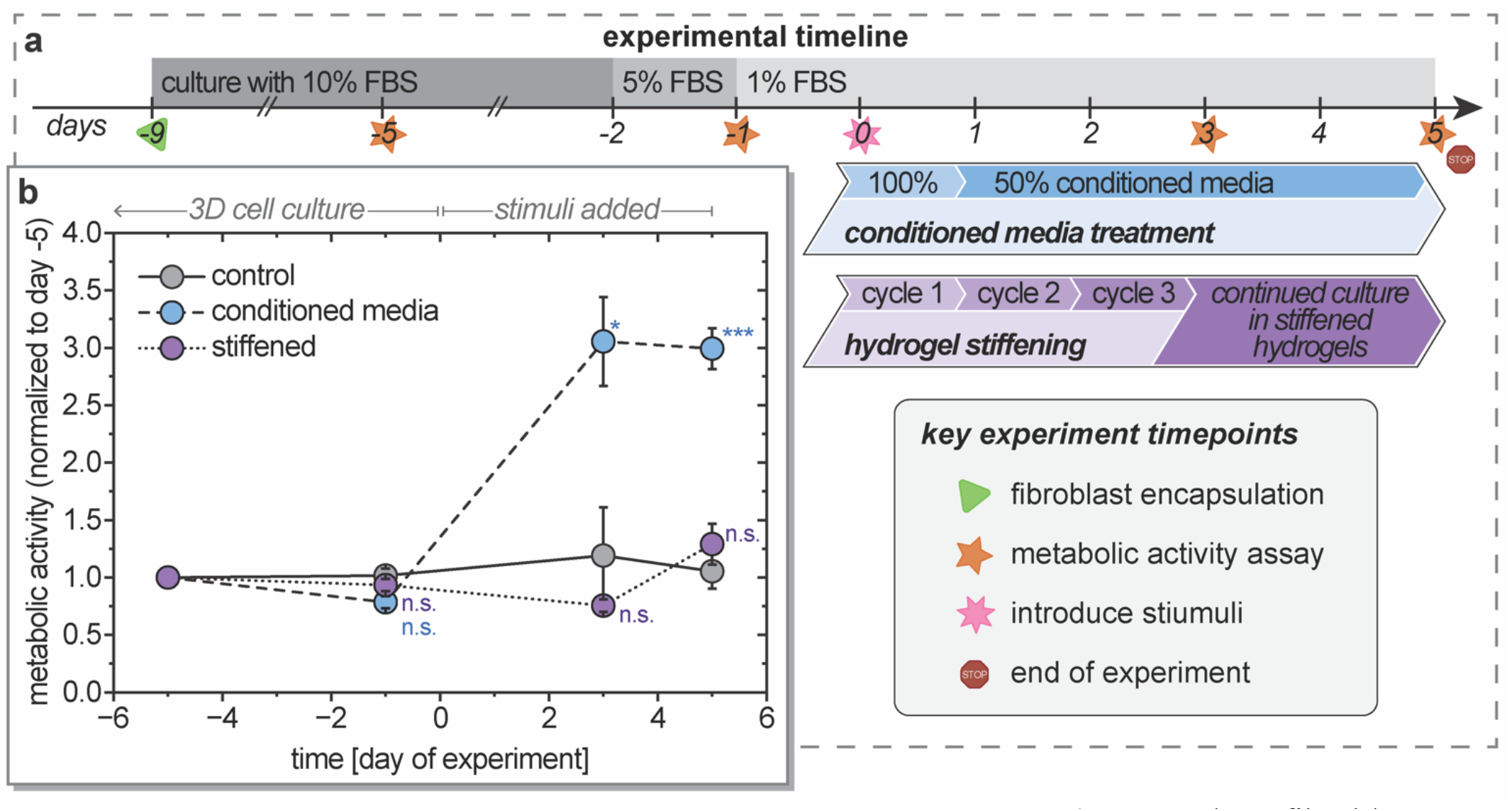
3D cell culture experimental timeline and metabolic activity. a) Human lung fibroblasts were encapsulated during the initial hydrogel formation, cultured for 7 days to acclimate to the 3D culture system, and weaned from culture media supplemented with 10% FBS down to 1% FBS. Experimental stimuli then were introduced: encapsulated cells were either treated with macrophage-conditioned media (CM) for the remainder of the experiment (blue, dashed line) or subjected to hydrogel stiffening over the course of 3 days and cultured in the stiffened hydrogels for the remainder of the experiment (purple, dotted line), in comparison to an untreated control group (gray, solid line). b) Metabolic activity was assessed throughout the experiment both before (days -9 to 0) and after (days 0 to 5) implementation of experimental stimuli, where each hydrogel was internally normalized to its day -5 metabolic activity. Metabolic activity data with complete multiple comparison analyses and individual data points are available in **Figure S10** for each timepoint; statistical analyses within conditions over time are available in **Table S3.** (Mean ± SE; n = 3 independent samples for each condition; Significance determined via one-way ANOVA followed by Tukey’s post-hoc test: * indicates a significant difference from the control at the indicated timepoint, where * p < 0.05, ** p < 0.01, *** p < 0.001, and n.s. indicates that the means are not significantly different.)

Metabolic activity was monitored over the course of the experiment as an assessment of cell health and general level of metabolism in response to stimuli (**Figure 2b**). Prior to the introduction of stimuli at day 0, there were no deviations in metabolic activity from the control: an expected result as all conditions were undergoing the same culture conditions over that time. Importantly, the stiffening treatment did not have a negative impact on metabolic activity, as there was no significant difference from the control at timepoints after inducing mechanical changes. Conversely, the metabolic activity for cells cultured with CM was significantly higher than both the control and stiffened conditions for all timepoints after the introduction of stimuli (**Figure S10**). This increase in metabolic activity was not due to an increase in cell number, as there were no differences in cell density between conditions at day 5 (**Figure S11**), but rather an increase in cell-level metabolic activity. This increase in fibroblast metabolic activity in response to CM is consistent with previous observations that TGF-β leads to increased metabolic activity, specifically glycolysis and glutaminolysis, for increased collagen production (a main driver of ECM stiffening),^[69]^ and that general changes in fibroblast metabolic activity are associated with macrophage-mediated inflammation.^[66]^

### 2.3 Endpoint immunostaining suggests no significant changes in αSMA expression or fiber organization

Fibroblast activation to a myofibroblast phenotype is critical for proper wound closure and general tissue remodeling and is characterized in part by the production and organization of αSMA, a profibrotic gene product.^[16, 70-72]^ In addition, activation can result in morphological changes from spindle shaped cells (fibroblasts) to a more stellate and elongated morphology (myofibroblasts) with increased contractility and reduced migration.^[73-74]^ Toward characterizing myofibroblast activation response, cells were immunostained for nuclei (Hoechst), F-Actin, and αSMA (**Figure 3**). *z*-stack images were collected with confocal microscopy and were analyzed as either 3D renderings or orthogonal projections, where all analyses were based on individual cell objects (either a single cell or cluster of cells in contact with each other) or total image fluorescence.

**Figure 3.**
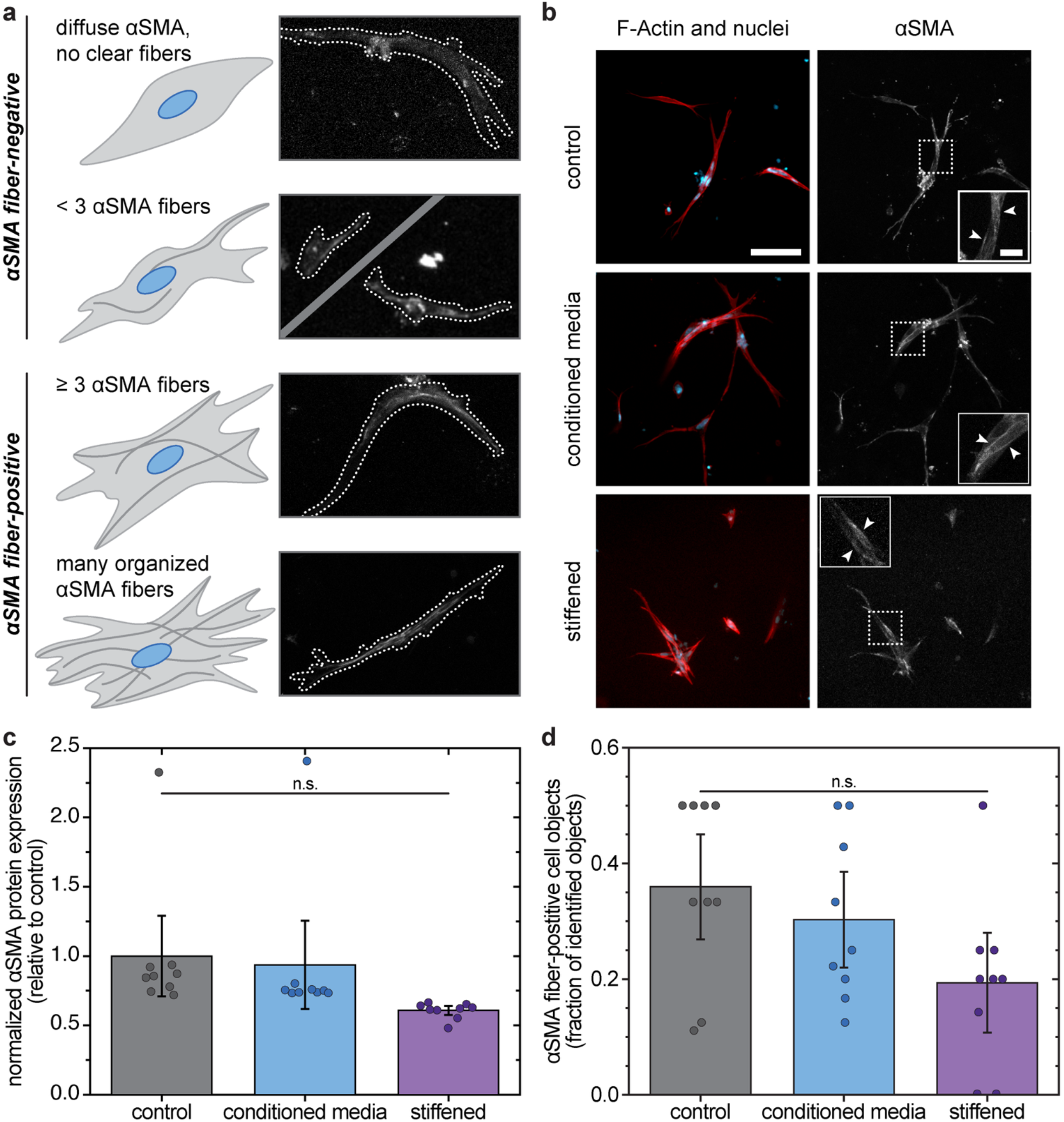
Impact of biochemical or mechanical stimuli on αSMA protein expression and organization in human lung fibroblasts. a) Schematic depicting different levels of αSMA organization (left), along with example images of fibroblasts immunostained for αSMA with cell body outlines indicated (right). Cell objects were categorized as negative for αSMA fibers if fewer than 3 individual fibers were detected and positive if 3 or more fibers were detected. b) Orthogonal projections of confocal *z*-stacks of fixed and immunostained fibroblasts in 3D culture (14 days after encapsulation, 5 days after introducing experimental stimuli; scale bar = 100 μm; inset image scale bar: 25 μm). Cells were stained for F-Actin and nuclei (left; red and blue, respectively) as well as αSMA (right). Inset images more clearly depict examples of organized αSMA fibers, indicated by white arrows. c) Endpoint quantification of total αSMA protein expression relative to the control. (Mean ± SE with individual datapoints each representing the measurement from a single image; n = 3 independent samples for each condition, with measurements from ≥40 cell objects per condition; Significance determined via one-way ANOVA followed by Tukey’s post-hoc test: n.s. indicates that the means are not significantly different. Additional analyses available in **Figure S12**.) d) Quantification of cell objects positive for visually identifiable αSMA fibers as a fraction of the total population of identified cell objects.

Toward analyzing αSMA expression and organization at the protein level, αSMA immunostaining images were quantified to determine total fluorescence and whether each cell object contained αSMA-positive stress fibers (**Figure 3b**). For each 3D image, the total αSMA fluorescence was normalized to the total Hoechst fluorescence, and no significant differences were found between conditions (**Figure 3c**). Similarly, quantification of αSMA fiber-positive cell objects resulted in no significant differences between any of the conditions (**Figure 3d**). For the purposes of this analysis, diffuse αSMA staining with no clear fibers and cell objects with fewer than three αSMA-associated fibers were categorized as negative for αSMA-positive stress fibers (with some cells negative for any F-Actin stress fibers), while cell objects with ≥ 3 αSMA-associated fibers were considered positive for αSMA-positive stress fibers.

Qualitatively, fibroblasts in the control and CM conditions appeared more spindle-like, whereas some cell objects in the stiffened condition appeared to adopt more stellate morphologies. These observations were quantified using the 3D shape factor (a measurement of sphericity) to detect any differences in cell morphology between conditions. Three-dimensional renderings of the collected *z*-stacks were analyzed to determine the shape factor of each cell object. When comparing only the control and stiffened conditions, the overall shape factor of cells after stiffening was significantly lower, indicating greater deviation from a spherical morphology, than that of cells in the control (p = 0.021; direct comparison with Student’s t-test) (**Figure S13**). While cell morphologies in the stiffened condition trended toward less spherical, there was not a significant difference in overall shape factor amongst the conditions when analyzed together with an ANOVA given the variance amongst conditions.

While useful, these endpoint immunostaining methods fail to capture the real-time impact of treatments, lending only a narrow insight into cell response. Further, although immunostaining for αSMA and quantification of αSMA-positive stress fibers is commonly used to identify myofibroblastic phenotypes in 2D culture, this approach yields limited success in 3D systems, as αSMA-positive stress fibers are more difficult to observe in 3D culture.^[75]^ Based on the metabolic activity and morphology results, the cells are exhibiting some response, yet the dynamics of the cellular responses may not be well captured by implementing only endpoint assays. An alternative approach for monitoring fibroblast activation in 3D matrices is therefore necessary. One technique implements lentivirus-transduced reporter cells and has enabled real-time monitoring of αSMA expression in human mesenchymal stem cells and fibroblasts.^[52, 76]^ This approach has been applied to cells in 2D culture with successful benchmarking versus αSMA expression by immunostaining, suggesting opportunities for use in 3D culture applications.

### 2.4 Live cell reporter indicates an increase in αSMA expression during active stiffening

Toward understanding the real-time fibroblast response to stimuli treatment, we monitored αSMA expression in live 3D cell culture as lentivirus-transduced reporter cells (human lung fibroblasts, CCL-151) were exposed to ECM stiffening or CM. These cells (*i*) constitutively express a red fluorescent protein, DsRed-Express2 (DsRed) and (*ii*) conditionally express a green fluorescent protein, ZsGreen, under the promotor for the profibrotic gene *ACTA2*, which encodes the αSMA protein.^[52]^ Notably, the *ACTA2* gene is not normally expressed in non-activated fibroblasts but is a myofibroblast-associated gene.^[16]^.

Confocal *z*-stack images of live cells were captured over the course of the experiment (2, 15, 37, and 65 hours after introducing stimuli), while fibroblasts were actively undergoing stiffening or CM treatment (**Figure 4a**). After creating sum intensity orthogonal projections to preserve total fluorescence, cell objects were detected using the combined DsRed and ZsGreen channels, and the total fluorescence intensity of each channel was determined for each identified cell object (**Figure 4b**). The normalized αSMA intensity of each cell object was calculated by normalizing the variable green fluorescence intensity to the constitutive red fluorescence intensity (ZsGreen:DsRed ratio). Due to the abundance of direct cell-cell interactions, we were unable to separate cell objects into individual cells, and the normalized αSMA intensity for each cell object was weighted according to the area of the corresponding object to account for the variations in object size.

**Figure 4.**
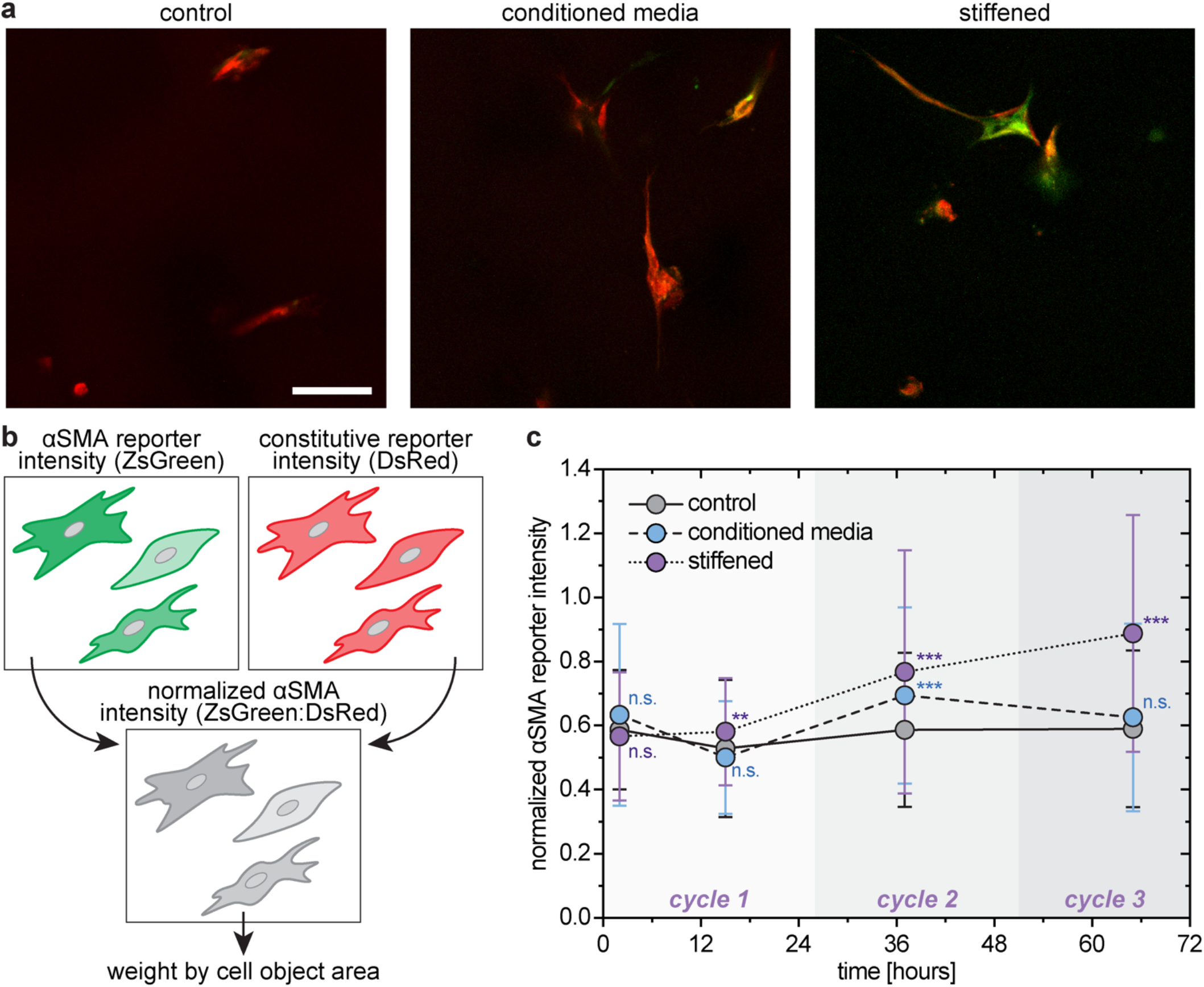
Real-time impact of active biochemical treatment or environmental stiffening on αSMA expression in human lung fibroblasts. a) Representative live reporter images (orthogonal projections of confocal *z*-stacks; 37 hours after introduction of stimuli; scale bar = 100 μm). DsRed constitutive reporter (red), ZsGreen αSMA reporter (green), and overlap (orange-yellow). b) Schematic representing the confocal imaging detection of the 2 reporter proteins (conditionally expressed ZsGreen that is produced when αSMA expression is upregulated and constitutively expressed DsRed), followed by normalization of the αSMA reporter (e.g., ZsGreen:DsRed) within each cell object. The normalized αSMA reporter intensity was then weighted by the cell object area to account for disparities between single cell objects and cell cluster objects. c) Normalized αSMA reporter intensity was initially quantified for each condition immediately after introducing the experimental stimuli (*t* = 2 hours). All remaining timepoints were chosen near the middle of each stiffening cycle (for all conditions). Example reporter images, full data (half violin plots with individual data points), and complete multiple comparison analyses are available in **Figure S14** and **S15** for each timepoint; statistical analyses within conditions over time are available in **Table S4.** (Mean ± SE; n = 3 independent samples for each condition, with measurements from ≥48 cell objects per condition at each timepoint; Significance determined via one-way ANOVA followed by Tukey’s post-hoc test: * indicates a significant difference from the control at the indicated timepoint, where * p < 0.05, ** p < 0.01, *** p < 0.001, and n.s. indicates that the means are not significantly different.)

Significant differences between conditions were observed with the real-time monitoring of αSMA expression (**Figure 4c**), allowing observations of the dynamics of cellular responses not provided by endpoint assays. At the earliest timepoint (2 hours after introducing the stimuli), the normalized αSMA reporter intensity of each treated condition was statistically the same as the control. The lack of differences at such an early timepoint was expected as both the CM and stiffening treatments are diffusion limited and changes in αSMA expression become apparent on the order of hours after application of a stimulus.^[77-78]^ Notably, by 15 hours, a slight but significant increase in αSMA expression emerged in fibroblasts subjected to hydrogel stiffening relative to control. αSMA expression continued to increase over the course of the 3 stiffening cycles with differences of increasing significance compared to the control. Interestingly, treatment with CM did not appear to have as notable or prolonged an effect on αSMA expression. Fibroblasts treated with CM exhibited a temporary increase in αSMA relative to control at 37 hours but returned to levels within the control baseline by 65 hours. The differences observed between the real-time and endpoint αSMA expression could further indicate transient fibroblast activation like that observed during wound healing and tissue remodeling events, which is followed by a persistently activated phenotype in fibrotic environments.^[79]^

These live cell αSMA expression results demonstrate that real-time monitoring of cell response is crucial to understanding how cells respond to changes in their environment. While fixed cell immunostaining offers valuable insight into a wide range of cell responses and mechanisms, in this instance hydrogel stiffening directed cell response beyond what we were able to detect at a single final timepoint. Active hydrogel stiffening appeared to have a significant and prolonged impact on fibroblast response, where significant increases in αSMA expression indicate a shift toward activated myoblast phenotypes. In comparison, the significant but temporary increase in αSMA expression in response to CM treatment may capture a cycle of fibroblast activation and reversion, as transient αSMA expression is associated with non-fibrotic healing.^[80]^ Ultimately, the stiffening hydrogel system presented in this work significantly impacts the expression of fibroblast activation marker αSMA, indicating that this stiffening approach has utility as a model ECM to study various cell responses to a stiffening environment. This is of particular interest as early diagnoses in lung diseases such as IPF are crucial to improving patient prognosis and treatment efficacy; however, overlapping symptoms often result in late or inaccurate diagnoses.^[23, 81]^ The use of biomarkers for improving the speed and accuracy of lung disease diagnoses is of increasing interest, and the stiffening system presented here allows for future exploration of biomarkers associated with a particular fibrotic disease to enable early detection and precision medicine.^[82]^

### 2.5 Fibroblast motility is impacted by dynamic stiffening or biochemical cues

As fibroblasts play a critical role in healthy tissue healing, from late inflammatory to final remodeling stages, initial fibroblast migration to the injury site is essential in the transition from the inflammatory phase to the proliferative phase.^[18]^ Accordingly, rapid fibroblast migration to injured tissue is crucial for wound repair: decreased cell migration often leads to fibrotic healing such as scarring, while higher rates of cell migration facilitate non-fibrotic ECM remodeling and correlate with lower αSMA expression.^[21]^ Further, fibroblast migration contributes to fibrotic disease progression through the process of durotaxis (i.e., migration from soft to stiff ECM), which can create a pro-fibrotic feedback loop: (*i*) fibroblasts migrate to fibrotic areas; (*ii*) fibroblasts activate in response to increased ECM stiffness; and (*iii*) myofibroblasts further increase ECM deposition, further increasing tissue stiffness.^[83]^ Given this context, we were interested in studying how fibroblast motility would be impacted within this 3D culture system when subjected either to a stiffening environment or CM treatment.

Utilizing the lentivirus-transduced reporter fibroblasts, we were able to track the cells during live 3D imaging with the constitutive DsRed reporter protein. Timelapses were captured over the course of the experiment while fibroblasts were treated with either a stiffening environment or CM (timelapses: 2-12, 15-24, 37-45, and 65-74 hours). Within the 3D rendering of each timelapse, cell objects were identified and tracked for the duration of the timelapse (or for the period the object was in frame), and hydrogel drift was tracked using brightfield *z*-stacks such that background movement could be subtracted from the raw cell object positions. This information was used to generate 3D track plots for each object to visualize cell movement (**Figure 5a**), namely the total distance traveled by the object (*D*, sum of distances traveled between each frame) and the displacement (*d*, distance between the initial and final locations for each object).

**Figure 5.**
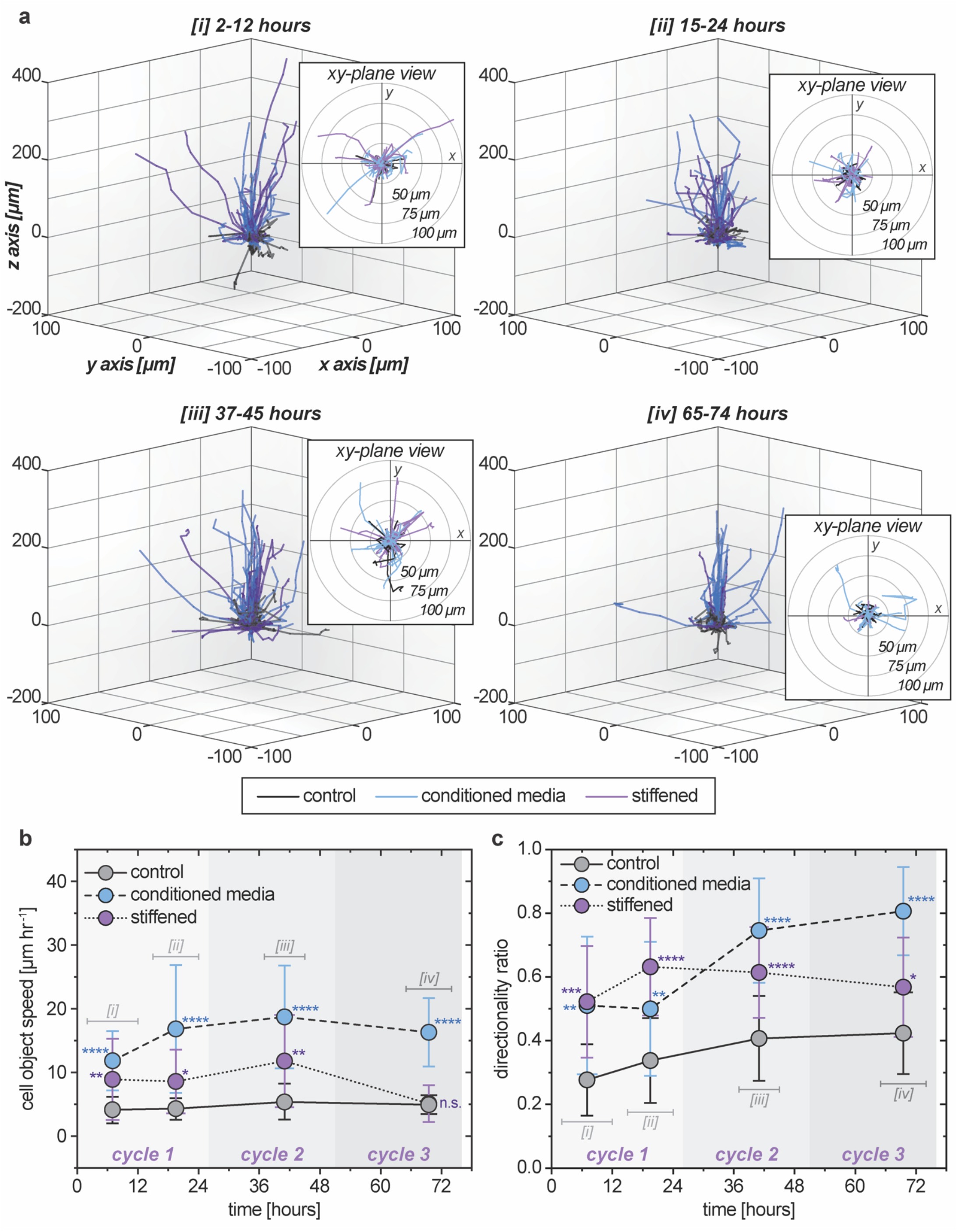
Impact of active biochemical treatment or matrix stiffening on human lung fibroblast cell motility in three dimensions. a) 3D traces of cell tracks over the indicated time lapse from beginning of the application of experimental stimuli, with insets of the *xy*-plane view (2D top-down). Each identified cell object (individual cell or cluster of multiple cells) was tracked in three dimensions over the indicated time frame to determine measurements of motility. Cell tracks separated by condition are available in **Figure S16 and S17.** b) Cell object speed was tracked over 4 different timelapses, indicated on the plot and labeled [*i*]-[*iv*]. (Mean ± SE; n = 3 independent samples for each condition, measurements from ≥47 ([*i*]), ≥52 ([*ii*]), ≥51 ([*iii*]), or ≥36 ([*iv*]) cell objects per condition; Significance determined via one-way ANOVA followed by Tukey’s post-hoc test: * indicates a significant difference from the control at the indicated timepoint, where * p < 0.05, ** p < 0.01, *** p < 0.001, **** p < 0.0001, and n.s. indicates that the means are not significantly different.) c) The directionality ratio of each cell object was tracked over the same 4 timelapses. Values closer to 0 are indicative of meandering cell movement (low directional persistence), whereas values closer to 1 are indicative of more directional cell movement (high directional persistence). Full data (half violin plots with individual data points) and complete multiple comparison analyses are available in **Figure S18** and **S19** for each timepoint; statistical analyses within conditions over time are available in **Table S5** and **S6**.

Cell motility was quantified by calculating the speed and directionality ratio for each track (**Figure 5b**,**c**). Speed was calculated by the distance traveled (*D*) divided by the track time, and the directionality ratio was calculated by the displacement (*d*) divided by *D*. Fibroblasts cultured in the control hydrogel maintained a consistent speed over the course of the experiment and began moving in a slightly more directional manner at late timepoints, perhaps due to cell-mediated matrix degradation that allowed for less inhibited local movement along cell tracks and/or paracrine signaling between cells. In comparison, fibroblasts cultured in a hydrogel undergoing active stiffening exhibited significantly faster motility over the first 48 hours and then slowed as stiffening progressed in Cycle 3, reducing to speeds comparable to that of fibroblasts in the control. Notably, the fibroblasts moved with significantly more directionality than the control throughout the experiment. Fibroblasts treated with CM maintained a fairly consistent speed over the course of treatment, where they initially moved significantly faster than those in the control condition but not the stiffened hydrogels. However, by the 15–24-hour timepoint, cells exposed to CM moved significantly faster than the fibroblasts in both the control and stiffened conditions, which continued for the remainder of the experiment. Further, while the fibroblasts treated with CM maintained significantly higher directional persistence than the control throughout the experiment, the fibroblasts in CM exhibited a significant increase in directionality between the second and third timepoints.

Interestingly, both experimental conditions exhibited a high directionality ratio relative to the control, a migration result we hypothesize is in response to the respective gradient. Both cases featured a ‘solute’ diffusing into the hydrogel from the surrounding solution: (*i*) soluble factors associated with fibroblast signaling (CM) or (*ii*) polymer that reacted with the hydrogel and theoretically resulting in a modulus gradient as the polymer moved toward the center of the hydrogel (stiffened). In response to the CM, fibroblasts were likely migrating toward higher concentrations of soluble biochemical factors, analogous to fibroblast recruitment by macrophages in early stages of tissue healing and remodeling. With respect to cell responses to the stiffening condition, several factors may play a role in the response differences initially and over time. Given sufficient cell-matrix interactions (relative to cell-cell interactions), fibroblasts within a matrix are known to undergo durotaxis, and cells likely sense matrix tensions over long distances (relative to the length scale of the cell) to perceive such modulus gradients.^[84-85]^ In this context, the temporary increase in fibroblast speed (through stiffening cycle 2) and sustained high directional persistence in the stiffened hydrogels (compared to the control) may be a result of durotaxis, especially given that fibroblasts in the stiffened condition appear to be primarily migrating upward toward the source of the stiffening polymer, and consequently in the direction of increased matrix stiffness. Further, cells in 3D culture are known to migrate more slowly in stiff matrices compared to soft matrices;^[86]^ additionally, human lung fibroblasts have been shown to move more slowly on fibrotic lung tissue than normal lung tissue in 2D.^[87]^ In this context, the increased hydrogel stiffness over time may slow down migration speed, which is observed to be similar to the control by 72 hours, without impacting the directionality of the fibroblasts. The initial peak in cell speed followed by a significant taper also could be attributed to and influenced by the transient nature of signaling during fibroblast activation in response to a stimulus: for example, fibroblasts activated with TGF-β1 have demonstrated high migration potential at early timepoints, which is significantly reduced over time as αSMA expression increases.^[88]^ Ultimately, the stiffening model developed here promotes an increase in the fibroblast activation marker αSMA and an apparent change in phenotype from migrating fibroblasts to contractile myofibroblasts.

### 2.6 Mathematical models enable additional insight into stimuli impact on fibroblast response

Pairing experimental data with computational models can provide insights into experimental results and phenomena, enable outcome predictions, and assist in experimental design. For example, computational models have been used to better understand time-dependent processes within hydrogels (including swelling behavior, stress distribution, and changes in crosslink density)^[89-90]^ and gain insight into diffusion-limited reaction kinetics.^[91]^ In this work, we aimed to create a mathematical model for understanding the dynamics of microenvironment changes within the 3D culture system by describing the reaction-diffusion kinetics of biochemical or mechanical stimuli for insights into the timescales of these processes and improving our understanding of associated fibroblast responses. Specifically, our goal was to model a solute diffusing into the 3D culture construct (a non-semi-infinite cylinder) while undergoing degradation (in the case of CM) or reaction with the hydrogel network (in the case of functionalized PEG).

#### 2.6.1 Problem formulation

In this model, the hydrogel and surrounding solution represent a 2D axisymmetric, closed system held at a constant temperature (**Figure 6a**), such that the relevant flux equations become:

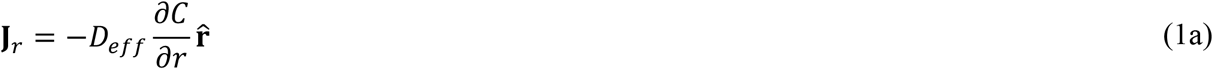

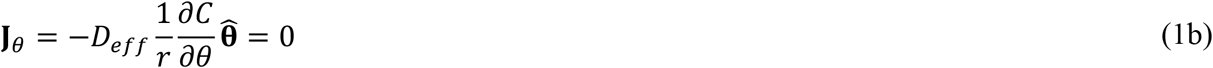

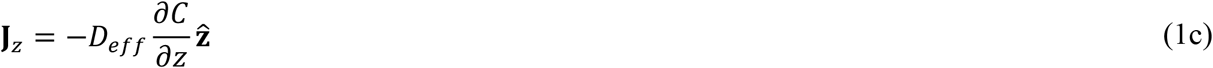

where *Deff* is the solute effective diffusivity within the hydrogel and *C* is the molar concentration of the solute within the hydrogel. The hydrogel is a vertical cylinder of radius R and height *H*, assumed to be stationary, and with the bottom of the hydrogel in contact with the well bottom, creating a no-flux boundary condition. Surrounding the hydrogel is a finite volume of solution (incompressible water-based fluid) carrying a solute (CM or functionalized PEG, i.e., single species diffusion). The solute concentration in solution (*C*_∞_) is assumed to be at equilibrium without the effects of convection. Further, *C*_∞_ is treated as a constant for the purpose of solving the system. Importantly, a constant *C*_∞_ is not a valid representation of the system, as the hydrogel and solution represent a closed system; rather, as the solute (protein or polymer) diffuses into the hydrogel, *C*_∞_ will decrease. This inaccuracy was later accounted for in the computational model by calculating molar flux into the hydrogel and updating *C*_∞_ at discrete timepoints, changing the representation of *C*_∞_ from a physically impossible constant to a more realistic step function.

**Figure 6.**
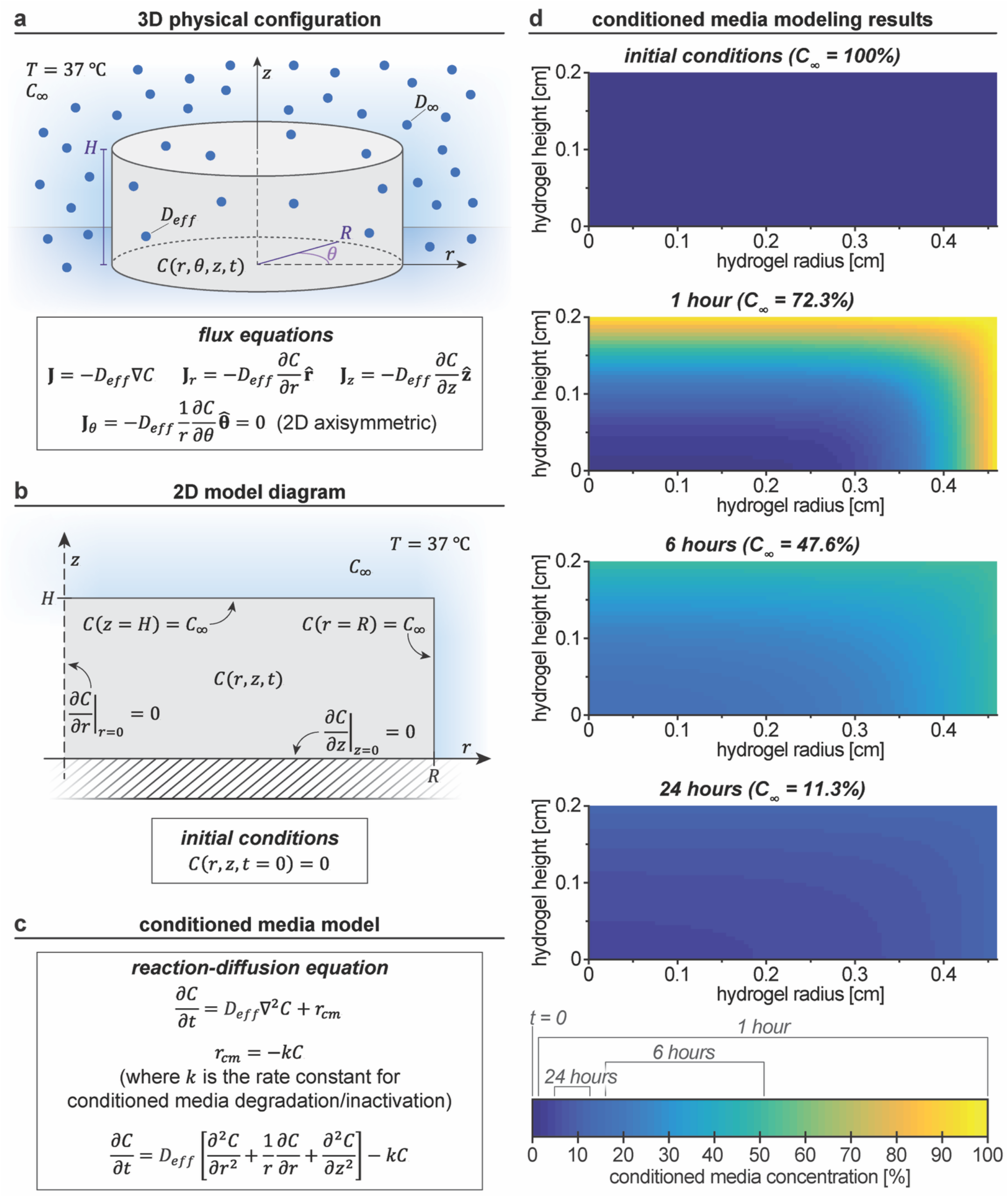
General reaction-diffusion modeling approach and conditioned media model results. a) Schematic of the 3D physical configuration of diffusion into a hydrogel, with the bulk solution concentration (*C*_*∞*_), diffusivity in water (*D*_*∞*_), and effective diffusivity (*D*_*eff*_, hindered diffusion in hydrogel) are indicated, in addition to the governing flux equations. b) Simplified 2D axisymmetric model diagram with boundary conditions and initial conditions (axis of symmetry at *r* = 0 and bottom of hydrogel sitting on the bottom of a well plate at *z* = 0). c) Governing reaction-diffusion equations for the CM model. d) CM modeling results (solved analytically) for select timepoints throughout the first 24-hour incubation (model parameter values available in **Table S7**). Additional timepoints are available in **Figure S20**.

#### 2.6.2. Diffusion and degradation of conditioned media in the 3D culture system

For simplicity, CM was modeled as a single protein species that degrades according to rate constant *k*. By assuming 2D axisymmetric diffusion, the continuity equation was then expressed as

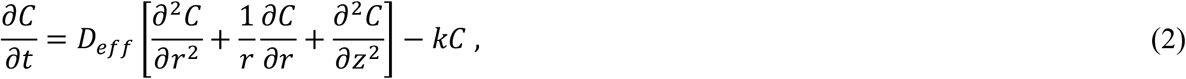

where *Deff* is the effective diffusivity within the hydrogel and *C* is the molar concentration of CM (**Figure 6b**,**c**). Boundary conditions (BCs, including conditions at the axis of symmetry and the no-flux assumption at the bottom of hydrogel) and initial conditions (ICs) are as follows:

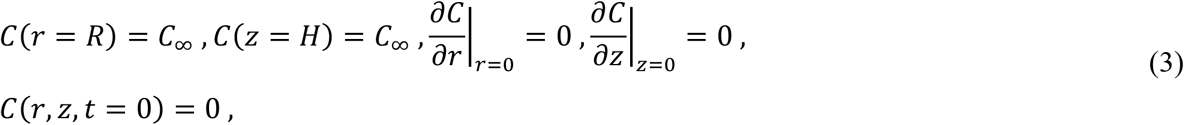

where *C*_∞_ is the concentration of CM in the solution surrounding the hydrogel. Toward solving this system, various transformations were performed to determine the full analytical solution for a constant *C*_∞_ (**Equation S18-S49**):

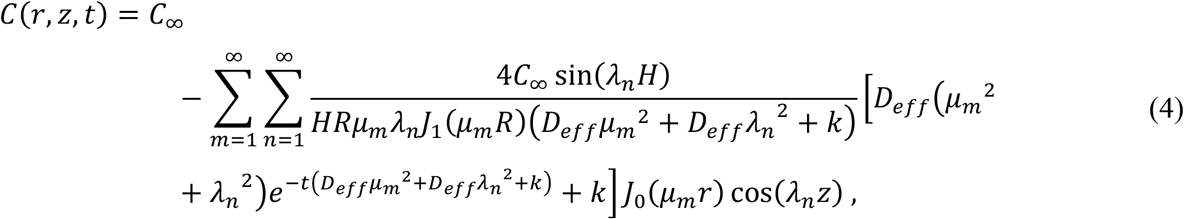

where µ*m* and λ*n* represents the Finite Hankle Transform and Finite Fourier Transform eigenvalues, respectively. This hydrogel reaction-diffusion system was modeled in MATLAB (MathWorks, Natick, MA), where the hydrogel concentration was (*i*) discretized in space to create a mesh in the *r-z* plane and (*ii*) discretized in time with interface flux calculations to account for a transient solution CM concentration, *C*_∞_ (**Equation S50-S64**).

The resulting model output for the first 24 hours is shown in **Figure 6d**, with additional timepoints available in **Figure S20**. These results suggest that the CM concentration within the hydrogel is dominated by solute diffusion during the first ∼5 hours, which is then eclipsed by protein degradation over the remaining time prior to the modeled media exchange, a trend that repeats after each ‘media exchange’. Paired with the cell motility results, the modeling data provides additional insight into fibroblast response to biochemical gradients. The motility measurements were conducted over timelapses approximate to the model timepoints of 0-12 ([*i*]), 12-24 ([*ii*]), 36-48 ([*ii*]), and 60-72 hours ([*iv*]). Of these timelapses, the steepest gradient and highest protein concentrations would be present during time course [*i*] (particularly in the first ∼6 hours), followed by a more shallow gradient with lower concentrations during time course [*ii*], and the lowest protein concentrations and most narrow gradient ranges during time courses [*iii*] and [*iv*]. At each timepoint, the fibroblasts exhibited significantly increased speed and directionality in response to CM (compared to fibroblasts in the control), demonstrating the ability of the cells to migrate in response to the protein gradient almost immediately. This is a point of interest, as within the CM condition, the lowest fibroblast directional persistence was observed during the first 24 hours of the experiment (timelapses [*i*] and [*ii*]), where the CM gradient is predicted to be most steep. Further, directional persistence increased significantly during timelapses [*iii*] and [*iv*], which were predicted to have the shallowest gradients and lowest CM concentrations. Based on these results and observations, we speculate that fibroblasts may be physically rearranging the local matrix in response to the steep CM protein gradient, taking advantage of the strain-yielding properties of the physical crosslinks between mfCMP fibrils^[34]^ to enable early cell motility (i.e., first ∼24 hours). At later timepoints (i.e., timelapses [*iii*] and [*iv*]), the fibroblasts have had more time to degrade the synthetic matrix through the MMP-degradable sites, which may account for the significant increase in directional persistence even at a relatively shallow protein gradient.^[92]^

#### 2.6.3. Diffusion and reaction of functionalized PEG in the 3D culture system

Modeling PEG diffusion into and reaction with the existing hydrogel network presents additional complexity. Multi-arm PEGs with reactive handles were modeled as a single species diffusing through the 3D cylindrical hydrogel based on the PEG diffusivity and then becoming fixed to the system based on the functional group reaction rate. We assumed that all “available” functional groups on the free PEG must react for the PEG to become fixed (**Figure 7a**). The number of “available” functional groups per PEG monomer (*f*) was calculated by *f* = 4 ×[available functional group on hydrogel prior to incubation]/[stiffening PEG functional group in solution at equilibrium], where the functional group concentrations are assumed to be homogeneously distributed throughout the hydrogel and full system, respectively (**Table S2**). The two reactions that govern this system were described as follows:

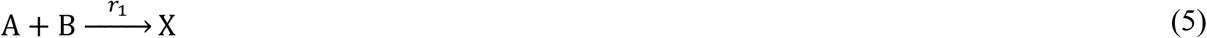

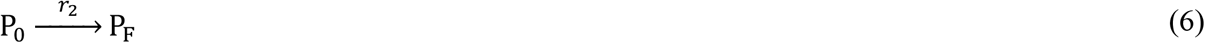

where A is the functional group fixed to the hydrogel, B is the “available” functional group on diffusing PEG, X is the resulting covalent linkage, P_0_ is unreacted and freely diffusing (not fixed) PEG, and P_F_ is fixed PEG after functional group reaction with the hydrogel. The associated reaction rates follow:

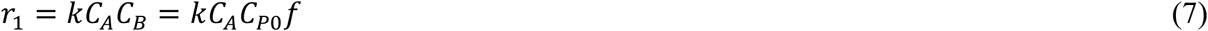

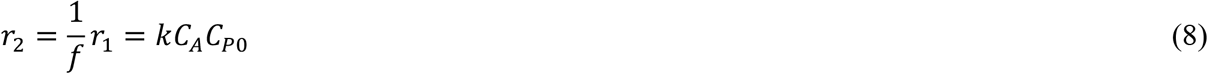

**Figure 7.**
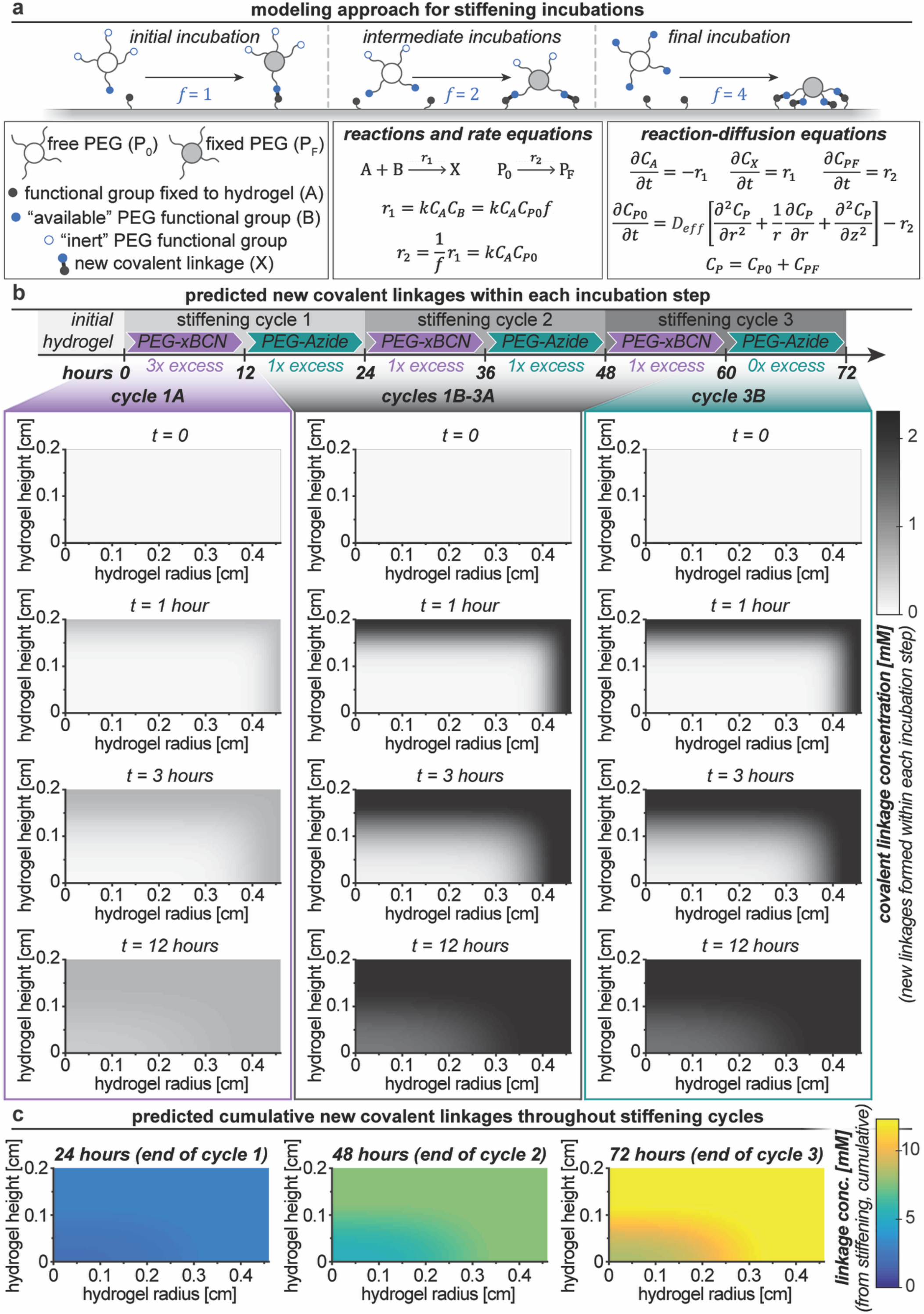
Reaction-diffusion modeling approach and results for stiffening incubations. a) Schematic of PEG reaction to the hydrogel network. The number of “available” functional groups per PEG molecule (*f*) were assigned according to the stoichiometric excess of PEG functional groups in the hydrogel at equilibrium, and all “available” functional groups on a single PEG were assumed to react for the PEG to move from freely diffusing (P_0_) to fixed to the hydrogel (P_F_). Reactions and appropriate rate equations, along with the corresponding governing reaction-diffusion equations, are indicated. b) Modeling results discretized and solved numerically for new covalent linkages formed within each 12-hour incubation step (model parameter values available in **Table S7**). Additional results for each incubation step are available in **Figure S21-S26.** c) Modeling results for cumulative new linkages formed over the course of 3-day stiffening at the end of each full stiffening cycle. New covalent linkage concentration due to stiffening ranges from 2.67–3.08 mM (24 hours), 6.77–7.70 mM (48 hours), and 10.9–12.3 mM (72 hours) throughout the hydrogel. Additional results for each stiffening incubation are available in **Figure S27**, including changes in PEG solution concentration.

The rate constant, *k*, describes the SPAAC reaction between an azide and a BCN group under physiologically relevant conditions.

This model required the concentrations of four components: hydrogel functional group, A; newly formed covalent linkage, X; diffusing PEG, P_0_; and fixed PEG, P_F_. As species A, X, and P_F_ are fixed to the hydrogel and unable to diffuse, the concentration equations for these three species will depend solely on reaction rate:

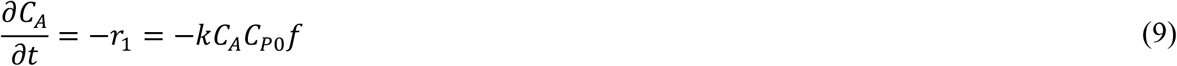

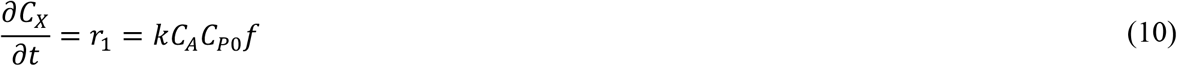

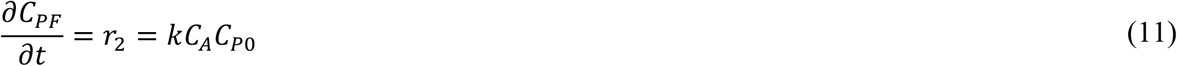

Conversely, free PEG will diffuse through the hydrogel until reacted and will depend on the concentration gradient of both free PEG and fixed PEG (i.e., the concentration of total PEG: *C*_*p*_=*C*_*po*_ + *C*_*PF*_), as both will contribute to the chemical potential gradient. In a 2D axisymmetric system, the continuity equation for free PEG was then expressed as

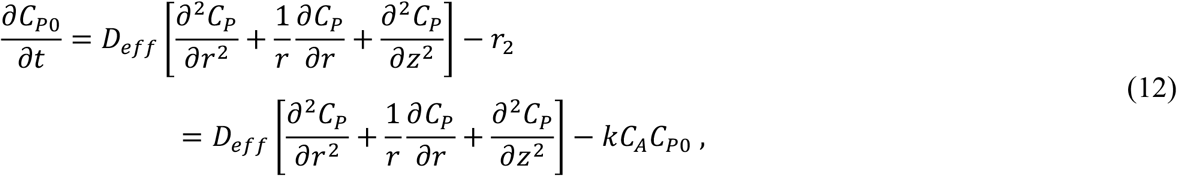

where *D*_*eff*_ is the effective diffusivity of the PEG monomer within the hydrogel (assumed to be constant, even with increased crosslink density over time). Boundary conditions (including conditions at the axis of symmetry and the no-flux assumption at the bottom of hydrogel) include

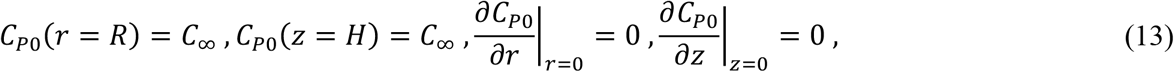

(where *C*_∞_ is the concentration of free PEG in the solution surrounding the hydrogel), with the following initial conditions for each species:

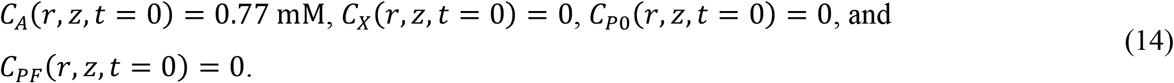

Due to the interdependence of the relevant species concentrations, we proceeded with a numerical solution (finite difference method) to solve this problem (**Equation S82-S112**). The non-diffusing species were discretized as follows:

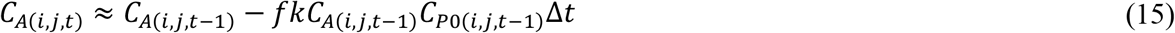

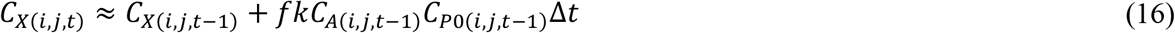

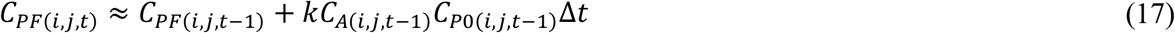

Discretization of **Equation 12** was more involved, where 6 separate equations were required to describe the system: one each for the internal nodes, the boundary condition at *r = R*, the boundary condition at *z = H*, the boundary condition at *z* = 0 (*r* ≠ 0), the axis of symmetry at *r* = 0 (*z* ≠ 0), and the special case where both *r* = 0 and *z* = 0 (**Equation 18-23**, respectively).

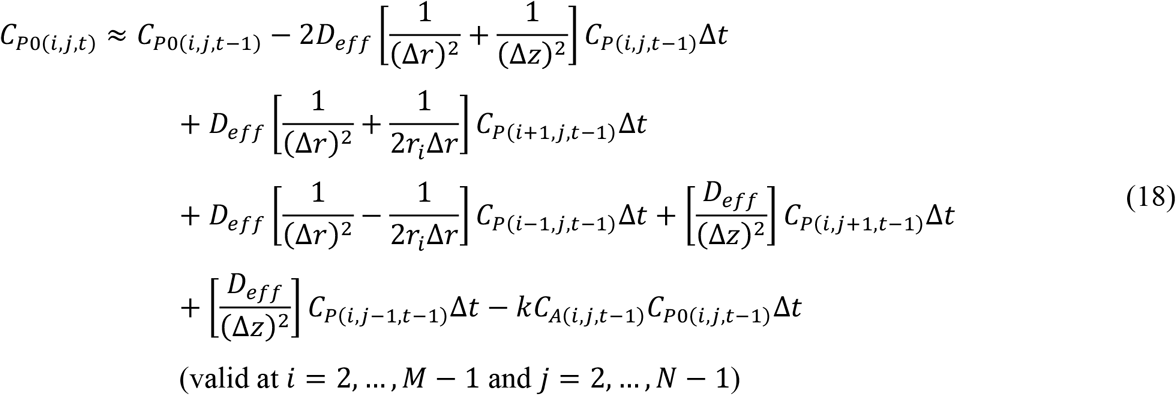

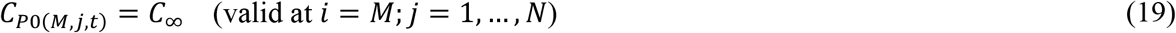

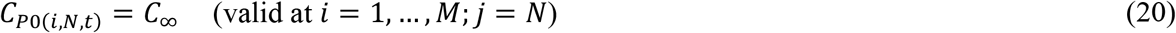

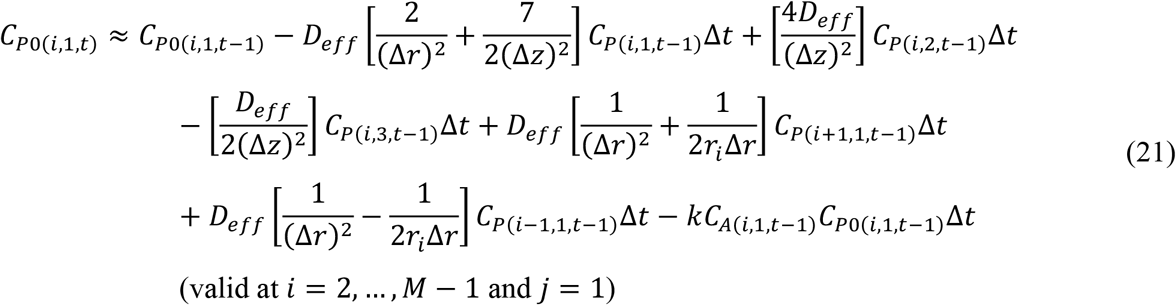

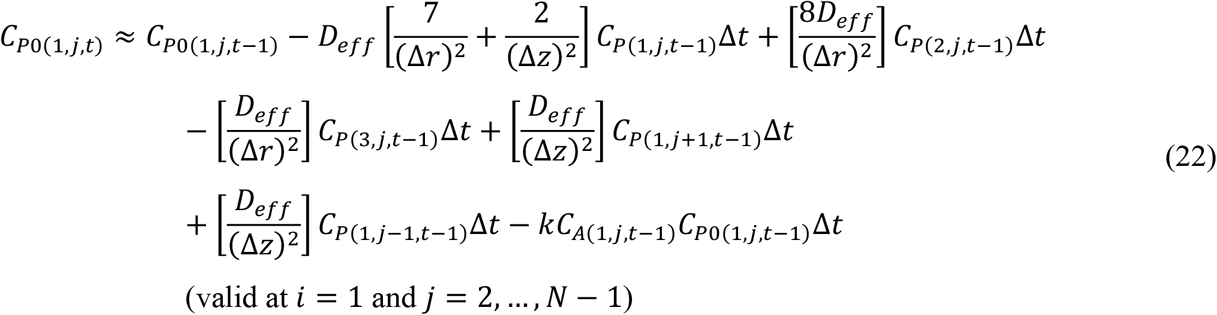

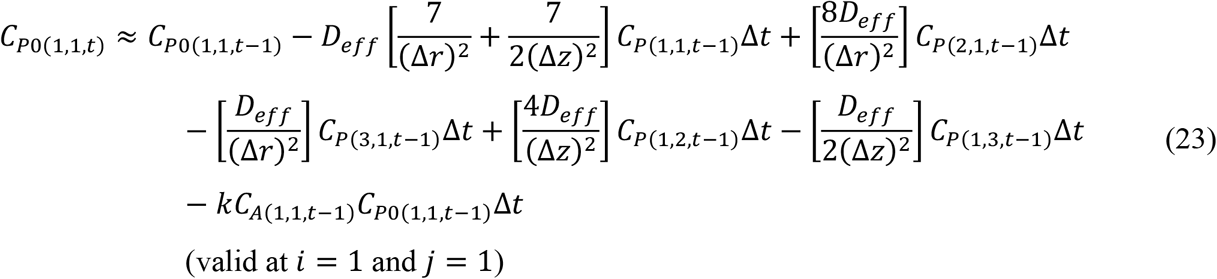

This system was then modeled in MATLAB using the same space discretization as in the CM model with one-second intervals for Δ*t*.

Key results of the model are displayed, including new covalent linkages formed within each incubation (**Figure 7b**; with additional outputs in **Figure S21-S26**) and the cumulative linkages formed during stiffening, shown at the end of each stiffening cycle (**Figure 7c**, with additional timepoints in **Figure S27**). These results indicate that the diffusing PEG does not fully react throughout the hydrogel over the course of our chosen 12-hour stiffening incubation, leading to a new linkage concentration (and corresponding crosslink density) gradient that compounds over stiffening cycles. As hydrogel modulus increases with increased crosslink density,^[93]^ this linkage concentration gradient likely results in a modulus gradient capable of promoting durotaxis,^[83]^ which we hypothesized to influence cell speed and directional persistence. Specifically, in response to hydrogel stiffening, we noted increased cell motility, where fibroblasts appeared to largely be migrating upward within the 3D culture system, in addition to moving in the *x*- and *y*-directions to a smaller extent. This observation corresponds with the gradient predicted in both the *r-* and *z-* directions by our stiffening model, where the gradient in the *z-*direction is at a smaller length scale than that of the *r-*direction, and therefore steeper, which has been shown to promote a stronger driving force for durotaxis.^[94-95]^ Further, our model predicts a cumulative linkage density increase of 8.35-12.3 mM throughout the hydrogel, corresponding to an approximate increase in crosslink density of *ρ*_*x*_= 4.04-5.97 mM. This increase is substantial and significant, as we estimate the initial hydrogel crosslink density to be ∼0.88 mM according to rubber elasticity theory.^[93]^ Such a significant increase in linkage density supports our hypothesis that the observed increase in fibroblast migration speed (relative to control) is in response to the modulus gradient; as the modulus continues to increase, the stiffness appears to counteract the gradient to some extent, slowing cell migration to the pace of the control, while continuing to promote more directional movement than observed in the control. Note, this model makes multiple assumptions, including constant hydrogel dimensions, constant *D*_*eff*_for the polymer, and no network loops or defects, which leads to an overestimate of the final crosslink density. Nevertheless, the model is an effective tool for insights into observed cell responses and has future utility for predicting and optimizing various experimental parameters.

#### 2.6.4. Model outlook

In this work, both the CM model and the stiffening model were used to compare predicted property gradients to cell motility results with the intention of gaining insight into fibroblast response to biochemical or biophysical stimuli. While these computational models were useful for understanding the experimental results within this work, they have the potential to be used as tools in other applications where approximating diffusion as 1-dimentional is not sufficient for a given geometry. These models also may be used to focus and winnow various experimental conditions rather than implementing a wide range of preliminary experimental conditions, saving valuable time and resources. For example, the CM model could be modified to reflect a different protein of interest and predict how frequently media replacements are required and what protein concentrations are needed to maintain a particular gradient range or steepness. Alternatively, one could employ a version of the stiffening model to predict the experimental conditions (e.g., incubation times, solution concentrations) needed to achieve a particular crosslinking result, whether that is uniform polymer attachment throughout the existing hydrogel or a specific crosslinking density gradient. Ultimately, the goal is to provide a tool to improve understanding of the physical implications of multidimensional reaction-diffusion kinetics and the resulting cell response.

## 3. Conclusion

Here, we have developed an approach for hydrogel stiffening that affords significant tunability, from initial hydrogel composition to the degree and rate of increase in modulus over time. This system was established with the goal of providing a platform for modeling a range of dynamic processes, including healthy versus maladaptive wound healing toward fibrotic disease progression. Importantly, a stiffening process that takes place on the order of days to weeks, a time scale consistent with stiffening in native tissues, is achieved with the combination of the relatively slow reaction rate of the BCN group amongst other cyclooctynes, the diffusion-limited nature of the stiffening polymer, and multiple cycles of stiffening. Toward exploring the timescales of this stiffening system, we included a numerical solution to the reaction-diffusion problem presented by this cylindrical hydrogel geometry and designed a computational model to better understand the kinetics of the stiffening process. The computational model to approximate the reaction-diffusion characteristics will be enabling in future hypothesis testing, including in probing various cell responses to matrix stiffening and treatment with soluble factors. With the initial use of the stiffening system, we have demonstrated the efficacy of the stiffening approach and explored the impact on fibroblast response. The modulus range explored in this work 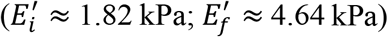 is particularly relevant for modeling early stages of maladaptive wound healing and fibrotic lung disease. While fixed cell immunostaining provides valuable insights into various cell responses, this study highlights the importance of real-time monitoring to better understand how cells dynamically react to changes in their environment. Active hydrogel stiffening had a notable and sustained effect on fibroblast behavior, marked by an apparent phenotypic shift from migratory fibroblasts to contractile myofibroblasts. This transition was indicated by a significant increase in αSMA expression, along with an initial phase of rapid cell motility that progressively slowed as stiffness and αSMA levels increased. Given the tunability of the stiffening approach developed here, this system may be adapted to model various tissues and ECM stiffening applications. This flexibility could provide an approach to studying disease-specific biomarkers for early detection of hard-to-diagnose conditions and offers a platform to investigate the effects of early intervention.

## Supporting Information

Supporting Information is available: Experimental methods (materials synthesis and characterization methods, hydrogel methods, cell methods, image analysis methods, mathematical modeling methods, statistical methods); List of symbols; Supplemental calculations; Reaction schemes (PEG-SH, LAP, PEG-xBCN); NMR results (PEG-SH, LAP, PEG-xBCN); Peptide structures and mass spectrometry results (non-assembling peptide linker, RGDS, mfCMP-Azide); Circular dichroism (mfCMP-Azide); TEM images (assembled mfCMP-Azide); Additional rheometry results; Detailed metabolic activity results with multiple comparisons analysis; Cell density results; Additional cell fluorescence analysis; Fibroblast morphology analysis; Additional live reporter images; Detailed reporter analysis; Detailed motility analysis; Additional computational model results (CM, stiffening); Hydrogel volume results; Hydrogel formulation table; Stiffening solution concentrations table; Statistical analysis tables; and Model parameter values (PDF).

## Acknowledgements

The authors would like to acknowledge financial support from the New Innovator Award funded by the National Institutes of Health (NIH) (DP2-HL152424) and the University of Delaware Center for Hybrid, Active, and Responsive Materials (UD CHARM) Materials Research Science and Engineering Center (MRSEC) program supported by the National Science Foundation (NSF) (DMR-2011824). Support for instrumentation at the University of Delaware was provided in part by the Delaware Centers of Biomedical Research Excellence (COBRE) program supported by a grant funded by an Institutional Development Award from the National Institute of General Medical Sciences (NIGMS) of the NIH (P30 GM110758-02) and the UD CHARM MRSEC program. This publication was also made possible by UD Core facilities including the Keck Center for Advanced Microscopy and Microanalysis and the University of Delaware NMR and Mass Spectrometry Core facilities. The authors would like to acknowledge the Delaware Biotechnology Institute (DBI) Bio-Imaging Center—in particular Dr. Chandran Sabanayagam—for access to and support surrounding the Imaris image analysis software. Finally, the authors would also like to thank Dr. Kartik Bomb for providing the macrophage-conditioned media and Prof. Antony Beris and Prof. Phillip Taylor for discussions around solving the 2D reaction-diffusion problems.

## Author Contributions

E.M.F. and A.M.K. conceived the ideas and designed the experiments. E.M.F. synthesized and characterized materials (excluding PEG-xBCN and BCN precursors). B.P.S. synthesized and characterized (1*R*,8*S*,9*R*)-bicyclo[6.1.0]non-4-yn-9-yl-methyl (4-nitrophenyl) carbonate and S.L.S. functionalized and characterized the PEG-xBCN. S.E.C produced and supplied the reporter cell line. E.M.F. performed all experiments and conducted the mathematical modeling. E.M.F. and A.M.K. analyzed the data. The manuscript was written through contributions of all authors. All authors have given approval to the final version of the manuscript.

## Table of Contents (TOC)

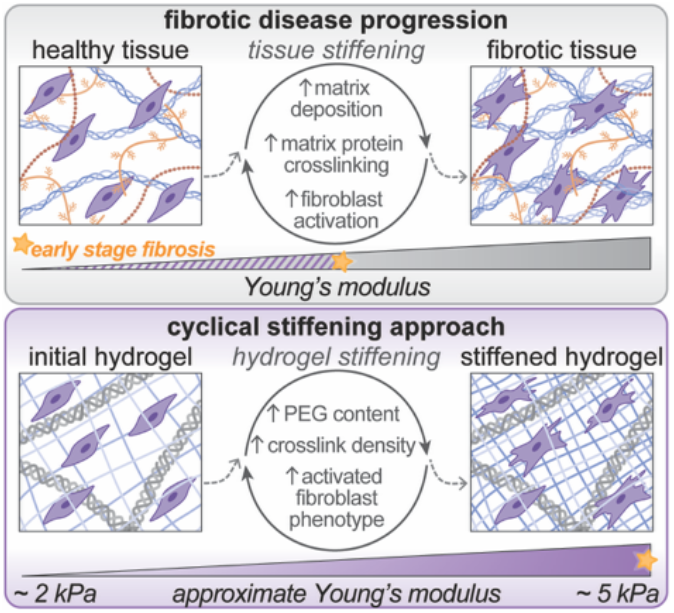

A synthetic extracellular matrix with collagen-like structure and user-controlled, bio-orthogonal cyclic stiffening provides a fully synthetic system to probe dynamic fibroblast activation over relevant timescales. Matrix stiffening promotes a pro-fibrotic fibroblast phenotype and durotactic migration, establishing a robust platform for property modulation and stiffening independent from the initial matrix composition for studying cell responses in the context of early-stage fibrosis.

