## Supporting Information for "Matrix stiffening toolbox: dynamic hydrogels for three-dimensional cell culture with real-time cell response"

S. L. Swedzinski, A. M. Kloxin
Department of Materials Science and Engineering
University of Delaware
Newark, DE 19716, USA

Keywords: bioinspired synthetic hydrogel-based extracellular matrices, bio-orthogonal click chemistry, matrix stiffening, dynamic 3D cell cultures, mathematical modeling, real-time fibroblast activation

### 1. Experimental Methods

#### 1.1 Materials synthesis and characterization methods

*PEG-SH functionalization and characterization*

To produce the four-arm poly(ethylene glycol) tetrathiol (PEG-SH) macromer used in this work, a four-arm poly(ethylene glycol) (PEG-OH; JenKem Technology USA, Plano, TX) was functionalized with thiol end groups, according to well-established methods (**Figure S1**).[1-2] In brief, PEG-OH ($M_n$ ~ 20,000 g $mol^{-1}$, 10 g) was reacted with allyl bromide (Thermo Fisher Scientific, Waltham, MA) to form allyl ether-functionalized PEG (PEG-Allyl Ether); the allyl ether end groups were then reacted with thioacetic acid (Acros Organics, Fair Lawn, NJ) through photo-initiated radical thiol–ene click chemistry, resulting in a PEG-Thioacetate monomer; and finally, the thioacetate end group was deprotected by reaction with concentrated sodium hydroxide (NaOH; Thermo Fisher Scientific, Waltham, MA) to yield the final PEG-SH product. To ensure the complete reduction of any disulfide bonds, PEG-SH was treated with tris(2-carboxyethyl)phosphine (TCEP; Millipore Sigma, St. Louis, MO) in deionized (DI) water (350 mg TCEP in ~40 mL water per 1 g PEG-SH) for 16 hours with continuous stirring. After TCEP treatment, the PEG-SH was dialyzed (MWCO 1 kDa; Spectrum Laboratories, New Brunswick, NJ) against acidified DI water (pH 4) for 24 hours, frozen, and lyophilized. The success of modification was confirmed and functionality determined by $^1$H-NMR (Neo 400 MHz NMR spectrometer; Bruker Corporation, Billerica, MA) (**Figure S2**). Stock solutions of PEG-SH (55 mM thiol end group concentration, confirmed with Ellman's assay[3]) were prepared in sterile Dulbecco's phosphate-buffered saline (DPBS; Thermo Fisher Scientific, Waltham, MA).

*Peptide synthesis, purification, and characterization*

A total of 3 peptides were synthesized for this work: a non-assembling linker peptide (bis-allyoxycarbonyl (alloc)), an integrin-binding peptide (mono-alloc), and a multifunctional collagen-mimetic peptide (mfCMP) containing two functional handles (alloc and azide). The non-assembling linker peptide (KK(alloc)GGPQG↓IWGQGK(alloc)K, referred to as linker peptide; **Figure S3a**) is a matrix metalloproteinase-degradable peptide variant derived from collagen, and the integrin-binding peptide (K(alloc)GWGRGDS, referred to as RGDS; **Figure S4a**) is derived from a range of extracellular matrix proteins including multiple collagens, fibronectin, and vitronectin.[2, 4-6] These peptides were included to promote cell-mediated degradation and remodeling of the hydrogel and cell-matrix interactions, respectively. The mfCMP (K(azide)G$(PKG)_4$PK(alloc)G$(POG)_6$$(DOG)_4$, referred to as mfCMP-Azide; **Figure S5a**) was included to impart fibrillar structure. In this work, K(alloc) represents an alloc-protected lysine amino acid residue (providing a reactive handle for photo-initiated thiol–ene click chemistry), K(azide) an azide-protected lysine amino acid residue (providing a reactive handle for azide-alkyne click chemistry), and O a hydroxyproline residue. All other amino acids are represented by standard abbreviations.

All three peptides were synthesized using microwave-assisted solid phase peptide synthesis (SPPS) (Liberty Blue; CEM Corporation, Matthews, NC). In brief, the non-assembling linker peptide and integrin-binding peptide were synthesized in 0.25 mmol batches, using standard MBHA rink-amide resin (0.51 meq $g^{-1}$; NovaBiochem, San Diego, CA) in a 125 mL Liberty Blue reaction vessel to prevent solvent overflow and enhance nitrogen bubbling mixing efficacy, following established protocols.[1] Fmoc deprotections were performed with 20% piperidine (Millipore Sigma, St Louis, MO) in standard dimethylformamide (DMF; Thermo Fisher Scientific, Waltham, MA) at 75 °C. Each amino acid residue (ChemPep, Wellington, FL) was double coupled at 75 °C for 8 minutes using a 5× molar excess of the activating reagents N,N'-diisopropylcarbodiimide (DIC, 1 M in DMF; AppTech, Carlsbad, CA) and ethyl(hydroxyimino) cyanoacetate (OxymaPure, 1 M in DMF; CEM Corporation, Matthews, NC). The synthesis of mfCMP-Azide followed a similar protocol (in 0.1 mmol batches on TentaGel Resin using the standard 35 mL Liberty Blue reaction vessel) but with the established modifications specific to residues prone to deletions (TentaGel Resin: 0.19 meq $g^{-1}$; Peptides International, Louisville, KY).[7]

All three peptides were cleaved from their respective resins for 2 hours using the following cleavage solution: TFA/TIPS/DI water (95%/2.5%/2.5% v/v) with 2.5% w/v phenol. (TFA: trifluoroacetic acid; TIPS: triisopropylsilane; TFA and TIPS from Thermo Fisher Scientific, Waltham, MA; phenol from MilliporeSigma, Burlington, MA.) After cleavage, the peptides were precipitated in diethyl ether (Thermo Fisher Scientific, Waltham, MA), and washed 3× with fresh diethyl ether, as previously described.[7] The washed peptides were dried overnight and then purified by reverse phase high performance liquid chromatography (HPLC; XBridge C18 OBD 5 µm column; Waters Corporation, Milford, MA). The linker peptide was purified using a linear acetonitrile gradient from 28 to 35%, RGDS was purified with a gradient from 22 to 28%, and mfCMP-Azide was purified with a gradient from 20 to 30% (acetonitrile: Thermo Fisher Scientific, Waltham, MA).

The collected HPLC fractions were frozen and lyophilized, and the product identities were confirmed using electrospray ionization (ESI+) mass spectrometry (either Xevo G2-S QTof or Single Quadruple Detector 2 SQD2; Waters Corporation, Milford, MA) (**Figure S3b**, **Figure S4b**, and **Figure S5b**). mfCMP-Azide was then dialyzed first against acidified DI water (pH 4, 36 hours) and then against neutral DI water (24 hours), followed by freezing and lyophilization, per the previously established protocol.[7] The lyophilized peptide was stored at -80 °C until use. Stock solutions for each peptide were prepared in sterile DPBS. mfCMP-Azide stock solution concentration (5 mM) was determined by dry weight, while linker peptide (71.1 mM alloc functional handle) and RGDS peptide (43.3 mM) stock solution concentrations were confirmed with ultraviolet-visible (UV-VIS) spectrometry (NanoDrop 2000c Spectrophotometer; Thermo Fisher Scientific, Waltham, MA), where the tryptophan absorbance was measured at 280 nm and converted to peptide and/or functional handle concentration using Beer's law.[1]

*mfCMP-Azide assembly and higher order structural characterization*

The assembling peptide mfCMP-Azide [K(azide)$(PKG)_4$PK(alloc)G$(POG)_6(DOG)_4$] allows the incorporation of collagen type-I-like assembled structure into a fully synthetic hydrogel matrix. Specifically, these mfCMPs are made up of five key components: *(i)* a hydrogen bonding block, *(ii)* a positively charged block, *(iii)* a negatively charged block, *(iv)* an alkene functional group, and *(iv)* an azide functional group. *(i)* Proline-hydroxyproline-glycine (POG) repeat units located in the middle of the peptide promote interpeptide hydrogen bonding to facilitate triple helix formation.[8-9] The two charged blocks enable end-to-end peptide interactions for the formation of elongated fibrillar structures: *(ii)* proline-lysine-glycine (PKG) repeat units form a positively charged block on the N-terminus of the peptide and *(iii)* aspartic acid-hydroxyproline-glycine (DOG) repeat units form a negatively charged block on the C-terminus of the peptide.[10] *(iv)* An allyloxycarbonyl-protected lysine residue (K(alloc)) provides an alkene functional group for participation in a photoinitiated thiol–ene reaction for covalent integration into a larger hydrogel network.[1, 11] Finally, *(v)* an azide-protected lysine residue (K(azide)) provides an unreacted functional group for later hydrogel modification.[7]

For assembly, mfCMP-Azide solutions were incubated on a heat block set to 85 °C for 15 minutes to dissociate any assembled peptide into individual peptide strands. Solutions were then slowly cooled to room temperature for continued assembly over the next 48 hours, as previously described.[7] Assembled mfCMP-Azide solutions were aliquoted, frozen, and lyophilized in preparation for use in hydrogel formation. Higher order structures were characterized using circular dichroism for verification of triple helix formation and determination of melting temperature (**Figure S5c**; assembled at 0.3 mM in DPBS; 1500 Circular Dichroism Spectrophotometer; JASCO Corporation, Easton, MD) and transmission electron microscopy for verification of fibril formation (**Figure S5d**; assembled at 1 mM DPBS and diluted to 0.3 mM in DI water for drop casting; Talos Transmission Electron Microscope; FEI Company, Hillsboro, OR), according to previously well-defined protocols for each technique.[7]

*LAP synthesis and characterization*

The photoinitiator lithium phenyl-2,4,6-trimethylbenzoylphosphinate (LAP), was synthesized according to a previously published protocol (**Figure S6a**).[2] In brief, equimolar amounts of 2,4,6-trimethylbenzoyl chloride (MilliporeSigma, Burlington, MA) and dimethylphenylphosphonite (Acros Organics, Fair Lawn, NJ) were reacted at room temperature for 16 hours under argon. Lithium bromide (4× molar excess; MilliporeSigma, Burlington, MA) in 2-butanone (MilliporeSigma, Burlington, MA) was then added, and the mixture was heated to 50 °C and stirred for 10 minutes. The resulting precipitate was filtered, rinsed 3× with 2-butanone, and dried under vacuum. The product identity was confirmed by $^1$H-NMR (Bruker AVIII400; Bruker Corporation, Billerica, MA) (**Figure S6b**). Stock solutions of LAP (40 mM, based on desiccated dry weight) were prepared in sterile DPBS after desiccation under vacuum for at least 2 hours.

*PEG-xBCN functionalization and characterization*

First, the compound (1*R*,8*S*,9*R*,4*Z*)-bicyclo[6.1.0]non-4-ene-9-yl-methanol was synthesized according to previously reported methods (**Figure S7a**).[12-14] Briefly, rhodium dimer tetraacetate (3.4 meq relative to ethyl diazoethyl ester; Combi-Blocks, San Diego, CA) was dissolved in (*Z*,*Z*)-1,5-cyclooctadiene (10 molar eq relative to ethyl diazoethyl ester; TCI America, Portland, OR), ethyl diazoethyl ester (containing ≥ 13 wt% dichloromethane (DCM); MilliporeSigma, Burlington, MA) was added dropwise, and the reaction proceeded for an hour under nitrogen. The crude product (ethyl (1*R*,8*S*,9*S*,4*Z*)-bicyclo[6.1.0]non-4-ene-9-yl-10-ate (*endo*-**Compound 1**) and ethyl (1*R*,8*S*,9*R*,4*Z*)-bicyclo[6.1.0]non-4-ene-9-yl-10-ate (*exo*-**Compound 1**), non-selective) was concentrated and purified by column chromatography (0-10% ethyl acetate (Thermo Fisher Scientific, Waltham, MA) gradient in hexanes (Thermo Fisher Scientific, Waltham, MA)). In the next step, anhydrous diethyl ether (Thermo Fisher Scientific, Waltham, MA) was added to potassium tert-butoxide (3 molar eq relative to **Compound 1**; Acros Organics, Geel, Belgium), followed by DI water (1.25 molar eq relative to **Compound 1**). The *endo*/*exo* mixture of **Compound 1** (where *exo* was the major product) was added, and the solution was stirred overnight at room temperature. A sodium hydroxide solution (10%) and DI water were then added, and the aqueous phase was collected (with an additional 10% sodium hydroxide wash of the organic phase), cooled in an ice bath, and acidified (to pH < 2) with concentrated hydrochloric acid (Thermo Fisher Scientific, Waltham, MA). The resulting precipitate ((1*R*,8*S*,9*R*,4*Z*)-bicyclo[6.1.0]non-4-ene-9-carboxylic acid (**Compound 2**)) was collected by filtration, washed with 0.1 N hydrochloric acid, and vacuum-dried overnight. Next, a suspension of lithium aluminum hydride ($LiAlH_4$; 3.5 molar eq relative to **Compound 2**; MilliporeSigma, Burlington, MA) and anhydrous tetrahydrofuran (THF; Thermo Fisher Scientific, Waltham, MA) was chilled on ice. **Compound 2** was dissolved in anhydrous THF, slowly added dropwise to the $LiAlH_4$ solution, and reacted for 2 hours at room temperature. The reaction was then chilled and quenched with DI water, followed by filtration to remove alumina and lithium salts (which were then washed with diethyl ether). The filtration flow through and diethyl ether wash were combined, and an extraction with DI water was performed, setting aside the organic phase. The aqueous phase was reextracted with diethyl ether, and the organic phase was collected. The organic phases were combined, dried over sodium sulfate (Thermo Fisher Scientific, Waltham, MA), filtered, and concentrated, resulting in the (1*R*,8*S*,9*R*,4*Z*)-bicyclo[6.1.0]non-4-ene-9-yl-methanol product (**Compound 3**).

Next, the compound (1*R*,8*S*,9*R*)-bicyclo[6.1.0]non-4-yn-9-yl-methanol was synthesized according to previously reported methods (**Figure S7b**).[12, 15] Briefly, **Compound 3** was dissolved in anhydrous DCM (MilliporeSigma, Burlington, MA), cooled in an ice bath, and bromine (approximately 1 molar eq relative to **Compound 3**; MilliporeSigma, Burlington, MA) in anhydrous DCM was slowly added. Once the mixture was consistently yellow, it was stirred for 5 minutes, followed by quenching with a 10% sodium thiosulfate solution (Thermo Fisher Scientific, Waltham, MA). The resulting solution was extracted twice with standard DCM (Thermo Fisher Scientific, Waltham, MA), and the organic phases were combined, dried over magnesium sulfate (Thermo Fisher Scientific, Waltham, MA), filtered, and concentrated with rotary evaporation, resulting in the crude product, (1*R*,8*S*,9*R*)-4,5-dibromobicyclo[6.1.0]nonan-9-yl-methanol (**Compound 4**). Crude **Compound 4** was then dissolved in anhydrous THF in an ice bath, and a solution of potassium tert-butoxide (2.5 molar eq relative to **Compound 4**; approximately 1 N in THF, as purchased; MilliporeSigma, Burlington, MA) was added dropwise. The mixture was stirred for 5 minutes on the ice bath, then fitted with a reflux condenser and transferred to an oil bath. The reaction proceeded for 2.5 hours at 75 °C under reflux, then was removed from the oil bath and stirred at room temperature for an additional 30 minutes. Next, the reaction was slowly quenched with a saturated aqueous ammonium chloride solution (MilliporeSigma, Burlington, MA). The product, (1*R*,8*S*,9*R*)-bicyclo[6.1.0]non-4-yn-9-yl-methanol (**Compound 5**), was extracted into standard DCM, dried over magnesium sulfate, filtered, and concentrated. **Compound 5** was then purified with flash chromatography (0-20% ethyl acetate in hexanes).

The compound (1*R*,8*S*,9*R*)-bicyclo[6.1.0]non-4-yn-9-yl-methyl (4-nitrophenyl) carbonate was then synthesized according to previously reported methods (**Figure S7c**).[12, 16] Briefly, **Compound 5** was dissolved in anhydrous DCM and pyridine (2.5 molar eq relative to **Compound 5**; MilliporeSigma, Burlington, MA) was added, followed by nitrophenyl chloroformate (1.1 molar eq relative to **Compound 5**; TCI America, Portland, OR). The reaction proceeded for 30 minutes, followed by quenching with a saturated aqueous ammonium chloride solution. The organic phase was collected, and the aqueous phase was reextracted 3× with standard DCM. The organic phases were combined, dried over magnesium sulfate, filtered, and concentrated with rotary evaporation, resulting in the crude product, (1*R*,8*S*,9*R*)-bicyclo[6.1.0]non-4-yn-9-yl-methyl (4-nitrophenyl) carbonate (**Compound 6**). **Compound 6** was then purified with flash chromatography (0-25% ethyl acetate in hexanes).

Finally, a four-arm PEG Amine (PEG-$NH_2$; JenKem Technology USA, Plano, TX) was functionalized with the synthesized bicyclononyne groups according to previously reported methods (**Figure S7d**).[12, 17] Briefly, PEG-$NH_2$ ($M_n \sim 10{,}000$ g mol$^{-1}$, no more than 1 g per batch) and **Compound 6** (1.5 molar eq relative to $NH_2$ groups on the PEG) were fully dissolved in a minimal volume of extra dry DMF (Thermo Fisher Scientific, Waltham, MA). Diisopropylethylamine (DIPEA; MilliporeSigma, Burlington, MA) was then added to the solution (4 molar eq relative to $NH_2$ groups on the PEG) and the reaction proceeded for 24 hours in the dark. The polymer product was precipitated into cold diethyl ether and held at -20 °C for 15 minutes, followed by centrifugation. The solvent was decanted, and the polymer pellet was redissolved in standard DMF. Once again, the polymer product in DMF was precipitated into cold diethyl ether, held at -20 °C for 15 minutes, centrifuged, and the solvent decanted. The pellet was washed 2× with cold diethyl ether and dried under a steady stream of nitrogen then further dried in a vacuum desiccator. The polymer product was dissolved in DI water, syringe-filtered (0.45 µm PES filter; VWR, Radnor, PA), and dialyzed against DI water for 36-48 hours (SnakeSkin, 3.5 kDa MWCO, free from sodium azide; Thermo Fisher Scientific, Waltham, MA). Finally, the dialysis product was collected, frozen, and lyophilized to yield the final four-arm poly(ethylene glycol) tetra-bicyclononyne (*exo*) (PEG-xBCN) product, which was stored at -20 °C until use. The product identity was confirmed and functionality determined by $^{1}$H-NMR (Bruker AVIII600; Bruker Corporation, Billerica, MA) (**Figure S8**).

##### 1.2 Hydrogel methods

*Hydrogel formation*

Stock solutions of PEG-SH (55 mM SH, 4 functional groups per polymer), LAP (40 mM), linker peptide (71.1 mM alloc, 2 functional groups per peptide), RGDS (43.4 mM alloc, 1 functional group per peptide), and mfCMP-Azide (5 mM alloc, 5 mM azide, 1 of each functional group per peptide) were prepared in sterile DPBS. mfCMP-Azide stocks were melted and assembled according to the methods described above (*mfCMP-Azide assembly and higher order structural characterization*, derived from previously established protocols[1, 7, 11]), aliquoted, flash frozen in liquid nitrogen, and lyophilized. Hydrogel precursor solutions were prepared such that they contained 10 wt% PEG-SH (20 mM SH), 2.2 mM LAP, 6.5 mM linker peptide (13 mM alloc), 2 mM RGDS, and DPBS. The lyophilized mfCMP-Azide was brought to room temperature and resuspended directly in the precursor solution for a final mfCMP-Azide concentration of 5 mM alloc within the final hydrogel precursor solution (for an overall 1:1 thiol:alloc ratio, **Table S1**). Hydrogel precursor solutions then were transferred to syringe molds (20 μL into a 1 mL syringe with the tip removed; Thermo Fisher Scientific, Waltham, MA) and irradiated with a cytocompatible dose of long wavelength UV light to complete the photo-initiated radical thiol–ene crosslinking reaction (4 minutes, 365 nm, 10mW $cm^{-2}$; Exfo Omnicure Series 2000 light source, 365nm bandpass filter; Excelitas Technologies Corp., Waltham, MA).[1]

*Stiffening protocol*

Hydrogel stiffening of equilibrium swollen hydrogels was achieved by conducting cycles of sequential incubations: first with a four-arm poly(ethylene glycol) bicyclo[6.1.0]nonyne (PEG-xBCN, 10,000 g $mol^{-1}$, 85% functionality) followed by a four-arm poly(ethylene glycol) azide (PEG-Azide, 10,000 g $mol^{-1}$, 99% functionality; Creative PEGWorks, Durham, NC). For the work presented here, three cycles of stiffening were performed as detailed below.

Briefly, lyophilized vials of PEG-xBCN and PEG-Azide were brought to room temperature and desiccated under vacuum for at least 2 hours prior to weighing. Stock solutions of PEG-xBCN and PEG-Azide were prepared in sterile DPBS (50 mM BCN and 50 mM Azide, respectively, accounting for the percent functionality of the PEG monomer). Working stiffening solutions were prepared by diluting the appropriate stock solution into pre-warmed (37 °C) cell culture media with 1% fetal bovine serum (FBS) to achieve the desired stiffening solution concentration (see **Table S2** for the specific concentrations used in this work). (*Note*: cell culture media with 1% FBS consisted of Ham's F12K media supplemented with 50 U $mL^{-1}$ penicillin, 50 μg $mL^{-1}$ streptomycin, 0.2% v:v amphotericin B/Fungizone, and 1% v:v FBS; all from Thermo Fisher Scientific, Waltham, MA.) For each cycle of stiffening, the solution surrounding the hydrogel was removed, the hydrogel was briefly rinsed with warmed, fresh media, and each hydrogel was moved to a new well of the 48-well plate (non-tissue culture treated; CELLTREAT; Pepperell, MA). PEG-xBCN stiffening solution (250 μL) at the appropriate concentration was then added to the hydrogel and incubated at 37 °C for approximately 12 hours. After the PEG-xBCN incubation, the solution was removed and the hydrogel was briefly rinsed with warmed, fresh media and moved to a new well. PEG-Azide stiffening solution (250 μL) at the appropriate concentration was then added to the hydrogel and incubated at 37 °C for approximately 12 hours. After the final cycle of stiffening, hydrogels were washed 3× for 1 hour each in warmed, fresh media to remove any remaining monomers.

Specific to the stiffening protocol used in this work, hydrogels at the time of preparation (20 µL) contained 5 mM free azide groups, presented on the mfCMP-4Azide. After swelling to approximately 130 µL, the azide concentration decreased to 0.77 mM. The initial cycle of PEG-xBCN stiffening solution (Cycle 1A, 250 µL, 4.68 mM BCN) was added such that after diffusion, the BCN concentration within the hydrogel would reach 3.08 mM (3× molar excess). For following stiffening steps of PEG-Azide (Cycle 1B) → PEG-xBCN (Cycle 2A) → PEG-Azide (Cycle 2B) → PEG-xBCN (Cycle 3A), the stiffening solutions (250 µL, 7.02 mM functional group) were each added such that after diffusion, the stiffening solution functional group concentration would reach 4.62 mM (1× molar excess) within the hydrogel. In the final cycle of PEG-Azide stiffening, the stiffening solution (Cycle 3B, 250 µL, 3.51 mM Azide) was added such that after diffusion, the azide concentration within the hydrogel would reach 2.31 mM (equimolar to the expected free BCN in the hydrogel), resulting in a BCN:Azide stoichiometric ratio of 1:1.

*Rheometry on equilibrium swollen hydrogels*

Twenty-microliter syringe-mold hydrogels were formed according to the protocol described above, transferred to 48-well plates (non-tissue culture treated) with cell culture media, and incubated under conditions relevant for mammalian cell culture (i.e., 37 °C, 5% $CO_2$, humid conditions). After equilibrium swelling and stiffening treatment (all under mammalian cell culture conditions), the hydrogels were briefly rinsed with warmed fresh media. The diameters were measured, and the hydrogels were transferred to a Discovery HR-30 rheometer with the Peltier plate set to 37 °C and set up with sandblasted top and bottom geometries (TA Instruments, New Castle, DE). The gap height was adjusted to match the height of each hydrogel, with a slight axial normal force (0.01 N) applied to prevent slipping between the hydrogel and the geometry. Oscillatory shear storage modulus ($G'$) and loss modulus ($G''$) measurements were then taken over 90 seconds using parameters within the linear viscoelastic regime (1% strain, 2 rad $s^{-1}$ frequency).

Young's modulus ($E$) of the materials was approximated from the shear storage modulus results using rubber elasticity theory and assuming the hydrogel behaves as an isotropic homogeneous material. Briefly, Young's modulus was calculated according to the equation, $E = 2G'(1 + v)$, where the material is assumed to be incompressible and thus $v = 0.5$. For each condition, at least 4 hydrogels were measured.

##### 1.3 Cell methods

*Fibroblast 2D cell culture and expansion*

Lentivirus-transduced reporter cells were used for all experiments conducted in this work. Specifically, healthy human lung fibroblasts (LL24 (RRID:CVCL_2575); CCL-151, male; ATCC, Manassas, VA) were transduced with a 'stable' alpha smooth muscle actin (αSMA) reporter plasmid, according to a previously reported protocol.[18] The reporter plasmid included two fluorescent reporting genes: (1) a red fluorescent protein, DsRed-Express2 (DsRed), expressed by the constitutive PGK promoter and (2) a green fluorescent protein, ZsGreen, conditionally expressed when the ACTA2 promoter is activated, where the ACTA2 promoter is associated with αSMA expression. The 1:1 ratio of red and green fluorescent reporting genes in each transduced cell allows for normalization of the variable green fluorescence intensity relative to the constitutive red fluorescence intensity.

Reporter lung fibroblasts were expanded on tissue culture treated polystyrene flasks (75 $cm^2$; CELLTREAT, Pepperell, MA) at 37 °C and 5% $CO_2$ under sterile conditions and using sterile technique. From thaw, fibroblasts were cultured in 10% FBS media: Ham's F12K media supplemented with 50 U mL$^{-1}$ penicillin, 50 μg mL$^{-1}$ streptomycin, 0.2% v:v amphotericin B/Fungizone, and 10% v:v FBS. During cell expansion, the media (10% FBS) was replaced every 2-3 days and cells were passaged upon reaching approximately 85% confluency.

*Hydrogel encapsulation of fibroblasts and 3D cell culture*

For cell encapsulation, hydrogel precursor solution was prepared in a protocol similar to that described above (*Hydrogel formation*), using sterile stock solutions and sterile DPBS. Specifically, the appropriate volumes of PEG-SH, LAP, linker peptide, 2 mM RGDS, and DPBS (accounting for the volume of DPBS that would later be provided from the cell solution) were combined and thoroughly vortex-mixed. Reporter fibroblasts were trypsinized for approximately 6 minutes (0.25% Trypsin/2.21mM EDTA solution; Thermo Fisher Scientific, Waltham, MA) after which 10% FBS media (described for cell expansion) was added to neutralize the trypsin. A sample of the cell suspension was removed for cell counting using a hemocytometer while the remaining cell suspension was centrifuged. The centrifuged cells were resuspended in DPBS at a cell density of $20 \times 10^6$ cells mL$^{-1}$. The hydrogel precursor solution was added to the appropriate volume of cell suspension and gently pipette-mixed (mixing the viscous precursor solution into the non-viscous cell suspension allows for better mixing) for a final cell density of $5 \times 10^6$ cells mL$^{-1}$ (passage 11-12). The hydrogel precursor solution with suspended fibroblasts was transferred to the dry mfCMP-Azide (previously assembled, aliquoted, lyophilized, and brought to room temperature) for a final hydrogel composition of PEG-SH (20 mM SH), 2.2 mM LAP, bis-alloc linker peptide (13 mM alloc), mono-alloc RGDS (2 mM alloc), and mono-alloc/mono-azide mfCMP-Azide (5 mM alloc, 5 mM azide), resulting in a final 1:1 ratio of thiol:alloc. The complete hydrogel precursor solution with suspended fibroblasts was transferred to syringe molds (20 μL) and irradiated with a cytocompatible dose of long wavelength UV light as described above for complete gelation.

The final hydrogels were transferred to a non-treated 48-well plate with 10% FBS media at 37 °C and 5% $CO_2$. To remove any unreacted hydrogel components, the media was replaced 30 minutes after encapsulation and again 2 hours later. The fibroblasts were cultured within the 3D hydrogels in 10% FBS media at 37 °C and 5% $CO_2$, where the media was replaced every 2 days (starting at 24 hours after encapsulation) over the course of 7 days to allow for the fibroblasts to become established in the hydrogel environment and transition from rounded to elongated morphologies similar to those seen in native environments. To minimize the impact of FBS on fibroblast activation during the experiment, the media was replaced with 5% FBS media after 7 days of 3D culture (Ham's F12K media supplemented with 50 U mL$^{-1}$ penicillin, 50 μg mL$^{-1}$ streptomycin, 0.2% v:v amphotericin B/Fungizone, and 5% v:v FBS) and 1% FBS media 24 hours later (Ham's F12K media supplemented with 50 U mL$^{-1}$ penicillin, 50 μg mL$^{-1}$ streptomycin, 0.2% v:v amphotericin B/Fungizone, and 1% v:v FBS). After 24 hours of culture with 1% FBS media, treatment with macrophage-conditioned media or with hydrogel stiffening was initiated. For the remainder of the experiment, 1% FBS media was used unless otherwise stated.

*Macrophage-conditioned media treatment*

The supernatant from MH-S macrophage cells (a murine alveolar macrophage cell line) was collected, centrifuged to remove any debris, and stored at -20 °C until use.[19] Prior to use, the supernatant was thawed, sterile filtered, and warmed to 37 °C. For encapsulated fibroblasts undergoing the MH-S-conditioned media treatment, the 1% FBS media was removed and replaced with 100% conditioned media (pure MH-S supernatant; 250 μL gel$^{-1}$) for 24 hours. After the first 24 hours, the media was replaced with 50% conditioned media (a 1:1 ratio of MH-S-supernatant:fresh 1% FBS media; 250 μL gel$^{-1}$), which was then replaced with new 50% conditioned media every 24 hours until the end of the experiment.

*Hydrogel stiffening treatment*

Hydrogel stiffening was conducted according to the protocol outlined above (*Stiffening protocol*) under sterile conditions. Each stiffening cycle consisted of an incubation in PEG-xBCN stiffening solution (approximately 12 hours) followed by an incubation in PEG-Azide stiffening solution (approximately 12 hours), where a total of 3 stiffening cycles were implemented over the course of 3 days. After the final cycle of stiffening, hydrogels were washed 3× for 1 hour each in warmed, fresh 1% FBS media to remove any remaining monomers, and the hydrogels were cultured in 1% FBS for an additional 2 days.

*Metabolic activity assay*

Fibroblast metabolic activity within each hydrogel was assessed on days -5, -1, 3, and 5 of the experiment (4, 8, 12, and 14 days after encapsulation, respectively). An alamarBlue assay (Thermo Fisher Scientific, Waltham, MA) was used to assess metabolic activity, according to the protocol provided by the manufacturer. Briefly, a solution of serum-free media (Ham's F12K media supplemented with 50 U $mL^{-1}$ penicillin, 50 µg $mL^{-1}$ streptomycin, 0.2% v:v amphotericin B/Fungizone) and alamarBlue was prepared (1:10 alamarBlue:serum-free media). Media solutions (day -5: expansion 10% FBS media; day -1: transition 5% FBS media; days 3/5: 1% FBS media, conditioned media, or stiffening media) were removed from each hydrogel and replaced with the alamarBlue working solution (400 µL $hydrogel^{-1}$), and the alamarBlue working solution was added to 3 wells of a new 48 well plate (400 µL $well^{-1}$) for background subtraction. Hydrogels were incubated in the solution for 4 hours at 37 °C.

alamarBlue solutions were removed and samples were plated in a 96-well plate (100 µL/well), and the fluorescence was measured on a plate reader (ex/em: 570/585; Spectramax i3X microplate reader; Molecular Devices, San Jose, CA). The average alamarBlue background reading was subtracted from each hydrogel output, and percent metabolic activity was determined for each hydrogel by setting the control hydrogels at 100% metabolic activity and a fluorescence of 0 RFU to be 0% metabolic activity. Each hydrogel was then internally normalized to the metabolic activity of that hydrogel at day -5.

After the alamarBlue solutions were removed from the hydrogels, the hydrogels were thoroughly washed with serum-free media (2× 30-minute incubations, 1× 60-minute incubation, and 1× 90-minute incubation) at 37 °C to remove the alamarBlue fluorophore prior to replacing the appropriate media or treatment condition. For each condition at each timepoint, metabolic activity was tested for n = 3 hydrogel-based 3D cultures.

*Immunostaining and imaging*

Encapsulated human lung fibroblasts were fixed in 4% methanol-free paraformaldehyde (PFA; Thermo Fisher Scientific, Waltham, MA) for 20 minutes after 14 total days of culture (including the 3D culture time and the experimental time). The hydrogels were then washed in DPBS and incubated overnight at 4°C in a bovine serum albumin (BSA) blocking solution (5% w:v BSA:DPBS; MilliporeSigma, Burlington, MA). All washes between labeling steps were performed in a 1.5% w:v BSA:DPBS solution containing 0.2% TWEEN-20 (MilliporeSigma, Burlington, MA). To permeabilize the fibroblasts, the hydrogels were treated with a 0.2% Triton X-100 solution (Thermo Fisher Scientific, Waltham, MA) for 30 minutes at room temperature. To immunostain for αSMA, hydrogels were incubated overnight at 4°C with mouse-anti-human αSMA antibody (ab7817, 1A4, 1:100; Abcam, Cambridge, UK). Next, the hydrogels were incubated with a fluorophore-conjugated goat-anti-mouse secondary antibody (Alexa Fluor 647, A21235, 1:250; Thermo Fisher Scientific; Waltham, MA) for 2 hours at room temperature, followed by an overnight incubation at 4°C. The hydrogels were then incubated with ActinRed ReadyProbe (40 µL mL$^{-1}$; Thermo Fisher Scientific, Waltham, MA) first for 2 hours at room temperature, followed by an overnight incubation at 4°C. Finally, hydrogels were incubated with Hoechst (4 µL mL$^{-1}$; Thermo Fisher Scientific, Waltham, MA) at room temperature for 20 minutes. The hydrogels were washed with DPBS and transferred to an 8-well chambered coverglass and covered with fresh DPBS. For each condition, n = 3 hydrogel-based 3D cultures were imaged.

Confocal microscopy (Zeiss LSM 800 confocal microscope) was used to capture *z*-stack images of the immunostained fibroblasts (3 gels per condition, 3 images per gel, 200 µm thick *z*-stacks). The resulting images were processed and analyzed according to the methods laid out below. Note that while the reporter cell lines were used here, much of the red fluorescent and green fluorescent proteins become inactive with a 20-minute fixation process, likely because the crosslinking during fixation interferes with the fluorophore structures. Any residual fluorescence from the DsRed protein will only overlap with the F-Actin staining, where F-Actin is used to identify the cell bodies, and any remaining DsRed will be located within the cell body. As no quantification of the red channel fluorescence intensity will be conducted on the fixed cells, residual DsRed fluorescence will not interfere with the results. The green channel was not used for imaging, so any residual ZsGreen protein will not interfere with other protein immunostaining detection.

*Live cell imaging*

Live cell imaging experiments were performed to assess changes in αSMA expression and fibroblast motility in response to either soluble factors or environmental stiffening over time, in real-time. The reporter proteins (DsRed and ZsGreen) were utilized for cell detection; specifically, the constitutive DsRed was used as a live cell-tracker protein to monitor cell motility over multiple time lapses, and DsRed and ZsGreen were monitored at specific time points to capture changes in αSMA expression throughout the experiment. Brightfield images were collected along with the red and green channels to allow for tracking of hydrogel drift during the timelapse. Timelapses were conducted at 2, 15, 37, and 65 hours from the time the experimental conditions were introduced (where time lapses ranged from 9 to 13 hours). For each condition at each timepoint, live imaging was performed on n = 3 hydrogel-based 3D cultures.

At each timepoint hydrogels were transferred to a Nunc Lab-Tek II 8 well chambered coverglass (Thermo Fisher Scientific, Waltham, MA) with the appropriate treatment media. Fluorescent images were captured every hour over the course of each timelapse using a Zeiss LSM 800 confocal microscope equipped with an incubation chamber (3 gels per condition, 3 images per gel, 150 µm thick *z*-stacks; incubation settings: 37 °C, 5% $CO_2$, with humidity control; Zeiss, Oberkochen, Germany).

##### 1.4 Image analysis methods

*Cell morphology and cell counts*

Using *z*-stack images of fixed, immunostained samples, cell bodies were detected using Volocity 3D image analysis software (PerkinElmer, Waltham, MA). Cell bodies were identified from the unprocessed images with the red channel (ActinRed-labeled F-actin), and the detected objects were filtered to improve measurement accuracy (Close filter, 4 iterations; Fill Holes in Objects filter; Remove Noise From Objects filter, fine; and Exclude Objects by Size filter, objects below 5000 µm$^3$ were excluded). Nuclei were then detected with the blue channel (Hoechst), with further filtering applied (Remove Noise From Objects filter, coarse; Exclude Objects by Size filter, objects below 1500 µm$^3$ were excluded; and Separate Touching Objects filter, object size guide 2500 µm$^3$). Some detected nuclei were not located within the identified cell objects (individual cells or multiple cells in contact); the cells belonging with these nuclei were classified as 'not intact at the time of fixation' and were excluded from the nuclei count using the Compartmentalize function, which divided the detected nuclei among the identified cell objects. The resulting Volocity data were exported and MATLAB (MathWorks, Natick, MA) was used to sort data and perform statistical analyses on shape factor (i.e., cell object sphericity) and intact cell count (i.e., number of nuclei located within cell objects) measurements. Three hydrogels were imaged per condition, with >20 intact cells counted per hydrogel (43-45 cells per hydrogel on average, depending on the condition), and a total of > 120 cells were analyzed for each condition.

*Protein fluorescence quantification*
Using the same unprocessed *z*-stack images of fixed, immunostained samples, Volocity was used to measure the Raw Integrated Density (RID, sum of the values of the pixels in the full image, across all z slices) of the blue (stained nuclei) and far-red (stained αSMA) channels. The total αSMA protein in the image correlated to the far-red channel RID, which was then normalized within each image to the blue channel RID (representing the total amount of nuclear content in the image). αSMA fluorescence was normalized to the nuclei fluorescence (including nuclei without an intact cell body) to avoid the αSMA associated with cell debris around the naked nuclei from artificially skewing the results. All conditions were then normalized to the control.

*Mature stress fiber quantification*
Maximum intensity orthogonal projections of the same *z*-stack images were used to visually assess the presence of stress fibers in each cell object previously identified in Volocity. Cell objects were considered positive for stress fiber formation if the object contained ≥3 distinct stress fibers. Stress fiber counts were performed manually on brightened maximum intensity orthogonal projections.

*αSMA reporter quantification*
Toward assessing the real-time impact of biochemical treatment and environmental stiffening on αSMA protein expression in fibroblasts, images from the live cell imaging experiment were analyzed at discrete timepoints (images at 2, 15, 37, and 65 hours after the start of experimental conditions, *i.e.*, the initial image from the corresponding timelapse). The resulting 3D images were then transformed into orthogonal projections using Fiji (ImageJ).[20] Specifically, the *Z Projection: sum slices* process was used to preserve the total fluorescence intensity information from 3D to 2D. In addition to the two detected channels (DsRed, constitutive reporter and ZsGreen, αSMA reporter) a third channel (blue) was created to assist in cell body detection during analysis. The blue "cell body" channel was the result of combining the DsRed and ZsGreen channels into a single (blue) channel such that cell body detection would take both reporters into account. All three channels were merged into a composite image and exported for analysis.

The resulting images were then analyzed using Imaris microscopy image analysis software (Oxford Instruments, Abingdon, UK). Cell objects were identified using the artificially created blue channel with the cell only detection type (smoothing function: 0.624 μm; no background subtraction; manual threshold; no splitting), and objects with an area below 50 $\mu m^2$ were filtered out. Once cell objects were identified, the sum intensity of each channel (DsRed and ZsGreen) were exported for each object, as well as the object area detected using the blue channel.

The exported Imaris data were then sorted using MATLAB, and any cell objects with areas below 200 $\mu m^2$ were filtered out. For each cell object, the normalized ZsGreen (αSMA) intensity was determined by calculating the ZsGreen sum intensity:DsRed sum intensity within each object, which was then weighted according to the cell object area. MATLAB was used to perform all statistical analyses. Three hydrogels were imaged per condition, with ≥48 cell objects (ranging from individual cells to clusters of multiple cells) analyzed per hydrogel (for each condition and timepoint, with an average of 65-70 objects per condition across all timepoints).

*Cell motility quantification*

Using the 3D *z*-stack timelapse images from the live imaging experiments, cell bodies were detected and tracked using Volocity. Cell bodies were identified from the unprocessed images with the red channel (DsRed constitutive reporter protein), and the detected objects were further processed using Dilate (3 iterations) and Erode (3 iterations) to remove some of the noise around the cells. The cell objects were then filtered to improve measurement accuracy: (1) Remove Noise From Objects filter (fine) and (2) Exclude Objects by Size filter (objects below 2000 $\mu m^3$ were excluded); Finally, the Track function (Shortest Path, 50 μm maximum distance between objects) was used to track each cell object over the course of the timelapse. The results of each image set were examined to ensure that any broken tracks were manually grouped together, and the cell objects were accurately tracked. Background hydrogel drift was determined by tracking a stationary non-cell object throughout the timelapse using the brightfield 3D image. Specifically, an object visible in brightfield was chosen, and at each timepoint within the timelapse the *z* location was adjusted to bring the object into focus, and a track point was manually placed at the center of the object. The track point location (*x*,*y*,*z*) was then tracked through the entire timelapse with the same method as tracking cells. This process was repeated for each image within each timelapse period.

The resulting Volocity data were exported, and MATLAB was used to determine the speed and directional persistence of each identified cell object. For each timelapse, the data were filtered to remove any datapoints classified either as tracks or cell objects detected for less than 3 hours (to limit bias from cells tracked for short periods of time) and were then sorted into conditions. The background hydrogel movement was then removed; specifically, for each image, the location (*x*,*y*,*z*) of the stationary non-cell object in each frame was subtracted from the center point (*x*,*y*,*z*) of all cell objects in the corresponding frame. For each cell object, the displacement (*d*, distance between the cell object in the first and last frames that object was detected) and total distance traveled (*D*, sum of the displacements between each frame) were measured. Rate of movement (speed) was then calculated for each cell object (*D* divided by the time over which the object was tracked), as well as the directionality ratio (calculated as *d*:*D*), where values closer to 0 are indicative of meandering cell movement (low directional persistence), and values closer to 1 are indicative of more direct cell movement (high directional persistence).[21]

#### 1.5 Mathematical modeling methods

*Mathematical modeling of reaction-diffusion in a cylindrical hydrogel*

The analytical solution of the PDE modeling the reaction-diffusion of conditioned media in the hydrogel was evaluated using finite transforms (Hankel, Fourier). The numerical solution of the system of PDEs modeling the reaction-diffusion of PEG monomers in the hydrogel was evaluated using the finite difference method. To maintain numerical accuracy, the system of equations modeling PEG was solved with second order central and forward difference schemes. The resulting solutions were modeled in MATLAB (MathWorks, Natick, MA) at discrete *r*- and *z*-locations. The numerical PEG solution was modeled with 1 second timesteps. To conserve computing time and power, the solute concentration (either conditioned media or PEG) in the surrounding solution was treated as a step function that was updated at each hour timepoint. Specifically, at each timepoint the amount of solute that had diffused into the hydrogel was calculated and subtracted from the amount of solute in the surrounding solution. The updated solution concentration was then used for the following hour of the model. Parameter values used in the model were either found in literature or measured for the materials in this work (**Table S7**).

#### 1.6 Statistical methods

*Statistical analysis*

All reported values are presented as the mean ± standard error for each condition, based on measurements from three independent samples ($n = 3$) unless otherwise specified. Statistical significance was assessed using a one-way ANOVA followed by Tukey's post-hoc test. In the plots, statistical significance is indicated as either not statistically different (n.s.) or as significantly different (* $p < 0.05$; ** $p < 0.01$; *** $p < 0.001$; **** $p < 0.0001$).

#### 2. List of Symbols

| | |
|---|---|
| $k_{side}$ | Thiol–yne reaction rate constant ($M^{-1}$ $s^{-1}$) |
| $r_{side}$ | Thiol–yne reaction rate (M $s^{-1}$) |
| $C_{SH}$ | Thiol concentration (M) |
| $C_{Oct}$ | Cyclooctyne concentration (M) |
| $\xi_r$ | Extent of reaction |
| $t$ | Time (s) |
| $C_{Oct,0}$ | Initial cyclooctyne concentration (M) |
| $\mathbf{J}$ | Total flux (mol $m^{-2}$ $s^{-1}$) |
| $\mathbf{J}_r$ | Flux in the radial direction (mol $m^{-2}$ $s^{-1}$) |
| $\mathbf{J}_\theta$ | Flux in the azimuthal direction (mol $m^{-2}$ $s^{-1}$) |
| $\mathbf{J}_z$ | Flux in the vertical direction (mol $m^{-2}$ $s^{-1}$) |
| $r, \theta, z$ | Cylindrical coordinates |
| $\hat{\mathbf{r}}, \hat{\boldsymbol{\theta}}, \hat{\mathbf{z}}$ | Cylindrical coordinate unit vectors |
| $D_{eff}$ | Effective diffusivity ($m^2$ $s^{-1}$) |
| $C$ | Species concentration (M) |
| $r_{cm}$ | Rate of conditioned media degradation/inactivation |
| $k$ | Reaction rate constant (first order: $s^{-1}$; second order: $M^{-1}$ $s^{-1}$) |
| $C_\infty$ | Species concentration in solution (M; assumed to be constant in space throughout the solution) |
| $R$ | Hydrogel radius (m) |
| $H$ | Hydrogel height (m) |
| $u(r,z,t)$ | Conditioned media concentration after change of variables (M) |
| $Y_m(r)$ | Finite Hankel transform (FHT) eigenfunction |
| $J_0$ | Zero-order Bessel function |
| $\mu_m$ | FHT eigenvalues |
| $j_{0,m}$ | Positive roots of the zero-order Bessel function |
| $H_0\{u\}$ | Zero-order FHT operator |
| $\bar{u}_m(z,t)$ | FHT-transformed function $u(r,z,t)$ |
| $X_n(z)$ | Finite Fourier transform (FFT) eigenfunction |
| $\lambda_n$ | FFT eigenvalues |
| $F\{\bar{u}_m\}$ | FFT operator |
| $\bar{\bar{u}}_{m,n}(t)$ | FFT-transformed function $\bar{u}_m(z,t)$ |
| $J_1$ | First-order Bessel function |
| $A_1$ | Constant of integration |
| $A_{m,n}$ | Constant |
| $\mathrm{n}_R$ | Number of moles of solute (mol) through the top of the hydrogel |
| $\mathrm{n}_H$ | Number of moles of solute (mol) through the side of the hydrogel |
| $\mathrm{n}_T$ | Number of moles of solute (mol) through the exterior of the hydrogel |
| $V_S$ | Solution volume (L) |
| $r_1$ | Reaction rate 1 (M $s^{-1}$) |
| $r_2$ | Reaction rate 2 (M $s^{-1}$) |
| A | Species A: functional group fixed to the hydrogel |
| B | Species B: functional group attached to diffusing PEG |
| X | Species X: covalent linkage formed between A and B |

| | |
|---|---|
| $P_0$ | Species $P_0$: unreacted, freely diffusing PEG |
| $P_F$ | Species $P_F$: PEG fixed to hydrogel after functional group reaction |
| $f$ | Number of "available" functional groups per PEG molecule (determined according to PEG, $P_0$, functional group molar excess, i.e., $f = 4A/B$) |
| P | Species P: total PEG in hydrogel (diffusing + fixed) |
| $\Delta r$ | Discrete length in the $r$-direction (m) |
| $M$ | Number of nodes in the $r$-direction |
| $\Delta z$ | Discrete length in the $z$-direction (m) |
| $N$ | Number of nodes in the $z$-direction |
| $r_i$ | Distance from axis of symmetry at node $i$ (m) |
| $\varepsilon$ | Error term |
| $n_t$ | Total number of moles of PEG (free and fixed) within the hydrogel (mol) at the end of timestep t |
| $C_0$ | Initial PEG concentration in solution (M) |
| $\xi$ | Hydrogel mesh size (nm) |
| $\nu_r$ | Hydrogel polymer volume fraction (relaxed state) |
| $\nu_s$ | Hydrogel polymer volume fraction (equilibrium swollen state) |
| $Q_r$ | Hydrogel swelling ratio (relaxed state) |
| $Q_s$ | Hydrogel swelling ratio (equilibrium swollen state) |
| $\ell$ | Bond length along PEG backbone (nm) |
| $C_n$ | Flory characteristic ratio of PEG |
| $\overline{M_c}$ | Molecular weight between crosslinks (g mol$^{-1}$) |
| $M_r$ | PEG repeat unit molecular weight (g mol$^{-1}$) |
| $G'$ | Storage modulus (Pa) |
| $\rho$ | Density of PEG (g mL$^{-1}$) |
| $T$ | Temperature (K) |
| $\overline{M_n}$ | Total monomer average molecular weight (g mol$^{-1}$) |
| $\rho_x$ | Hydrogel crosslink density (mM) |
| $D_\infty$ | Diffusivity in water (cm$^2$ s$^{-1}$) |
| $k_B$ | Boltzmann constant (J/K) |
| $\eta$ | Dynamic viscosity (kg m$^{-1}$ s$^{-1}$) |
| $r_h$ | Hydrodynamic radius (nm) |
| $r_f$ | Polymer chain radius (nm) |

#### 3. Supplemental Calculations

*Side thiol–cyclooctyne reaction rate vs SPAAC reaction rate*

As with any system, one must consider the potential for side reactions that could interfere with the desired reaction. In the SPAAC stiffening approach, there is the possibility for side reaction of the cyclooctyne with free thiols (present in the media). [22-23] Toward assessing the reaction rate constant of side thiol–yne reactions, and the resulting reaction rate compared to that of the SPAAC reaction, we approximated the reaction rate constant, $k_{side}$, for the thiol–yne reaction. Using the results from the reported spontaneous reaction between thiol and cyclooctyne,[22] we calculated the $k_{side}$ value for each datapoint. Briefly, assuming a second order reaction (thiol + octyne $\xrightarrow{r_{side}}$ side product) and that the thiol and octyne concentrations are the same ($C_{SH} = C_{Oct}$), the reported extent of reaction values were converted to concentration:

$$C_{Oct} = 2\text{M} - (2\text{M})(\xi_r) \tag{S1}$$

where the initial octyne (and thiol) concentration was 2M. The reaction rate, concentration, and reaction rate constant were calculated as follows:

$$\frac{dC_{Oct}}{dt} = -k_{side}C_{SH}C_{Oct} = -k_{side}{C_{Oct}}^2 \tag{S2}$$

$$C_{Oct} = \frac{C_{Oct,0}}{1 + C_{Oct,0}k_{side}t} \tag{S3}$$

$$k_{side} = \frac{C_{Oct,0} - C_{Oct}}{C_{Oct,0}C_{Oct}t} \tag{S4}$$

The value of $k_{side}$ was calculated for each datapoint using **Equation S4**, and the overall $k_{side}$ approximation was determined by averaging the resulting values ($k_{side} = 0.015\ \text{M}^{-1}\ \text{s}^{-1}$). In comparison, the rate constant for a SPAAC reaction using a BCN functional group is an order of magnitude higher ($k_{BCN-Azide} = 0.14\ \text{M}^{-1}\ \text{s}^{-1}$),[23] which increases with pegylated functional groups reacted under physiological conditions ($k_{BCN-Azide} = 0.57\ \text{M}^{-1}\ \text{s}^{-1}$; 37 °C in human blood plasma).[24-25]

For this work, Ham's F-12K (Kaighn's) media was used (contains 0.40 mM L-cysteine and 0 mM glutathione).[26] Neglecting any thiol from added FBS, we calculated the thiol–yne side reaction rate to be 0.018 μM $\text{s}^{-1}$, which was two orders of magnitude lower than the anticipated SPAAC reaction rate of 1.3 μM $\text{s}^{-1}$. Notably, this calculation did not account for thiols present in FBS (due to the not well-defined nature of the product); therefore, all stiffening was conducted with 1% FBS media (rather than the standard 10%) to minimize thiol content in addition to limiting FBS-driven fibroblast activation without serum-starving the cells.

*Flux equations for a diffusing species*

Total flux and one-dimensional fluxes for a 2D axisymmetric cylindrical system are as follows:

$$\mathbf{J} = -D_{eff}\nabla C\ , \tag{S5}$$

$$\mathbf{J}_r = -D_{eff}\frac{\partial C}{\partial r}\hat{\mathbf{r}}\ , \tag{S6}$$

$$\mathbf{J}_\theta = -D_{eff}\frac{1}{r}\frac{\partial C}{\partial \theta}\hat{\boldsymbol{\theta}} = 0\ \text{, and} \tag{S7}$$

$$\mathbf{J}_z = -D_{eff}\frac{\partial C}{\partial z}\hat{\mathbf{z}}\,. \tag{S8}$$

*Conditioned media concentration within a hydrogel (analytical solution)*

The concentration of conditioned media within a hydrogel was solved analytically. Given the reaction rate

$$r_{cm} = -kC\,, \tag{S9}$$

the continuity equation was defined as follows:

$$\frac{\partial C}{\partial t} = D_{eff}\nabla^2 C + r_{cm} \tag{S10}$$

$$\frac{\partial C}{\partial t} = D_{eff}\left[\frac{1}{r}\frac{\partial}{\partial r}\left(r\frac{\partial C}{\partial r}\right) + \frac{1}{r^2}\frac{\partial^2 C}{\partial \theta^2} + \frac{\partial^2 C}{\partial z^2}\right] - kC \tag{S11}$$

$$\frac{\partial C}{\partial t} = D_{eff}\left[\frac{\partial^2 C}{\partial r^2} + \frac{1}{r}\frac{\partial C}{\partial r} + \frac{\partial^2 C}{\partial z^2}\right] - kC\,, \tag{S12}$$

where $C$ is a function of $r, z, t$. Here, the reaction-diffusion equation has been reduced, as there is no theta dependence within this system. The following boundary/axis of symmetry conditions and initial conditions apply to this system:

$$C(r = R) = C_\infty \tag{S13}$$

$$C(z = H) = C_\infty \tag{S14}$$

$$\left.\frac{\partial C}{\partial r}\right|_{r=0} = 0 \tag{S15}$$

$$\left.\frac{\partial C}{\partial z}\right|_{z=0} = 0 \tag{S16}$$

$$C(r, z, t = 0) = 0 \tag{S17}$$

Importantly, the concentration of the surrounding solution ($C_\infty$) was treated as a constant for the purpose of simplicity. As the hydrogel and surrounding solution are a closed system, $C_\infty$ will decrease as the protein diffuses into the hydrogel. This was later accounted for in the computer model by calculating molar flux into the hydrogel and updating $C_\infty$ at discrete timepoints.

Notably, this system contains non-homogeneous boundary conditions, requiring the application of a change of variables.

$$u(r, z, t) = C(r, z, t) - C_\infty \iff C(r, z, t) = u(r, z, t) + C_\infty \tag{S18}$$

$$\frac{\partial u}{\partial t} = D_{eff}\left[\frac{\partial^2 u}{\partial r^2} + \frac{1}{r}\frac{\partial u}{\partial r} + \frac{\partial^2 u}{\partial z^2}\right] - k(u + C_\infty) \tag{S19}$$

After the change of variables, the boundary conditions become:

$$u(r = R) = 0 \tag{S20}$$

$$u(z = H) = 0 \tag{S21}$$

$$\left.\frac{\partial u}{\partial r}\right|_{r=0} = 0 \tag{S22}$$

$$\left.\frac{\partial u}{\partial z}\right|_{z=0} = 0 \tag{S23}$$

Toward solving this system, we first applied a Finite Hankel Transform (FHT) according to the following equations valid for Dirichlet boundary conditions and a zero-order transform:

$$Y_m(r) = J_0(\mu_m r) \tag{S24}$$

$$J_0(\mu_m R) = 0 \tag{S25}$$

$$\mu_m = \frac{j_{0,m}}{R} \quad m = 1, 2, \dots \tag{S26}$$

$$\|Y_m(r)\|^2 = \frac{R^2}{2} {J_1}^2(\mu_m R) \tag{S27}$$

$$\overline{w}_m = H_0\{w(r)\} = \int_0^R w(r) J_0(\mu_m r) r dr \tag{S28}$$

$$H_0\left\{\frac{\partial^2 w(r)}{\partial r^2} + \frac{1}{r}\frac{\partial w(r)}{\partial r}\right\} = R\mu_m J_1(\mu_m R) w(R) - {\mu_m}^2 \overline{w}_m \tag{S29}$$

Applying the FHT to our system, we find:

$$H_0\left\{\frac{\partial u}{\partial t}\right\} = D_{eff} H_0\left\{\frac{\partial^2 u}{\partial r^2} + \frac{1}{r}\frac{\partial u}{\partial r}\right\} + D_{eff} H_0\left\{\frac{\partial^2 u}{\partial z^2}\right\} - H_0\{ku\} - H_0\{kC_\infty\} \tag{S30}$$

$$\frac{\partial \bar{u}_m}{\partial t} = D_{eff}\frac{\partial^2 \bar{u}_m}{\partial z^2} - \left[D_{eff}{\mu_m}^2 + k\right]\bar{u}_m - \frac{kC_\infty R J_1(\mu_m R)}{\mu_m} \tag{S31}$$

with the transformed remaining boundary conditions:

$$\bar{u}_m(z = H) = 0 \tag{S32}$$

$$\left.\frac{\partial \bar{u}_m}{\partial z}\right|_{z=0} = 0 \tag{S33}$$

We next applied a Finite Fourier Transform (FFT) according to the following equations valid for Neumann boundary conditions at $z = 0$ and Dirichlet boundary conditions at $z = H$:

$$X_n(z) = \cos(\lambda_n z) \tag{S34}$$

$$\cos(\lambda_n z) = 0 \tag{S35}$$

$$\lambda_n = \frac{2\pi n - \pi}{2H} \quad n = 1, 2, \dots \tag{S36}$$

$$\|X_n(z)\|^2 = \frac{H}{2} \tag{S37}$$

$$\overline{w}_n = F\{w(z)\} = \int_0^H w(z)\cos(\lambda_n z)\,dz \tag{S38}$$

$$F\left\{\frac{\partial^2 w(z)}{\partial z^2}\right\} = w(H)\lambda_n \sin(\lambda_n H) - \left.\frac{\partial w}{\partial z}\right|_{z=0}\cos(0) - {\lambda_n}^2\overline{w}_n \tag{S39}$$

Applying the FFT to our system, we find:

$$F\left\{\frac{\partial \overline{u}_m}{\partial t}\right\} = D_{eff}F\left\{\frac{\partial^2 \overline{u}_m}{\partial z^2}\right\} - \left[D_{eff}{\mu_m}^2 + k\right]F\{\overline{u}_m\} - \frac{kC_\infty R J_1(\mu_m R)}{\mu_m} \tag{S40}$$

$$\frac{\partial \overline{\overline{u}}_{m,n}}{\partial t} = -\left[D_{eff}{\mu_m}^2 + D_{eff}{\lambda_n}^2 + k\right]\overline{\overline{u}}_{m,n} - \frac{kC_\infty R J_1(\mu_m R)\sin(\lambda_n H)}{\mu_m \lambda_n} \tag{S41}$$

From here we solved the separable equation for $\overline{\overline{u}}_{m,n}$

$$\overline{\overline{u}}_{m,n}(t) = A_1 e^{-t\left[D_{eff}{\mu_m}^2 + D_{eff}{\lambda_n}^2 + k\right]} - \frac{kC_\infty R J_1(\mu_m R)\sin(\lambda_n H)}{\mu_m \lambda_n\left[D_{eff}{\mu_m}^2 + D_{eff}{\lambda_n}^2 + k\right]}, \tag{S42}$$

applied the inverse FFT

$$\overline{u}_m(z,t) = \sum_{n=1}^{\infty} \overline{\overline{u}}_{m,n}(t)\cos(\lambda_n z)\frac{2}{H}, \tag{S43}$$

and applied the inverse FHT

$$\begin{aligned} u(r,z,t) &= \sum_{m=1}^{\infty} \overline{u}_m(z,t)\, J_0(\mu_m r)\frac{2}{R^2 {J_1}^2(\mu_m R)} \\ &= \sum_{m=1}^{\infty}\sum_{n=1}^{\infty} \overline{\overline{u}}_{m,n}(t)\, J_0(\mu_m r)\cos(\lambda_n z)\frac{4}{HR^2 {J_1}^2(\mu_m R)} \end{aligned} \tag{S44}$$

$$\begin{aligned} u(r,z,t) = \sum_{m=1}^{\infty}\sum_{n=1}^{\infty} \Bigg( & A_{m,n} e^{-t\left[D_{eff}{\mu_m}^2 + D_{eff}{\lambda_n}^2 + k\right]} \\ & - \frac{4kC_\infty \sin(\lambda_n H)}{HR\mu_m \lambda_n J_1(\mu_m R)\left[D_{eff}{\mu_m}^2 + D_{eff}{\lambda_n}^2 + k\right]}\Bigg) J_0(\mu_m r)\cos(\lambda_n z) \end{aligned} \tag{S45}$$

Subsequently, we reversed the change of variables

$$\begin{aligned} C(r,z,t) = C_\infty & \\ + \sum_{m=1}^{\infty}\sum_{n=1}^{\infty} \Bigg( & A_{m,n} e^{-t\left[D_{eff}{\mu_m}^2 + D_{eff}{\lambda_n}^2 + k\right]} \\ & - \frac{4kC_\infty \sin(\lambda_n H)}{HR\mu_m \lambda_n J_1(\mu_m R)\left[D_{eff}{\mu_m}^2 + D_{eff}{\lambda_n}^2 + k\right]}\Bigg) J_0(\mu_m r)\cos(\lambda_n z) \end{aligned} \tag{S46}$$

and plugged the initial conditions into the concentration equation (**Equation S46**):

$$C(r,z,t=0) = 0 = C_\infty + \sum_{m=1}^{\infty}\sum_{n=1}^{\infty}\left(A_{m,n} - \frac{4kC_\infty \sin(\lambda_n H)}{HR\mu_m\lambda_n J_1(\mu_m R)\left[D_{eff}{\mu_m}^2 + D_{eff}{\lambda_n}^2 + k\right]}\right)J_0(\mu_m r)\cos(\lambda_n z) \tag{S47}$$

We then used orthogonality to solve for the constant, $A_{m,n}$. Specifically, we multiplied both sides of **Equation S47** by $J_0\left(\frac{r}{R}j_{0,m}\right)rdr$ and integrated over *r* from 0 to *R*, then multiplied both sides of the resulting equation by $\cos(\lambda_n z)\,dz$ and integrated over *z* from 0 to *H*:

$$A_{m,n} = -\frac{4C_\infty D_{eff}\left({\mu_m}^2 + {\lambda_n}^2\right)\sin(\lambda_n H)}{HR\mu_m\lambda_n J_1(\mu_m R)\left[D_{eff}{\mu_m}^2 + D_{eff}{\lambda_n}^2 + k\right]} \tag{S48}$$

Combining **Equation S46** with **Equation S48**, we achieved the analytical solution to the concentration of conditioned media throughout the hydrogel:

$$C(r,z,t) = C_\infty - \sum_{m=1}^{\infty}\sum_{n=1}^{\infty}\frac{4C_\infty \sin(\lambda_n H)}{HR\mu_m\lambda_n J_1(\mu_m R)\left(D_{eff}{\mu_m}^2 + D_{eff}{\lambda_n}^2 + k\right)}\left[D_{eff}\left({\mu_m}^2 + {\lambda_n}^2\right)e^{-t\left(D_{eff}{\mu_m}^2 + D_{eff}{\lambda_n}^2 + k\right)} + k\right]J_0(\mu_m r)\cos(\lambda_n z) \tag{S49}$$

*MATLAB model of conditioned media concentration within a hydrogel with a changing outer boundary condition*

Revisiting the fact that, in reality, $C_\infty$ would constantly be changing as conditioned media diffuses from outside to inside the hydrogel, the concentration equation, $C(r,z,t)$, was modified to recalculate constant $A_{m,n}$ as $C_\infty$ updates in a stepwise fashion. Briefly, the flux at the hydrogel-solution interface was calculated over the course of each 1-hour timestep, $C_\infty$, was updated, and $A_{m,n}$ was accordingly updated.

In detail, the concentration equation (**Equation S46**) was modified for each timestep, simplifying the constants as follows:

$$C_i(r,z,t) = C_{\infty,i} + \sum_{m=1}^{\infty}\sum_{n=1}^{\infty}\left(A_i e^{-t\left[D_{eff}{\mu_m}^2 + D_{eff}{\lambda_n}^2 + k\right]} - B_i\right)J_0(\mu_m r)\cos(\lambda_n z)\,, \tag{S50}$$

where

$$B_i = \frac{4kC_{\infty,i}\sin(\lambda_n H)}{HR\mu_m\lambda_n J_1(\mu_m R)\left[D_{eff}{\mu_m}^2 + D_{eff}{\lambda_n}^2 + k\right]}\,. \tag{S51}$$

With the knowledge that $C_{i-1}(r,z,t_f)$ from the previous step would then become the new initial conditions, $C_i(r,z,t=0)$, for the current timestep, we established the following relationship:

$$C_i(r,z,t=0) = C_{i-1}(r,z,t_f) = C_{\infty,i-1} + \sum_{m=1}^{\infty}\sum_{n=1}^{\infty}(\gamma_{i-1})J_0(\mu_m r)\cos(\lambda_n z)\,, \tag{S52}$$

where the previous time function at $t_f$ was simplified as

$$\gamma_{i-1} = A_{i-1} e^{-t_f[D_{eff}\mu_m{}^2 + D_{eff}\lambda_n{}^2 + k]} - B_{i-1} \,. \tag{S53}$$

Relating **Equation S50** and **S52**,

$$\begin{aligned} C_i(r,z,t=0) &= C_{\infty,i} + \sum_{m=1}^{\infty}\sum_{n=1}^{\infty}(A_i - B_i) J_0(\mu_m r)\cos(\lambda_n z) \\ &= C_{\infty,i-1} + \sum_{m=1}^{\infty}\sum_{n=1}^{\infty}(\gamma_{i-1}) J_0(\mu_m r)\cos(\lambda_n z) \,. \end{aligned} \tag{S54}$$

Once again using orthogonality principals to solve for constant $A_i$, we found that

$$A_i = \frac{4\sin(\lambda_n H)}{HR\mu_m\lambda_n J_1(\mu_m R)}\left(C_{\infty,i-1} - C_{\infty,i}\right) + B_i + \gamma_{i-1} \,, \tag{S55}$$

where $C_{\infty,i} = C_\infty$ and $A_i = A_{m,n}$ for the initial timestep.

In order to update $C_\infty$ at the end of each 1-hour time step, the molar flux through the top and side of the hydrogel was determined:

$$\mathrm{J}_r(r=R) = \sum_{m=1}^{\infty}\sum_{n=1}^{\infty} D_{eff}\,\mu_m\,J_1(\mu_m R)\cos(\lambda_n z)\left(A_i e^{-t[D_{eff}\mu_m{}^2 + D_{eff}\lambda_n{}^2 + k]} - B_i\right) \tag{S56}$$

$$\mathrm{J}_z(z=H) = \sum_{m=1}^{\infty}\sum_{n=1}^{\infty} D_{eff}\,\lambda_n\ \sin(\lambda_n H)\,J_0(\mu_m r)\left(A_i e^{-t[D_{eff}\mu_m{}^2 + D_{eff}\lambda_n{}^2 + k]} - B_i\right) \tag{S57}$$

The rate of protein transport across the respective interfaces was calculated by integrating over the appropriate surface:

$$\begin{aligned} \frac{d\mathrm{n}_R}{dt} &= \int_0^{2\pi}\int_0^{H} R\,\mathrm{J}_r(r=R)\,dz\,d\theta \\ &= \sum_{m=1}^{\infty}\sum_{n=1}^{\infty} 2\pi R D_{eff} J_1(\mu_m R)\sin(\lambda_n H)\frac{\mu_m}{\lambda_n}\Big(A_i e^{-t[D_{eff}\mu_m{}^2 + D_{eff}\lambda_n{}^2 + k]} \\ &\quad - B_i\Big) \end{aligned} \tag{S58}$$

$$\begin{aligned} \frac{d\mathrm{n}_H}{dt} &= \int_0^{2\pi}\int_0^{R} \mathrm{J}_z(z=H)\,r\,dr\,d\theta \\ &= \sum_{m=1}^{\infty}\sum_{n=1}^{\infty} 2\pi R D_{eff} J_1(\mu_m R)\sin(\lambda_n H)\frac{\lambda_n}{\mu_m}\Big(A_i e^{-t[D_{eff}\mu_m{}^2 + D_{eff}\lambda_n{}^2 + k]} \\ &\quad - B_i\Big) \end{aligned} \tag{S59}$$

Adding the two rates and integrating over time,

$$\mathrm{n}_T = \mathrm{n}_R + \mathrm{n}_H \longrightarrow \frac{d\mathrm{n}_T}{dt} = \frac{d\mathrm{n}_R}{dt} + \frac{d\mathrm{n}_H}{dt} \tag{S60}$$

$$\begin{aligned} \frac{d\mathrm{n}_T}{dt} &= \sum_{m=1}^{\infty}\sum_{n=1}^{\infty}\frac{2\pi R D_{eff} J_1(\mu_m R)\sin(\lambda_n H)\left(\mu_m{}^2 + \lambda_n{}^2\right)}{\mu_m\lambda_n}\Big(A_i e^{-t[D_{eff}\mu_m{}^2 + D_{eff}\lambda_n{}^2 + k]} \\ &\quad - B_i\Big) \end{aligned} \tag{S61}$$

$$n_T = \sum_{m=1}^{\infty}\sum_{n=1}^{\infty} \frac{2\pi R D_{eff} J_1(\mu_m R)\sin(\lambda_n H)\left({\mu_m}^2 + {\lambda_n}^2\right)}{\mu_m \lambda_n} \int_0^{t_f} \left(A_i e^{-t\left[D_{eff}{\mu_m}^2 + D_{eff}{\lambda_n}^2 + k\right]} - B_i\right) dt \tag{S62}$$

$$n_T = \sum_{m=1}^{\infty}\sum_{n=1}^{\infty} \left[\frac{A_i\left(1 - e^{-t_f\left[D_{eff}{\mu_m}^2 + D_{eff}{\lambda_n}^2 + k\right]}\right)}{D_{eff}{\mu_m}^2 + D_{eff}{\lambda_n}^2 + k} - B_i t_f\right] \frac{2\pi R D_{eff} J_1(\mu_m R)\sin(\lambda_n H)\left({\mu_m}^2 + {\lambda_n}^2\right)}{\mu_m \lambda_n} \tag{S63}$$

where $n_T$ is the total number of moles to cross the hydrogel-solution interface during the 1-hour timestep, $t_f = 1$ hour. Finally, the concentration for the next timestep becomes:

$$C_{\infty,i+1} = \frac{C_{\infty,i} V_S + n_T}{V_S}, \tag{S64}$$

and constants $A_{i+1}$ and $B_{i+1}$ should be calculated accordingly. For simplicity, the conditioned media was assumed to undergo no degradation (i.e., $k = 0$) outside of the hydrogel. The MATLAB model was run for the equivalent of the 72-hour cell experiment, where the CM concentration was represented as a percent to reflect the nomenclature used in the experiment ($C_\infty = 100\%$ at $t = 0$, i.e., $C_{\infty,1} = 100\%$). Similarly, the solution concentration was reset to $C_\infty = 50\%$ after 24 and 48 hours (i.e., $C_{\infty,25} = C_{\infty,49} = 50\%$) to represent the media replacement at the end of days 1 and 2.

*PEG concentrations (unreacted and reacted) within a hydrogel (numerical solution)*

For a functionalized PEG diffusing into and reacting with the hydrogel, diffusion of PEG had to be considered, along with the reaction rates of individual functional groups. Two reaction rates were established, $r_1$ (reaction of functional groups to form a covalent linkage) and $r_2$ (transition of untethered, freely diffusing PEG to fixed PEG after forming linkages with the hydrogel). Reaction 1 is described as follows:

$$\mathrm{A} + \mathrm{B} \xrightarrow{r_1} \mathrm{X} \tag{S65}$$

where A is the functional group fixed to the hydrogel, B is functional group on diffusing PEG, X is the resulting covalent linkage. The resulting reaction rate can be described as

$$r_1 = kC_A C_B = kC_A C_{P0} f \tag{S66}$$

where, in terms of the freely diffusing PEG ($P_0$), $f$ describes the number of “available” functional groups per PEG molecule (determined according to the PEG functional group molar excess at a theoretical diffusion equilibrium throughout the system volume, i.e., $f = 4 \times$ [available functional group on hydrogel prior to incubation]/[stiffening PEG functional group in solution at equilibrium]; see **Table S2**). The rate constant, k, describes the SPAAC reaction between an azide and a BCN group under physiologically relevant conditions. Reaction 2 is described as follows:

$$\mathrm{P_0} \xrightarrow{r_2} \mathrm{P_F} \tag{S67}$$

where $P_0$ is unreacted and freely diffusing (not fixed) PEG and $P_F$ is fixed PEG after functional group reaction with the hydrogel to establish new linkages. The resulting reaction rate can be described as

$$r_2 = \frac{1}{f} r_1 = k C_A C_{P0} \tag{S68}$$

where we made the assumption that all "available" functional groups (B) on the free PEG ($P_0$) must react for the PEG to become fixed ($P_F$). While this is not an accurate representation of free PEG becoming fixed to the network, this assumption allowed us to proceed with the modeling without implementing a stochastic model. Specifically, in a physically accurate system, PEG monomers would become fixed to the hydrogel with a single functional group reaction, which would then change the probability of reaction for the remaining unreacted functional groups. In order to model such a system, one would require implementation of a stochastic model, which was beyond the scope of this work.

As the functional group A and covalent linkage X are fixed to the hydrogel, and thus unable to diffuse, the concentration equations for these two species will depend solely on the reaction rate:

$$\frac{\partial C_A}{\partial t} = -r_1 = -k C_A C_{P0} f \tag{S69}$$

$$\frac{\partial C_X}{\partial t} = r_1 = k C_A C_{P0} f \tag{S70}$$

The diffusion of PEG through the hydrogel will depend on the concentration gradient of free PEG, but also that of the fixed PEG, as both will contribute to the chemical potential gradient. We therefore created an additional variable to represent the total PEG, fixed and free (P):

$$C_P = C_{P0} + C_{PF} \tag{S71}$$

The free PEG diffuses according to the total PEG concentration ($C_P$), and reacts to become fixed according to the equation:

$$\begin{aligned} \frac{\partial C_{P0}}{\partial t} &= D_{eff}\left[\frac{\partial^2 C_P}{\partial r^2} + \frac{1}{r}\frac{\partial C_P}{\partial r} + \frac{\partial^2 C_P}{\partial z^2}\right] - r_2 \\ &= D_{eff}\left[\frac{\partial^2 C_P}{\partial r^2} + \frac{1}{r}\frac{\partial C_P}{\partial r} + \frac{\partial^2 C_P}{\partial z^2}\right] - k C_A C_{P0} \end{aligned} \tag{S72}$$

Comparatively, fixed PEG does not experience diffusion, and the concentration equation depends solely on the reaction rate:

$$\frac{\partial C_{PF}}{\partial t} = r_2 = k C_A C_{P0} \tag{S73}$$

All concentrations are functions of time and space $(r, z, t)$. Boundary/axis of symmetry conditions for this system are defined as

$$C_{P0}(r = R) = C_\infty \tag{S74}$$

$$C_{P0}(z = H) = C_\infty \tag{S75}$$

$$\left.\frac{\partial C_{P0}}{\partial r}\right|_{r=0} = 0 \tag{S76}$$

$$\left.\frac{\partial C_{P0}}{\partial z}\right|_{z=0} = 0 \tag{S77}$$

and initial conditions are as follows:

$$C_{P0}(r, z, t = 0) = 0 \quad \text{(S78)}$$

$$C_A(r, z, t = 0) = known\ constant \quad \text{(S79)}$$

$$C_X(r, z, t = 0) = 0 \quad \text{(S80)}$$

$$C_{PF}(r, z, t = 0) = 0 \quad \text{(S81)}$$

Solving and modeling this problem required a numerical solution, as the concentration equations for A, X, $P_0$, and $P_F$ are interdependent. Definitions for spatial discretization are as follows:

$$\Delta r = \frac{R}{M-1} \quad i = 1, 2, \ldots, M-1, M \quad \text{(S82)}$$

$$\Delta z = \frac{H}{N-1} \quad j = 1, 2, \ldots, N-1, N \quad \text{(S83)}$$

$$r_i = (i-1)\Delta r \quad \text{(S84)}$$

and definitions for time discretization are as follows:

$$\frac{\partial w}{\partial t} = \lim_{\Delta t \to 0} \frac{w(t) - w(t - \Delta t)}{\Delta t} \quad \text{(S85)}$$

$$w(t) \approx w(t - \Delta t) + \Delta t \left(\frac{\partial w}{\partial t}\right) \quad \text{(S86)}$$

We were able to discretize the non-diffusion species fairly easily:

$$C_{A(i,j,t)} \approx C_{A(i,j,t-1)} - fkC_{A(i,j,t-1)}C_{P0(i,j,t-1)}\Delta t \quad \text{(S87)}$$

$$C_{X(i,j,t)} \approx C_{X(i,j,t-1)} + fkC_{A(i,j,t-1)}C_{P0(i,j,t-1)}\Delta t \quad \text{(S88)}$$

$$C_{PF(i,j,t)} \approx C_{PF(i,j,t-1)} + kC_{A(i,j,t-1)}C_{P0(i,j,t-1)}\Delta t \quad \text{(S89)}$$

However, the diffusing free PEG becomes much more involved to discretize. Specifically, we needed to find separate solutions for the internal nodes, boundary condition at $r = R$, boundary condition at $z = H$, boundary condition at $z = 0$, axis of symmetry at $r = 0$, and the special case where both $r = 0$ and $z = 0$.

Beginning with the internal nodes, we discretized the second derivatives in **Equation S72** by adding a single forward and a single backward Taylor series expansion (achieves second order accuracy) for each derivative.

$$\frac{\partial^2 C_{P(i,j)}}{\partial r^2} = \frac{C_{P(i+1,j)} - 2C_{P(i,j)} + C_{P(i-1,j)}}{(\Delta r)^2} + \varepsilon[(\Delta r)^2] \quad \text{(S90)}$$

$$\frac{\partial^2 C_{P(i,j)}}{\partial z^2} = \frac{C_{P(i,j+1)} - 2C_{P(i,j)} + C_{P(i,j-1)}}{(\Delta z)^2} + \varepsilon[(\Delta z)^2] \quad \text{(S91)}$$

We discretized the first derivative in **Equation S72** by subtracting a single backward Taylor series expansion from a single forward Taylor series expansion for second order accuracy.

$$\frac{\partial C_{P(i,j)}}{\partial r} = \frac{C_{P(i+1,j)} - C_{P(i-1,j)}}{2(\Delta r)^2} + \varepsilon[(\Delta r)^2] \quad \text{(S92)}$$

Substituting **Equation S90-S92** into **Equation S72** and truncating the error term results in

$$\frac{\partial C_{P0(i,j)}}{\partial t} \approx -2D_{eff}\left[\frac{1}{(\Delta r)^2}+\frac{1}{(\Delta z)^2}\right]C_{P(i,j)} + D_{eff}\left[\frac{1}{(\Delta r)^2}+\frac{1}{2r_i\Delta r}\right]C_{P(i+1,j)} + D_{eff}\left[\frac{1}{(\Delta r)^2}-\frac{1}{2r_i\Delta r}\right]C_{P(i-1,j)} + \left[\frac{D_{eff}}{(\Delta z)^2}\right]C_{P(i,j+1)} + \left[\frac{D_{eff}}{(\Delta z)^2}\right]C_{P(i,j-1)} - kC_{A(i,j)}C_{P0(i,j)} \tag{S93}$$

and discretizing in time gives us **Equation S94** (valid at $i = 2, \dots, M-1$ and $j = 2, \dots, N-1$):

$$C_{P0(i,j,t)} \approx C_{P0(i,j,t-1)} - 2D_{eff}\left[\frac{1}{(\Delta r)^2}+\frac{1}{(\Delta z)^2}\right]C_{P(i,j,t-1)}\Delta t + D_{eff}\left[\frac{1}{(\Delta r)^2}+\frac{1}{2r_i\Delta r}\right]C_{P(i+1,j,t-1)}\Delta t + D_{eff}\left[\frac{1}{(\Delta r)^2}-\frac{1}{2r_i\Delta r}\right]C_{P(i-1,j,t-1)}\Delta t + \left[\frac{D_{eff}}{(\Delta z)^2}\right]C_{P(i,j+1,t-1)}\Delta t + \left[\frac{D_{eff}}{(\Delta z)^2}\right]C_{P(i,j-1,t-1)}\Delta t - kC_{A(i,j,t-1)}C_{P0(i,j,t-1)}\Delta t \tag{S94}$$

The external nodes at the $r = R$ and $z = H$ boundary conditions (hydrogel-solution interface) were derived according to the concentration value at those boundary conditions:

$$C_{P0}(r = R) \rightarrow C_{P0(M,j,t)} = C_\infty \tag{S95}$$

$$C_{P0}(z = H) \rightarrow C_{P0(i,N,t)} = C_\infty \tag{S96}$$

and are valid at $i = M; j = 1, \dots, N$ and $i = 1, \dots, M; j = N$, respectively. The remaining special cases fall at $r = 0$

$$C_{P0}(r = 0, 0 < z < H) \rightarrow C_{P0(1,j,t)} \quad j = 2, \dots, N-1\,, \tag{S97}$$

$z = 0$

$$C_{P0}(0 < r < R, z = 0) \rightarrow C_{P0(i,1,t)} \quad i = 2, \dots, M-1\,, \tag{S98}$$

and $(r, z) = (0,0)$

$$C_{P0}(r = 0, z = 0) \rightarrow C_{P0(1,1,t)}\,. \tag{S99}$$

At $r = 0$ (axis of symmetry), the governing equation results in a singularity:

$$\frac{1}{r}\frac{\partial C_P}{\partial r} = \frac{0}{0} \tag{S100}$$

Here, we can use L'Hopital's rule to eliminate the singularity:

$$\lim_{r\to 0}\frac{1}{r}\frac{\partial C_P}{\partial r} = \frac{\left.\frac{\partial}{\partial r}\left(\frac{\partial C_P}{\partial r}\right)\right|_{r=0}}{\left.\frac{\partial}{\partial r}(r)\right|_{r=0}} = \frac{\left.\frac{\partial^2 C_P}{\partial r^2}\right|_{r=0}}{1} = \left.\frac{\partial^2 C_P}{\partial r^2}\right|_{r=0} \tag{S101}$$

Therefore, at $r = 0$:

$$\frac{\partial C_{P0}}{\partial t} = D_{eff}\left[2\frac{\partial^2 C_P}{\partial r^2}+\frac{\partial^2 C_P}{\partial z^2}\right] - kC_AC_{P0} \tag{S102}$$

While **Equation S91** can be used for the second derivative in the *z*-direction, only forward Taylor expansions in the *r*-direction may be used as there is no $i = 0$. Here, we implemented 2 forward expansions to keep the discretized approximation second order, i.e., $i = 2, i = 3$ ($C_{P(2,j)}, C_{P(3,j)}$). By plugging in the axis of symmetry condition, we can multiply by a constant and subtract the Taylor series expansions to eliminate the third derivative term:

$$8C_{P(2,j)} - C_{P(3,j)} \tag{S103}$$

Solving for the second derivative term,

$$2\frac{\partial^2 C_{P(1,j)}}{\partial r^2} = \frac{-7C_{P(1,j)} + 8C_{P(2,j)} - C_{P(3,j)}}{(\Delta r)^2} + \varepsilon[(\Delta r)^2], \tag{S104}$$

we then substituted **Equation S91** and **S104** into the governing equation (**Equation S102**) and discretized over time:

$$\begin{aligned}\frac{\partial C_{P0(1,j)}}{\partial t} \approx & -D_{eff}\left[\frac{7}{(\Delta r)^2} + \frac{2}{(\Delta z)^2}\right]C_{P(1,j)} + \left[\frac{8D_{eff}}{(\Delta r)^2}\right]C_{P(2,j)} - \left[\frac{D_{eff}}{(\Delta r)^2}\right]C_{P(3,j)} \\ & + \left[\frac{D_{eff}}{(\Delta z)^2}\right]C_{P(1,j+1)} + \left[\frac{D_{eff}}{(\Delta z)^2}\right]C_{P(1,j-1)} - kC_{A(1,j)}C_{P0(1,j)}\end{aligned} \tag{S105}$$

$$\begin{aligned}C_{P0(1,j,t)} \approx & \; C_{P0(1,j,t-1)} - D_{eff}\left[\frac{7}{(\Delta r)^2} + \frac{2}{(\Delta z)^2}\right]C_{P(1,j,t-1)}\Delta t \\ & + \left[\frac{8D_{eff}}{(\Delta r)^2}\right]C_{P(2,j,t-1)}\Delta t - \left[\frac{D_{eff}}{(\Delta r)^2}\right]C_{P(3,j,t-1)}\Delta t \\ & + \left[\frac{D_{eff}}{(\Delta z)^2}\right]C_{P(1,j+1,t-1)}\Delta t + \left[\frac{D_{eff}}{(\Delta z)^2}\right]C_{P(1,j-1,t-1)}\Delta t \\ & - kC_{A(1,j,t-1)}C_{P0(1,j,t-1)}\Delta t\end{aligned} \tag{S106}$$

(valid at $i = 1$ and $j = 2, \ldots, N-1$).

Similar to the axis of symmetry, at $z = 0$ (bottom of hydrogel, touching the well plate) we required only forward Taylor expansions in the *z*-direction as there is no $j = 0$. Here, we implemented 2 forward expansions to keep the discretized approximation second order, i.e., $j = 2, j = 3$ ($C_{P(i,2)}, C_{P(i,3)}$). By plugging in the zero-flux boundary condition, we can multiply by a constant and subtract the Taylor series expansions to eliminate the third derivative term:

$$8C_{P(i,2)} - C_{P(i,3)} \tag{S107}$$

Solving for the second derivative term,

$$\frac{\partial^2 C_{P(i,1)}}{\partial z^2} = \frac{-7C_{P(i,1)} + 8C_{P(i,2)} - C_{P(i,3)}}{2(\Delta z)^2} + \varepsilon[(\Delta z)^2], \tag{S108}$$

we then substituted **Equation S90**, **S92** and **S108** into the governing equation (**Equation S72**) and discretized over time:

$$\begin{aligned}\frac{\partial C_{P0(i,1)}}{\partial t} \approx & -D_{eff}\left[\frac{2}{(\Delta r)^2} + \frac{7}{2(\Delta z)^2}\right]C_{P(i,1)} + \left[\frac{4D_{eff}}{(\Delta z)^2}\right]C_{P(i,2)} - \left[\frac{D_{eff}}{2(\Delta z)^2}\right]C_{P(i,3)} \\ & + D_{eff}\left[\frac{1}{(\Delta r)^2} + \frac{1}{2r_i\Delta r}\right]C_{P(i+1,1)} + D_{eff}\left[\frac{1}{(\Delta r)^2} - \frac{1}{2r_i\Delta r}\right]C_{P(i-1,1)} \\ & - kC_{A(i,1)}C_{P0(i,1)}\end{aligned} \tag{S109}$$

$$C_{P0(i,1,t)} \approx C_{P0(i,1,t-1)} - D_{eff}\left[\frac{2}{(\Delta r)^2} + \frac{7}{2(\Delta z)^2}\right] C_{P(i,1,t-1)}\Delta t + \left[\frac{4D_{eff}}{(\Delta z)^2}\right] C_{P(i,2,t-1)}\Delta t - \left[\frac{D_{eff}}{2(\Delta z)^2}\right] C_{P(i,3,t-1)}\Delta t + D_{eff}\left[\frac{1}{(\Delta r)^2} + \frac{1}{2r_i\Delta r}\right] C_{P(i+1,1,t-1)}\Delta t + D_{eff}\left[\frac{1}{(\Delta r)^2} - \frac{1}{2r_i\Delta r}\right] C_{P(i-1,1,t-1)}\Delta t - kC_{A(i,1,t-1)}C_{P0(i,1,t-1)}\Delta t \tag{S110}$$

(valid at $i = 2, \ldots, M-1$ and $j = 1$).

The final special case occurs at $(r, z) = (0,0)$. We again used **Equation S102** as the governing equation to avoid the singularity at $r = 0$, with forward Taylor series expansions in both the *r*- and *z*-directions (**Equation S104** and **S108**):

$$\frac{\partial C_{P0(1,1)}}{\partial t} \approx -D_{eff}\left[\frac{7}{(\Delta r)^2} + \frac{7}{2(\Delta z)^2}\right] C_{P(1,1)} + \left[\frac{8D_{eff}}{(\Delta r)^2}\right] C_{P(2,1)} - \left[\frac{D_{eff}}{(\Delta r)^2}\right] C_{P(3,1)} + \left[\frac{4D_{eff}}{(\Delta z)^2}\right] C_{P(1,2)} - \left[\frac{D_{eff}}{2(\Delta z)^2}\right] C_{P(1,3)} - kC_{A(1,1)}C_{P0(1,1)} \tag{S111}$$

$$C_{P0(1,1,t)} \approx C_{P0(1,1,t-1)} - D_{eff}\left[\frac{7}{(\Delta r)^2} + \frac{7}{2(\Delta z)^2}\right] C_{P(1,1,t-1)}\Delta t + \left[\frac{8D_{eff}}{(\Delta r)^2}\right] C_{P(2,1,t-1)}\Delta t - \left[\frac{D_{eff}}{(\Delta r)^2}\right] C_{P(3,1,t-1)}\Delta t + \left[\frac{4D_{eff}}{(\Delta z)^2}\right] C_{P(1,2,t-1)}\Delta t - \left[\frac{D_{eff}}{2(\Delta z)^2}\right] C_{P(1,3,t-1)}\Delta t - kC_{A(1,1,t-1)}C_{P0(1,1,t-1)}\Delta t \tag{S112}$$

(valid at $i = 1$ and $j = 1$).

*MATLAB model of PEG diffusion and reaction within a hydrogel with a changing outer boundary condition*

Similar to the conditioned media case, we chose to treat $C_\infty$ as a step function, in which $C_{\infty,a}$ was updated at each hour, *a* of the model. We calculated the updated $C_{\infty,a+1}$ with the following equation (where timestep *t* and timestep *a* represent the same real time unit):

$$C_{\infty,a+1} = \frac{C_0 V_S - n_t}{V_S} \tag{S113}$$

where $C_0$ is the initial concentration of PEG in solution and $V_S$ is the volume of the bulk solution. $n_t$ is the total number of moles of PEG (free and fixed) within the hydrogel:

$$n_t = \pi R \sum_{i=1}^{M-1}\sum_{j=1}^{N-1} \Delta r\,\Delta z \frac{C_{P(i,j,t)} + C_{P(i+1,j,t)} + C_{P(i,j+1,t)} + C_{P(i+1,j+1,t)}}{4}, \tag{S114}$$

determined by adding the number of moles of PEG within each square between nodes on the *r-z* axis and integrating over the remaining volume of the hydrogel.

The MATLAB model was run for the equivalent of the 72-hour cell experiment, during which, the hydrogel dimensions were assumed to remain constant (although we did see some increases in hydrogel volume over the course of stiffening: **Figure S28**). Each incubation was modeled over the course of 12 hours, following the cycle parameters used in the stiffening experiments. At the end of each model incubation period, the concentration of "inert" PEG functional groups was calculated ($C_{B0(i,j,t)} = C_{PF(i,j,t)}[4 - f]$) and any unreacted hydrogel functional group (A, $C_{A(i,j,t)}$) was stored as a fixed Azide or BCN concentration, as appropriate. The new hydrogel functional group "initial condition" for the following incubation was then calculated by adding $C_{B0(i,j,t)}$ to the previously stored fixed Azide or BCN concentration (according to the identity of B). The remaining concentration values ($C_{X(i,j,t)}$, $C_{P0(i,j,t)}$, and $C_{PF(i,j,t)}$) were reset to zero for the beginning of the next incubation. Importantly, this approach neglects any remaining free PEG within the hydrogel at the time of the incubation switch. While this allows us to simplify the model, it results in the negation of (1) any reactions that will continue to take place between the hydrogel and remaining free PEG during the next incubation and (2) any reactions between the remaining free PEG-Azide/BCN and the PEG-xBCN/Azide diffusing into the hydrogel during the next incubation. Consequently, this model underestimates the number of linkages that any remaining free PEG will form.

Finally, the increase in crosslink density was approximated at the end of each cycle by calculating the concentration range of fixed PEG ($C_{PF}$) reacted throughout the hydrogel during the cycle. This approach assumed that each 4-arm PEG molecule reacted to the hydrogel will constitute one crosslink by the end of the cycle (for example, incubation 1A is expected to result in dangling PEG molecules, and will therefore not contribute crosslinks until incubation 1B has taken place, such that by the end of Cycle 1, PEG molecules reacted with the hydrogel from both incubations will theoretically contribute new crosslinks). This assumption will result in an overestimate of crosslink density, as it does not account for network loops or defects that will not contribute to crosslink density. The cumulative crosslink density increase from stiffening was calculated as: Cycle 1 = 1.28-1.93 mM, Cycle 2 = 2.86-4.24 mM, and Cycle 3 = 4.04-5.97 mM, where the ranges indicate the lowest crosslink density (bottom middle) to the highest crosslink density (outer boundary) throughout the hydrogel at the end of each cycle.

*Hydrogel property calculations*

Mesh size and crosslink density were calculated for use in the modeling portion of this work. The mesh size for the initial hydrogel was calculated using the rubber elasticity theory:[27]

$$\xi = v_s^{-1/3} \ell \left( \frac{2C_n \overline{M_c}}{M_r} \right)^{1/2} , \tag{S115}$$

where $\xi$ is the mesh size, $v_s$ is the polymer volume fraction in the swollen state, $\ell$ is the bond length along the PEG backbone (~ 0.15 nm), $C_n$ is the Flory characteristic ratio of PEG ($C_n = 4$), $\overline{M_c}$ is the molecular weight between crosslinks, and $M_r$ is the PEG repeat unit molecular weight (44 g/mol). $\overline{M_c}$ was calculated from the equation:

$$\frac{1}{\overline{M_c}} = \left( \frac{v_r}{v_s} \right)^{1/3} \frac{G'}{\rho RT} + \frac{2}{\overline{M_n}}, \tag{S116}$$

where $v_r$ is the polymer volume fraction in the relaxed state (i.e., at preparation), $G'$ is the equilibrium swollen storage modulus, $\rho$ is the density of PEG (1.07 g/mL), $R$ is the universal gas constant (8.314 $m^3$ Pa $K^{-1}$ $mol^{-1}$ = 8134 L Pa $K^{-1}$ $mol^{-1}$), $T$ is the temperature (37 °C = 310 K), and $\overline{M_n}$ is the total monomer average molecular weight (calculated to be 9709 g/mol for this hydrogel composition). $v_s$ and $v_s$ were calculated from equations:

$$\nu_r = \frac{1}{Q_r}, \nu_s = \frac{1}{Q_s}, \tag{S117}$$

where $Q_r$ and $Q_s$ are the swelling ratios in the relaxed (at preparation) and equilibrium swollen states, respectively.

At preparation and dry volumes were not measured for each hydrogel; rather, these measurements were assumed to be 20 μL (preparation, according to precursor solution volume) and 2.52 μL (dry, calculated from the hydrogel component molecular weights and assuming the density of PEG, as PEG is the main component with the dominating molecular weight). Mesh size was calculated to be ξ = 16.7 nm. This calculation was made neglecting any contributions from the linker peptide or mfCMP to the $C_n$ and $M_r$ values. Crosslink density was calculated using the equation:

$$\rho_x = \frac{G' Q_s^{1/3}}{RT}, \tag{S118}$$

finding that $\rho_x = 0.880$ mM in the initial hydrogel.

*Effective diffusivity calculations*

The effective diffusivity for PEG and PDGF were calculated for modeling diffusion through the initial hydrogel. We first calculated the diffusivity in water for each solute using the Stokes-Einstein equation:[28]

$$D_\infty = \frac{k_B T}{6\pi\eta r_h}, \tag{S119}$$

where $k_B$ is the Boltzmann constant (1.38 × $10^{-23}$ J/K), $\eta$ is the dynamic viscosity of water at 37 °C (0.000691 kg $m^{-1}$ $s^{-1}$), and $r_h$ is the hydrodynamic radius of the solute (3.5 nm for PDGF[29] and 6.83 nm for 4-arm 20 kDa PEG[30]). We found that for PDGF, $D_\infty = 9.38 \times 10^{-7}$ cm$^2$ s$^{-1}$, and for PEG, $D_\infty = 4.81 \times 10^{-7}$ cm$^2$ s$^{-1}$. We then approximated the effective diffusivity using an obstruction theory equation:[31]

$$D_{eff} = D_\infty \exp\left[-\pi \left(\frac{r_h + r_f}{\xi + 2r_f}\right)^2\right], \tag{S120}$$

where $r_f$ is the polymer chain radius (0.51 nm for PEG[32]). This resulted in $D_{eff} = 7.99 \times 10^{-7}$ cm$^2$ s$^{-1}$ for PDGF and $D_{eff} = 2.80 \times 10^{-7}$ cm$^2$ s$^{-1}$ for PEG within this hydrogel composition.

#### 4. Supplemental Figures

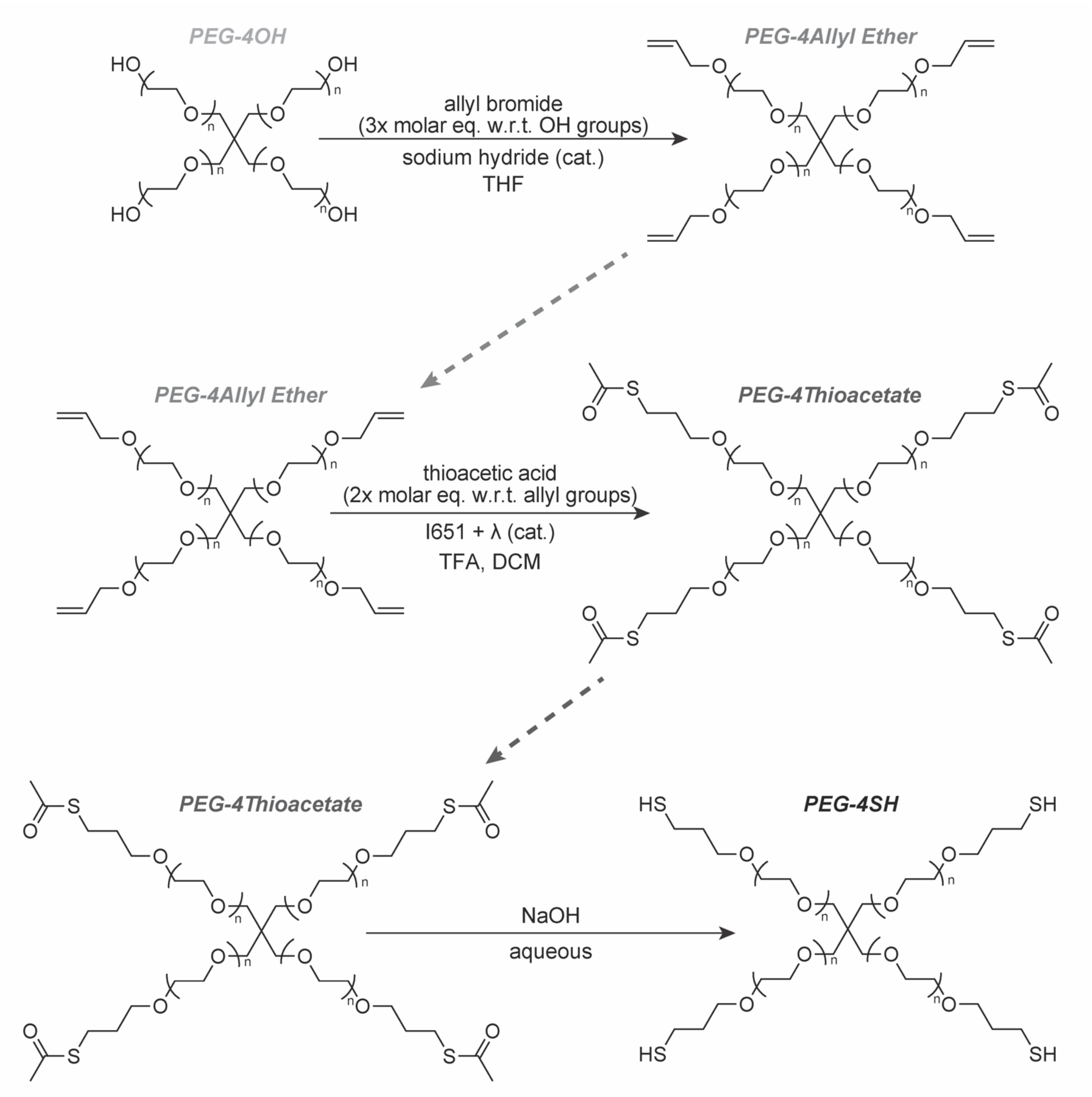

**Figure S1. Poly(ethylene glycol) tetra-thiol (PEG-SH) polymer synthesis.** Reaction schematic for functionalization of PEG-SH.

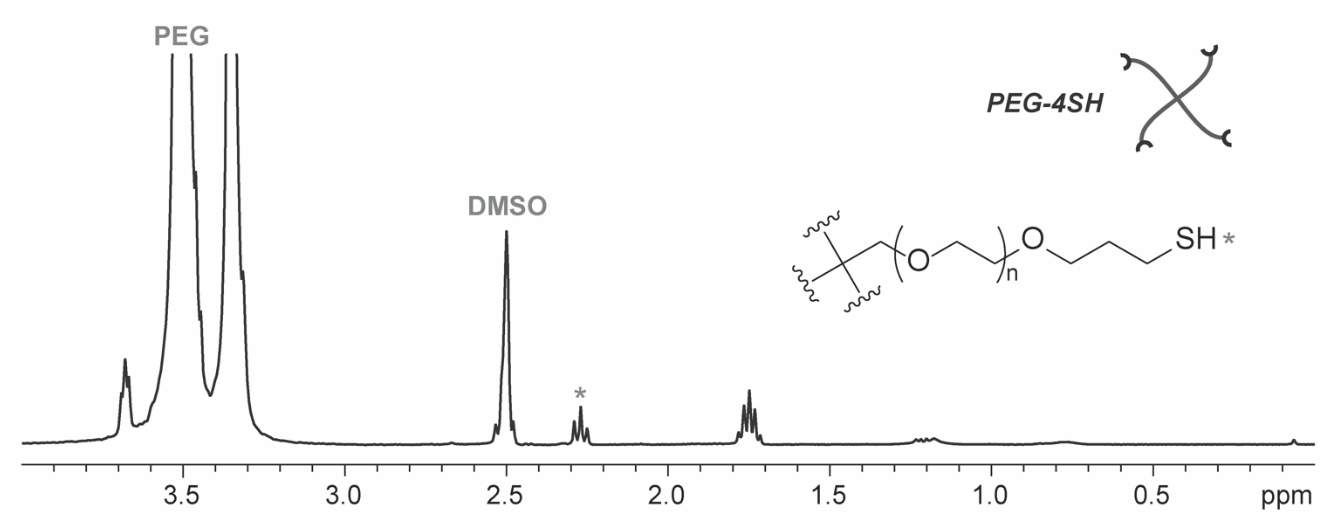

**Figure S2. PEG-SH $^1$H NMR.** $^1$H NMR spectrum (400 MHz, DMSO-$d_6$) of thiol-functionalized 4-arm PEG ($M_n$ ~ 20 kDa): δ 3.51 (s, 454H: PEG backbone), 2.27 (t, $J$ = 7.9 Hz, 1H: *), 1.75 (p, $J$ = 6.6 Hz, 2H). Functionality = 100% based on the proton shift at 2.27.

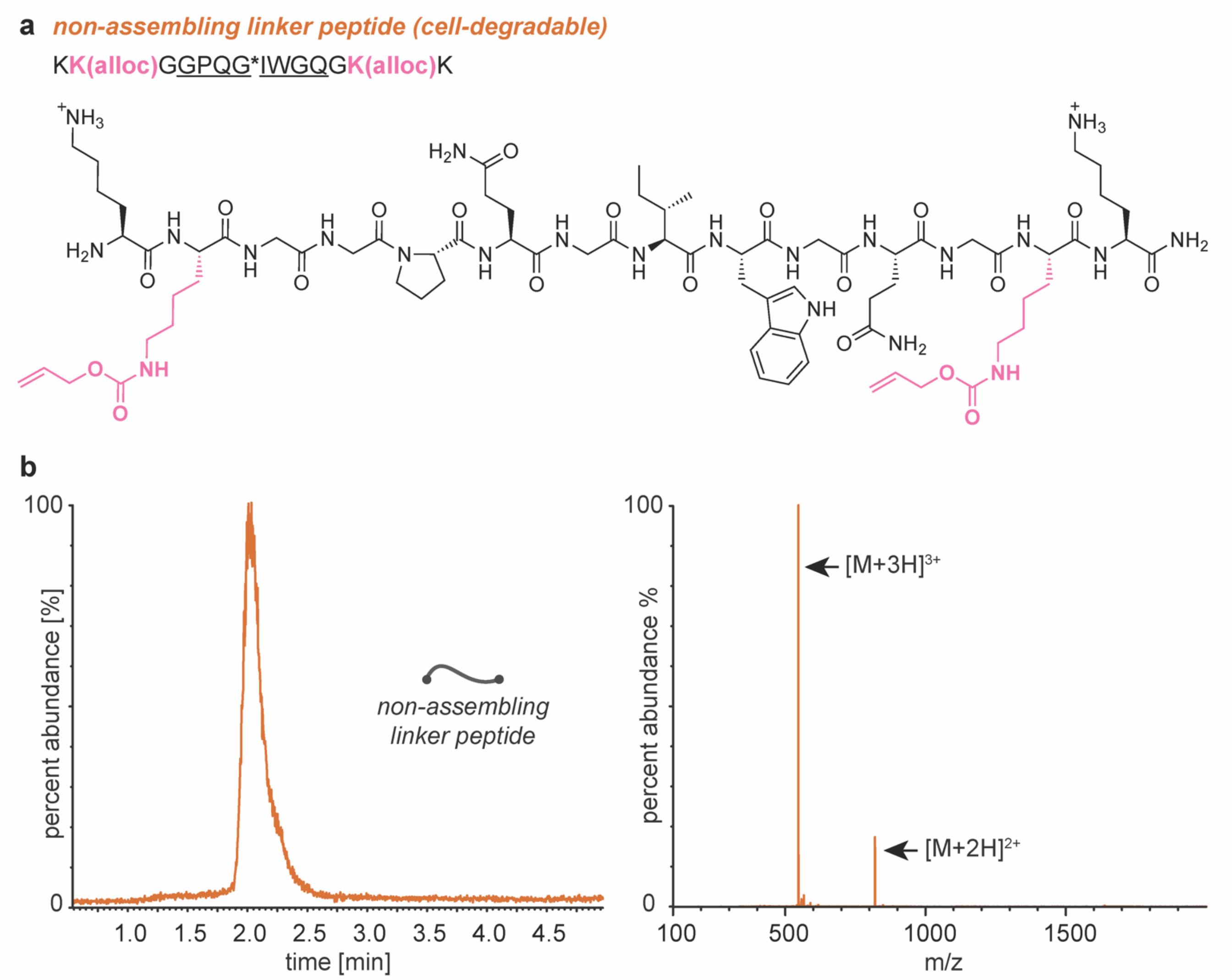

**Figure S3. Non-assembling linker peptide characterization.** a) Sequence and structure of the non-assembling, cell-degradable linker peptide: KK(alloc)GGPQG*IWGQGK(alloc)K (the underlined residues represent the matrix metalloproteinase (MMP)-responsive portion of the sequence, where * represents the cleavage site). The K(alloc) residues are displayed in pink for easy identification of the functional group. b) Single Quadruple Detector 2 (SQD2) UPLC trace (left) and electrospray ionization (ESI+) mass spectrometry (right) of the linker peptide. The desired product was observed with the expected molecular weight of 1636 g/mol ($[M+2H]^{2+}$ = 819 g/mol; $[M+3H]^{3+}$ = 546 g/mol).

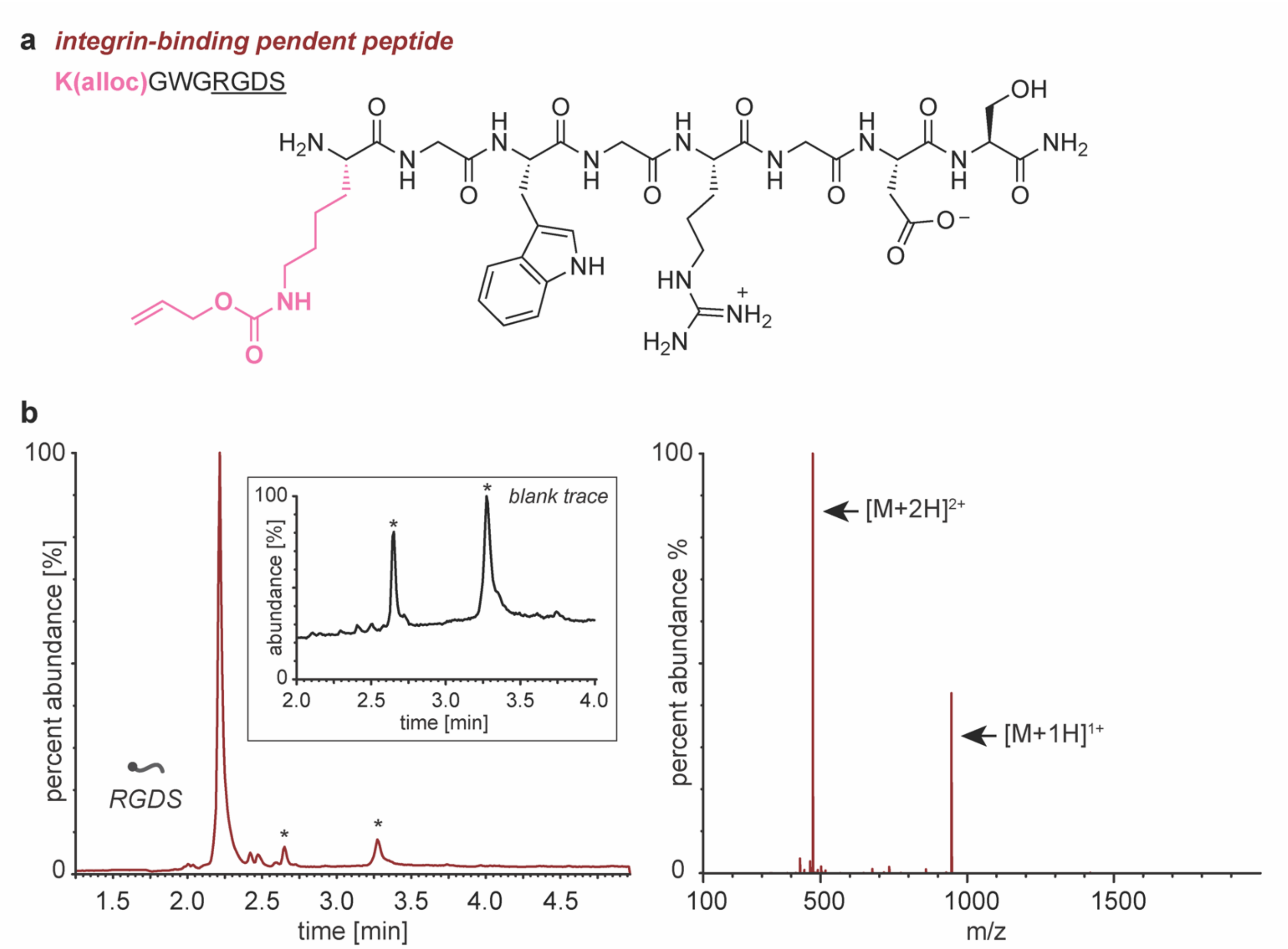

**Figure S4. Integrin-binding pendent peptide characterization.** a) Sequence and structure of the integrin-binding pendent peptide: K(alloc)GWG<u>RGDS</u> (the underlined residues represent the cell-recognized integrin-binding portion of the sequence). The K(alloc) residue is displayed in pink for easy identification of the functional group. b) Xevo G2-S QTof (XEVO) UPLC trace (left) and electrospray ionization (ESI+) mass spectrometry (right) of the pendent peptide. The desired product was observed with the expected molecular weight of 945 g/mol ($[M+1H]^{1+}$ = 946 g/mol; $[M+2H]^{2+}$ = 474 g/mol). A blank UPLC trace is shown in the inset image to demonstrate the that the starred peaks (not included in the ESI+ MS) were not in the peptide sample, but something on the column itself.

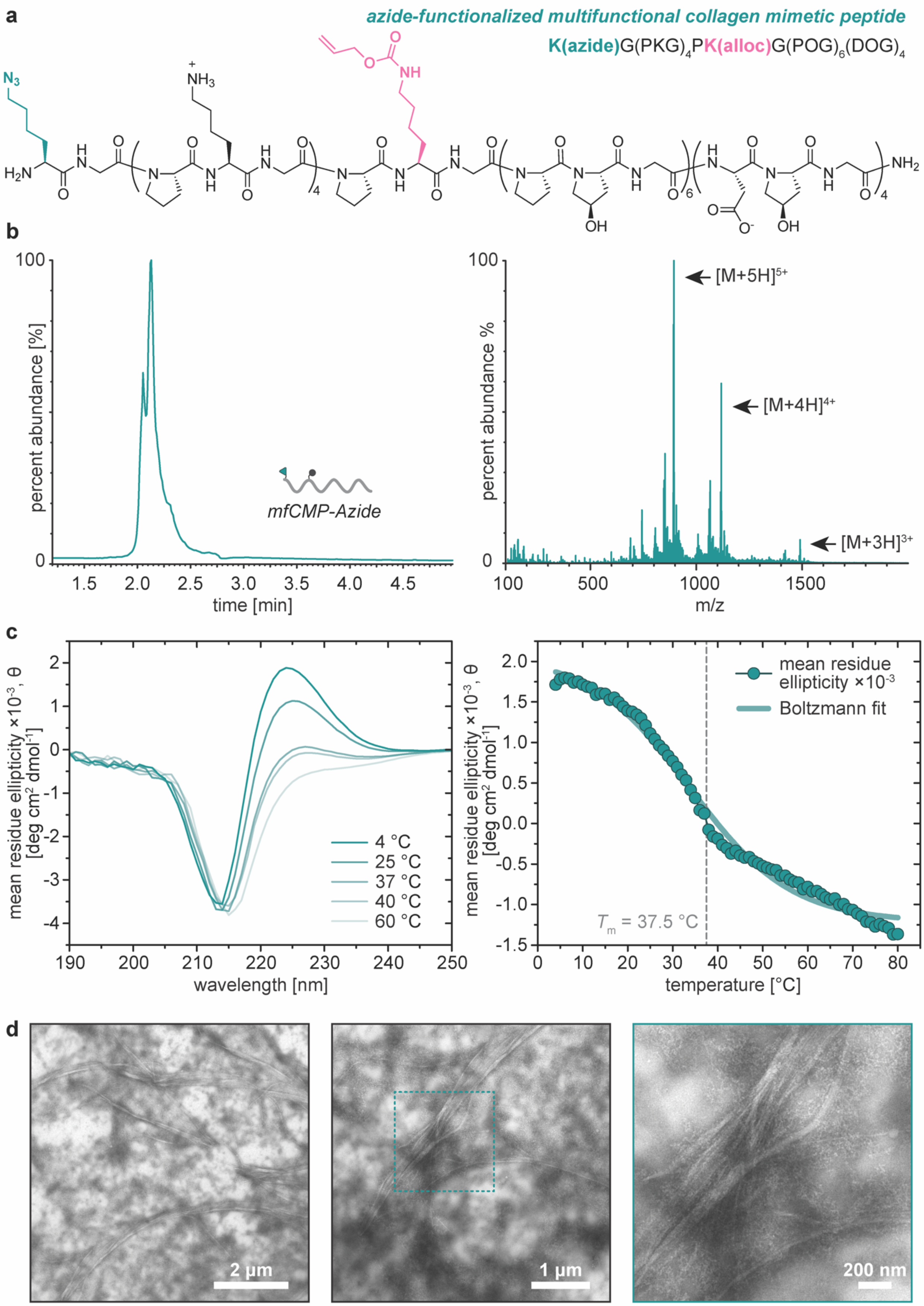
a
azide-functionalized multifunctional collagen mimetic peptide
K(azide)G(PKG)4PK(alloc)G(POG)6(DOG)4
b
percent abundance [%]
mfCMP-Azide
time [min]
percent abundance %
[M+5H]5+
[M+4H]4+
[M+3H]3+
m/z
c
mean residue ellipticity ×10-3, θ [deg cm2 dmol-1]
4 °C
25 °C
37 °C
40 °C
60 °C
wavelength [nm]
mean residue ellipticity ×10-3
Boltzmann fit
Tm = 37.5 °C
temperature [°C]
d
2 µm
1 µm
200 nm

**Figure S5. Azide-functionalized multifunctional collagen-mimetic peptide (mfCMP-Azide) characterization.** a) Sequence and structure of mfCMP-Azide: $K(azide)G(PKG)_4PK(alloc)G(POG)_6(DOG)_4$. The K(alloc) residue is displayed in pink and the K(azide) in teal for easy identification of the functional groups. b) Xevo G2-S QTof (XEVO) UPLC trace (left) and electrospray ionization (ESI+) mass spectrometry (right) of mfCMP-Azide. The desired product was observed with the expected molecular weight of 4469 g/mol ($[M+3H]^{3+}$ = 1491 g/mol, $[M+4H]^{4+}$ = 1118 g/mol, and $[M+5H]^{5+}$ = 895 g/mol). c) Circular dichroism (CD) of mfCMP-Azide (0.3 mM in 1× DPBS). Wavelength scans at increasing temperatures (left) indicate disassociation of the triple helices with rising temperature (decreasing peak at 225 nm). Temperature scans (right, 225 nm) were used to determine the melting temperature ($T_m$, where 50% of the peptide is assembled into triple helices and the remainder are individual peptide strands). The temperature scan was fit to a Boltzmann curve, which we would typically use to calculate the $T_m$ by finding the inflection point of the curve. However, we have observed that while a Boltzmann fit works well for peptides in DI water, the fit deviates from the data for peptides in DPBS (a common issue for POG-based sequences,[33] particularly near the $T_m$), leading to less accurate $T_m$ analysis (see Ford and Kloxin, 2021: Figures S7b and S8 as an example).[7] To avoid error stemming from using an ill-fitting curve, the slope between each point of the temperature scan was calculated to find the derivative of the data curve. The derivative minimum was used to determine the $T_m$, which was calculated by finding the temperature at the center of the derivative minimum: $T_m = 37.5 \pm 0.008$ °C. d) Transmission electronmicroscopy was used to confirm the presence assembled fibrils; representative images at 3 different magnifications are shown (assembled in 1× DPBS at 1 mM and diluted in DI water to 0.3 mM immediately before drop casting).

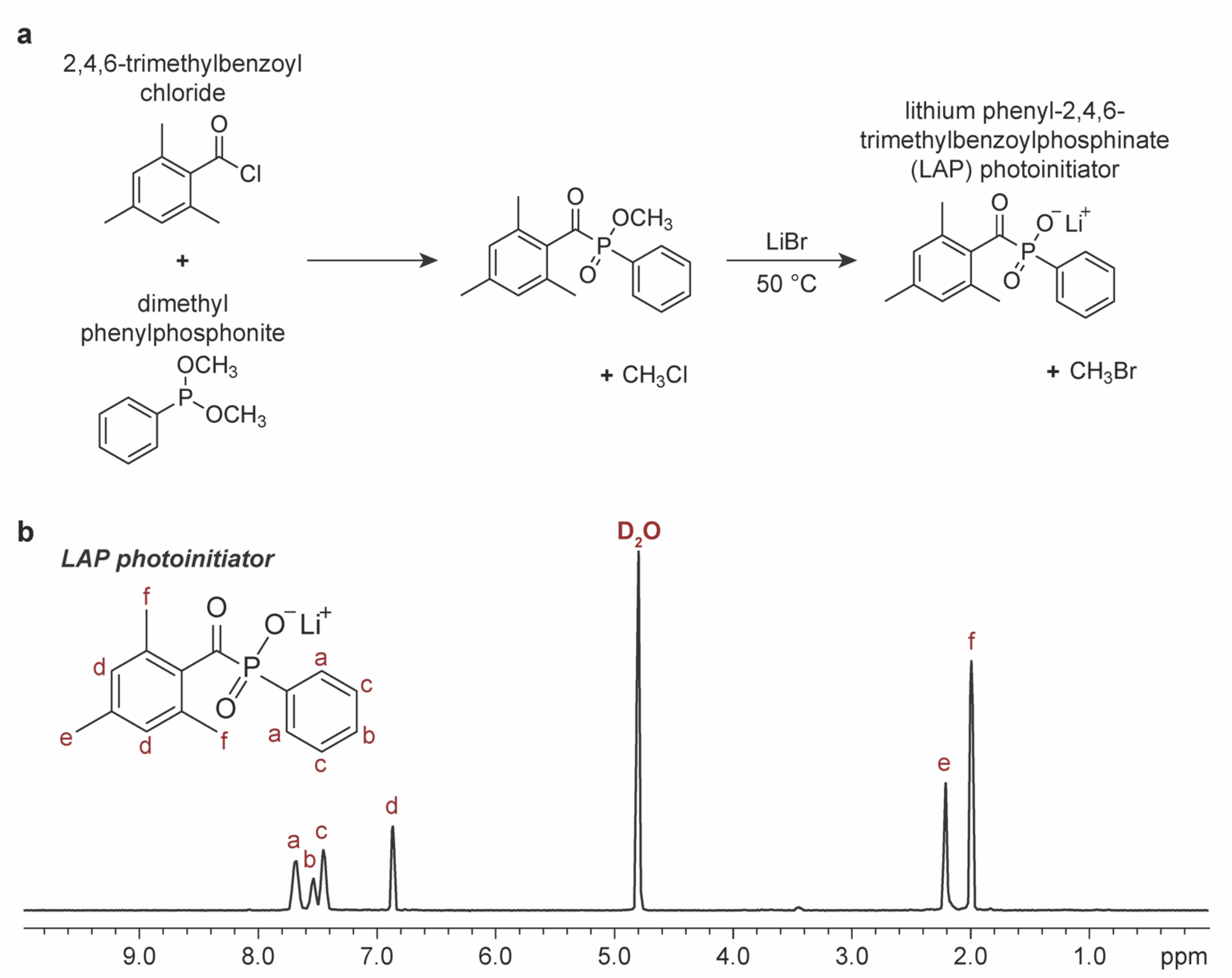

**Figure S6. Lithium phenyl-2,4,6-trimethylbenzoylphosphinate (LAP) synthesis.** a) Reaction schematic for LAP synthesis. b) $^{1}$H NMR spectrum (400 MHz, Deuterium Oxide) of LAP photoinitiator: δ 7.68 (dd, $J$ = 11.4, 6.6 Hz, 2H: a), 7.55 (d, $J$ = 8.1 Hz, 1H: b), 7.46 (t, $J$ = 7.8 Hz, 2H: c), 6.87 (t, $J$ = 5.0 Hz, 2H: d), 2.21 (d, $J$ = 5.3 Hz, 3H: e), 2.00 (t, $J$ = 4.8 Hz, 6H: f). Protons and associated peaks are indicated.

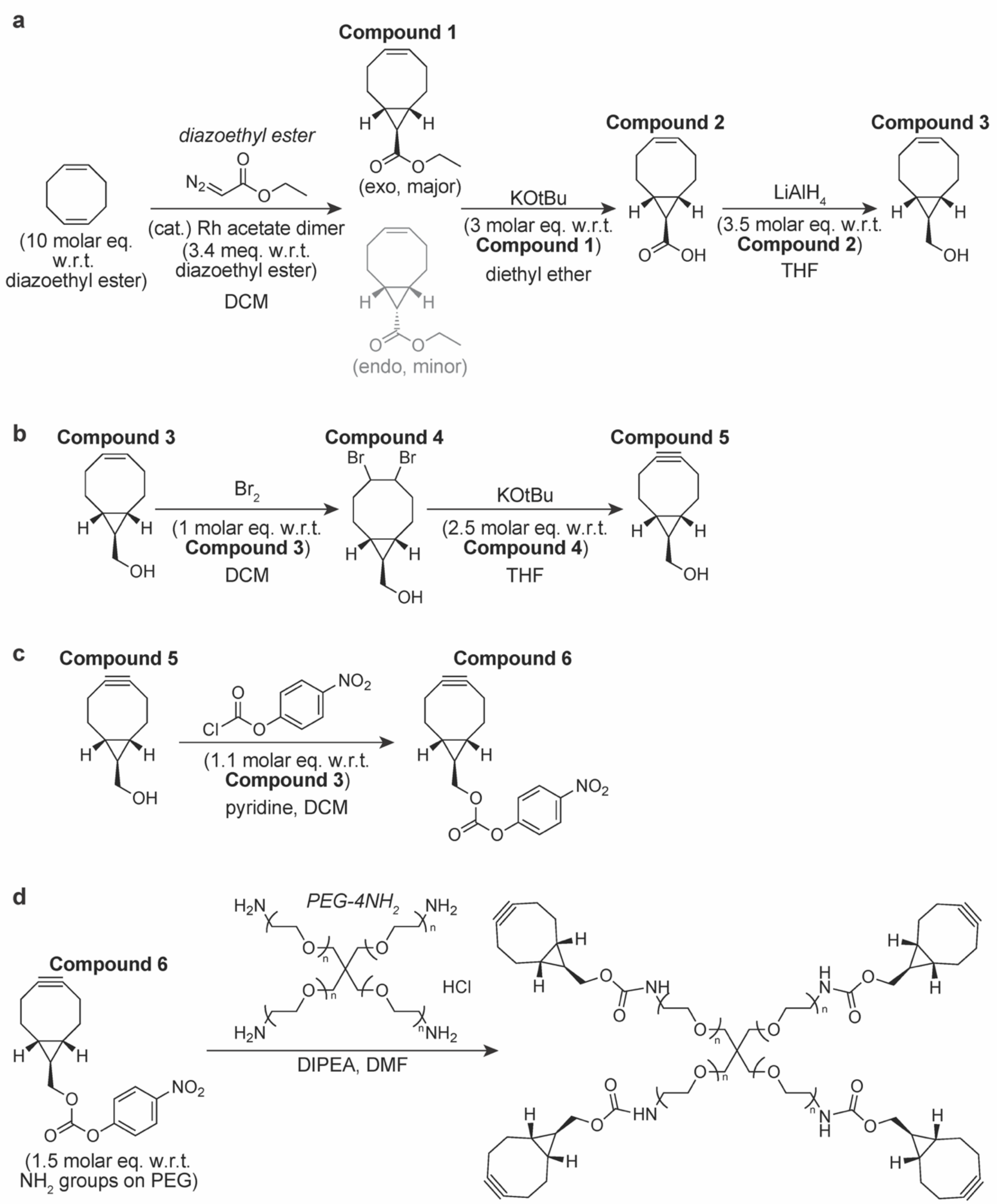

**Figure S7. Poly(ethylene glycol) tetra-bicyclononyne (*exo*) (PEG-xBCN) synthesis.** Reaction schematics for: a) (1*R*,8*S*,9*R*,4*Z*)-bicyclo[6.1.0]non-4-ene-9-yl-methanol synthesis, b) (1*R*,8*S*,9*R*)-bicyclo[6.1.0]non-4-yn-9-yl-methanol synthesis, c) (1*R*,8*S*,9*R*)-bicyclo[6.1.0]non-4-yn-9-yl-methyl (4-nitrophenyl) carbonate synthesis, and d) four-arm poly(ethylene glycol) tetra-bicyclononyne (*exo*) (PEG-xBCN) functionalization.

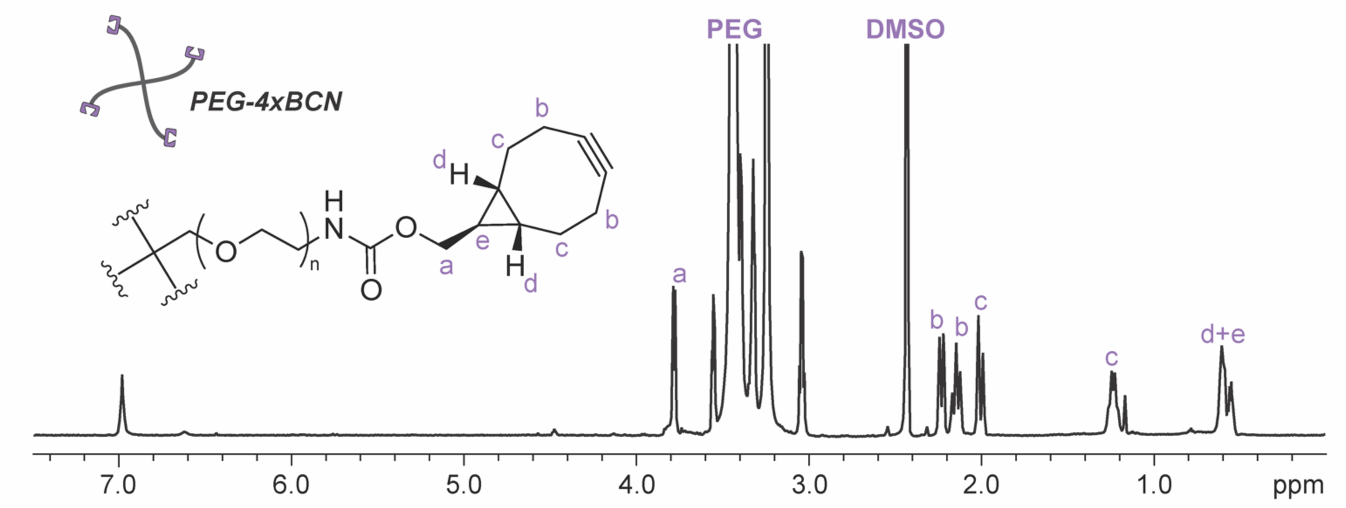

**Figure S8. PEG-xBCN $^1$H NMR.** $^1$H NMR spectrum (600 MHz, DMSO-$d_6$) of BCN-functionalized 4-arm PEG ($M_n$ ~ 10 kDa): δ 3.85 (d, $J$ = 6.8 Hz, 2H: a), 3.51 (s, 227H: PEG backbone), 2.35 – 2.26 (m, 2H: b), 2.22 (t, $J$ = 14.1 Hz, 2H: b), 2.11 – 2.03 (m, 2H: c), 1.36 – 1.25 (m, 2H: c), 0.74 – 0.58 (m, 3H: d+e). Functionality = 85% based on the proton shifts at 3.85, 2.35 – 2.26, 2.22, 2.11 – 2.03, 1.36 – 1.25, and 0.74 – 0.58.

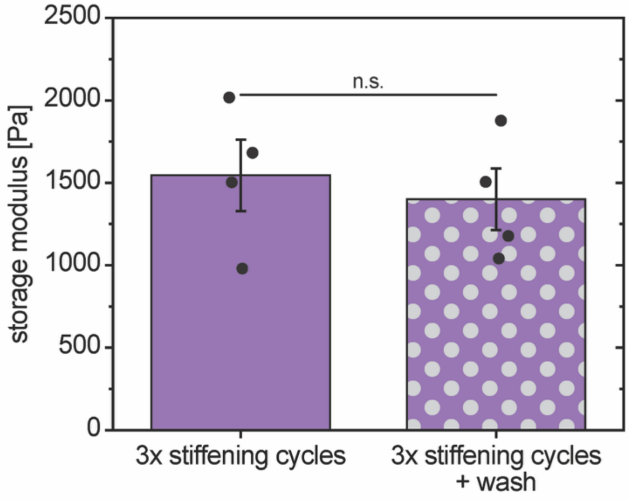

**Figure S9. Impact of washes on modulus after stiffening.** After 3 stiffening cycles, thorough washes (3× 1-hour washes in 1% FBS media at 37 °C and 5% $CO_2$ with humidity) were conducted to remove any remaining monomers. Oscillatory rheometry was performed on stiffened hydrogels before and after washes, demonstrating that washes after stiffening did not result in any changes to the modulus. All incubations and measurements were conducted at 37 °C. (Mean ± SE with individual datapoints; n = 4 independent samples; Significance determined via one-way ANOVA followed by Tukey's post-hoc test: * $p < 0.05$, n.s. indicates that the means are not significantly different.)

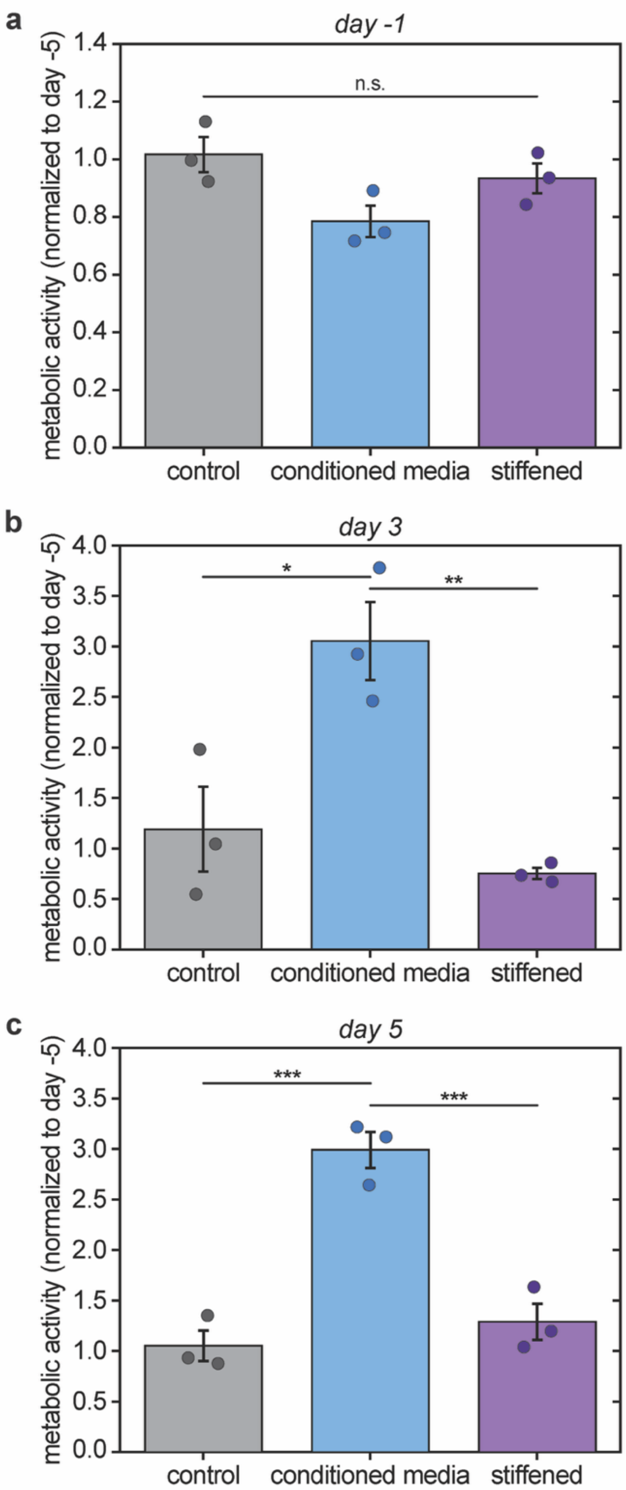

**Figure S10. Tukey's post-hoc multiple comparisons of metabolic activity at each timepoint.** Metabolic activity was assessed throughout the experiment both before (days -9 to 0) and after (days 0 to 5) implementation of experimental stimuli, where each hydrogel was internally normalized to its day -5 metabolic activity. a) Day -1. b) Day 3. c) Day 5. (Mean ± SE with individual datapoints; n = 3 independent samples; Significance determined via one-way ANOVA followed by Tukey's post-hoc test: * indicates a significant difference, where * $p < 0.05$, ** $p < 0.01$, *** $p < 0.001$; n.s. indicates that the means are not significantly different.) Statistical analyses within each condition over time are available in **Table S3**.

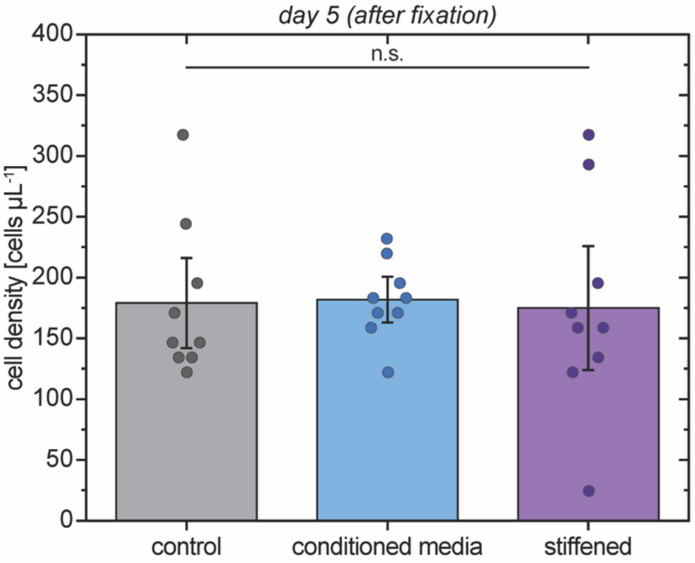

**Figure S11. Cell density at day five.** Hoechst-stained nuclei were detected and counted after fixation (14 days after encapsulation, 5 days after stimuli introduced). Some nuclei were not located within a cell (determined by F-Actin labeling). The cells belonging with these nuclei were classified as ‘not intact at the time of fixation’ and were excluded from the nuclei count. There were no differences between conditions in intact cell count after 5 days of treatment. (Mean ± SE with individual datapoints each representing the measurement from a single image; n = 3 independent samples, with measurements from ≥40 cell objects—totaling ≥125 individual cells—per condition; Significance determined via one-way ANOVA followed by Tukey’s post-hoc test: n.s. indicates that the means are not significantly different.)

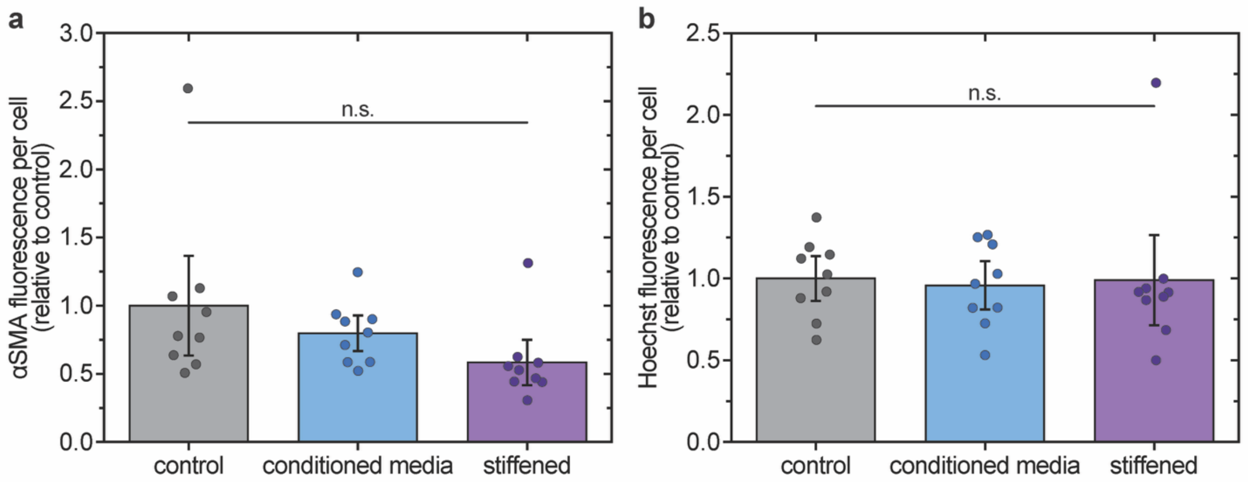

**Figure S12. Fixed cell fluorescence normalization to total nuclei count.** Cells were fixed and stained for F-Actin, nuclei, and αSMA (main text **Figure 3b**). Endpoint quantification of αSMA protein expression was determined by normalizing the total αSMA protein in each image to the total nuclear content (main text **Figure 3c**): the total fluorescence of the far-red channel (αSMA) was normalized to the total fluorescence of the blue channel (Hoechst). To confirm that normalization to the blue florescence (rather than nuclei count) was not artificially skewing the data, we a) normalized the αSMA fluorescence to the total nuclei count, and b) normalized the total Hoechst fluorescence to the total nuclei count. All conditions were normalized to the control. Normalization was performed relative to the total number of nuclei (including naked nuclei) or total Hoechst fluorescence to avoid the αSMA associated with cell debris around the naked nuclei from artificially skewing the results. (Mean ± SE with individual datapoints each representing the measurement from a single image; n = 3 independent samples, with measurements from ≥40 cell objects—totaling ≥125 individual cells—per condition; Significance determined via one-way ANOVA followed by Tukey's post-hoc test: n.s. indicates that the means are not significantly different.)

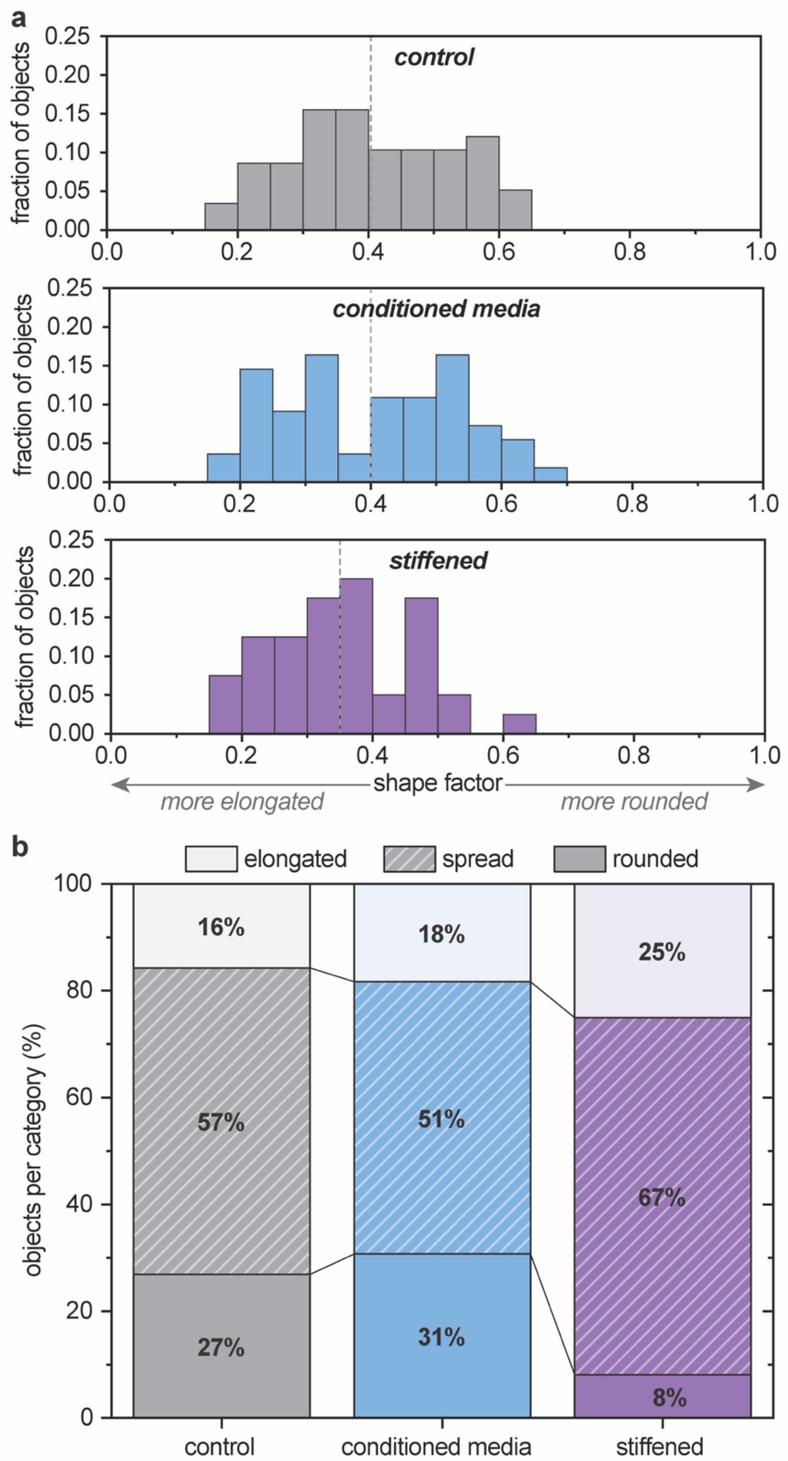

**Figure S13. Fibroblast morphology analysis.** After 5 days of treatment with conditioned media or a stiffening environment, fibroblasts were fixed, and cell morphology was analyzed using cytoskeletal staining. 3D shape factor ($\Psi$, a measurement of sphericity) of cell objects (individual cells or cell clusters) was calculated from the equation: $\Psi = \frac{\pi^{1/3}(6V)^{2/3}}{A}$. ($\Psi$ is the shape factor, $V$ is the volume of the object, and $A$ is the surface area of the object). a) Histograms for each condition demonstrate the shape factor distribution, where the dashed line indicates the shape factor mean for each condition. When comparing all conditions together with one-way ANOVA, there were no significant differences in overall shape factor. However, when comparing only the control and stiffened conditions, the overall shape factor of cells after stiffening is significantly lower (indicating greater elongation) than that of cells in the control ($p = 0.021$; direct comparison with Student's t-test). b) The shape factor data were then split into three categories according to elongation: *elongated* ($0 \leq \Psi \leq 0.25$), *spread* ($0.25 < \Psi \leq 0.5$), and *rounded* ($0.5 < \Psi \leq 1$). While there is a trend toward fewer rounded cells and more elongated cells in the stiffened condition, there were no statistical differences in the percent of objects per category between the three conditions. ($n = 3$ independent samples, with measurements from ≥40 cell objects—totaling ≥125 individual cells—per condition; Significance determined via one-way ANOVA followed by Tukey's post-hoc test.)

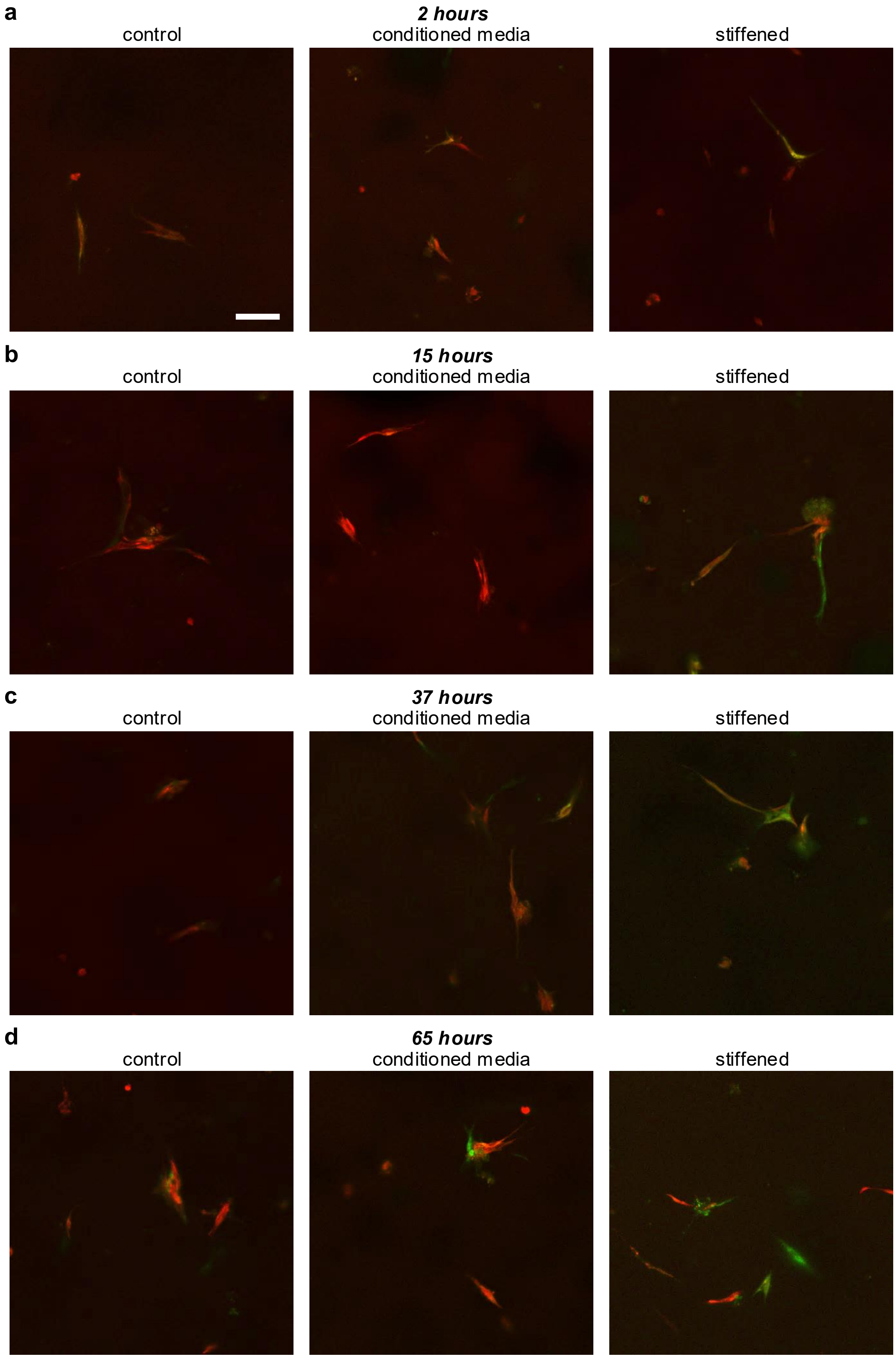
a
2 hours
control
conditioned media
stiffened
b
15 hours
control
conditioned media
stiffened
c
37 hours
control
conditioned media
stiffened
d
65 hours
control
conditioned media
stiffened

**Figure S14. Representative live reporter images of fibroblast αSMA expression.** Live reporter results at a) 2 hours, b) 15 hours, c) 37 hours, and d) 65 hours after stimuli were introduced. For each timepoint representative live reporter images are shown: orthogonal projections of confocal *z*-stacks; scale bar = 100 μm; DsRed constitutive reporter (red), ZsGreen αSMA reporter (green), and overlap (orange-yellow).

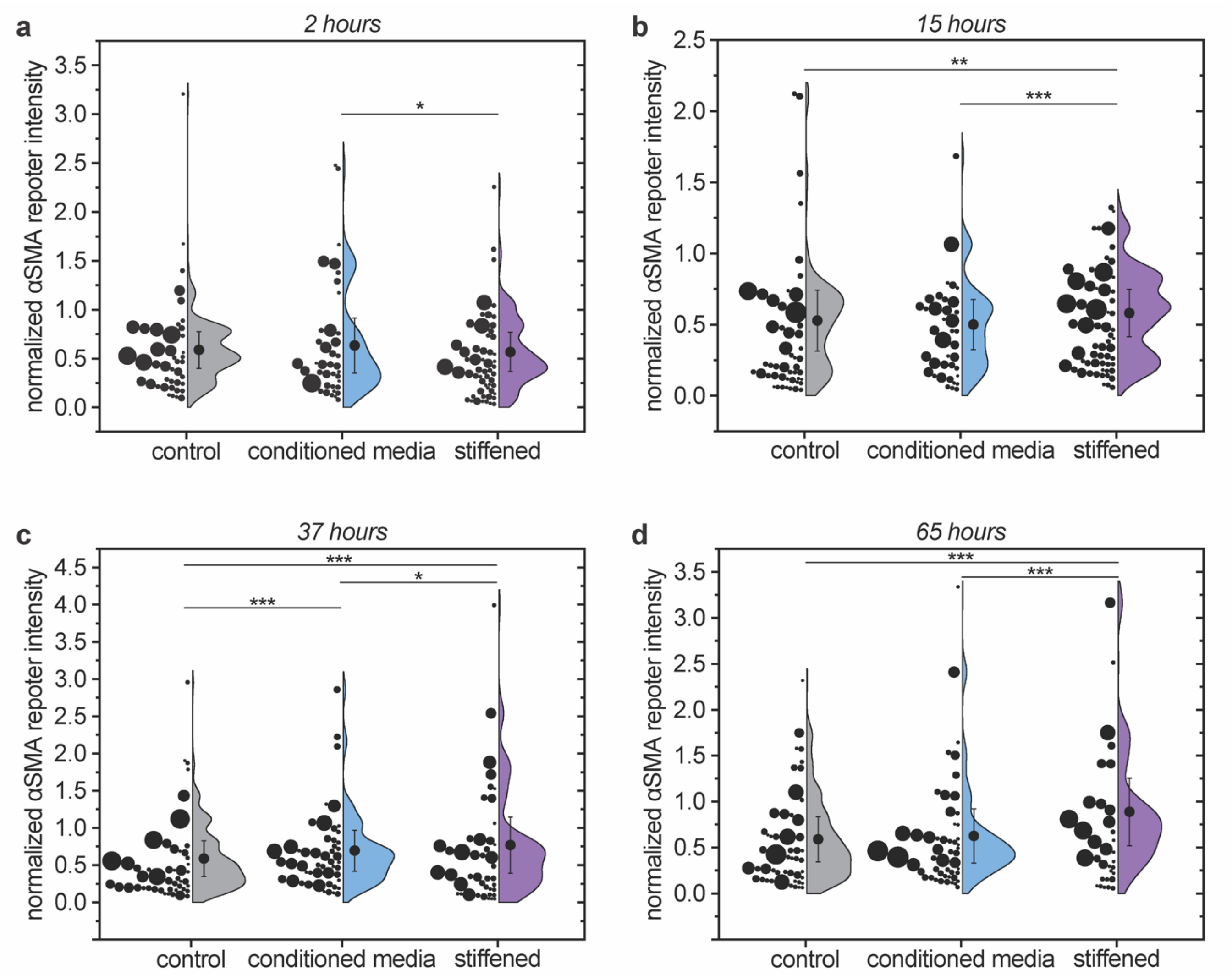

**Figure S15. Fibroblast αSMA expression and Tukey's post-hoc multiple comparisons at each timepoint.** Live reporter results at a) 2 hours, b) 15 hours, c) 37 hours, and d) 65 hours after stimuli were introduced. For each timepoint half violin plots with individual data points are shown, where the relative size (area) of each data point represents the relative weighting factor based on cell object area. Normalized αSMA reporter intensity weighted by cell area, with mean ± SE; n = 3 independent samples, with measurements from ≥48 cell objects per condition at each timepoint; Significance determined via one-way ANOVA followed by Tukey's post-hoc test: * indicates a significant difference, where * $p < 0.05$, ** $p < 0.01$, *** $p < 0.001$. Statistical analyses within each condition over time are available in **Table S4**.

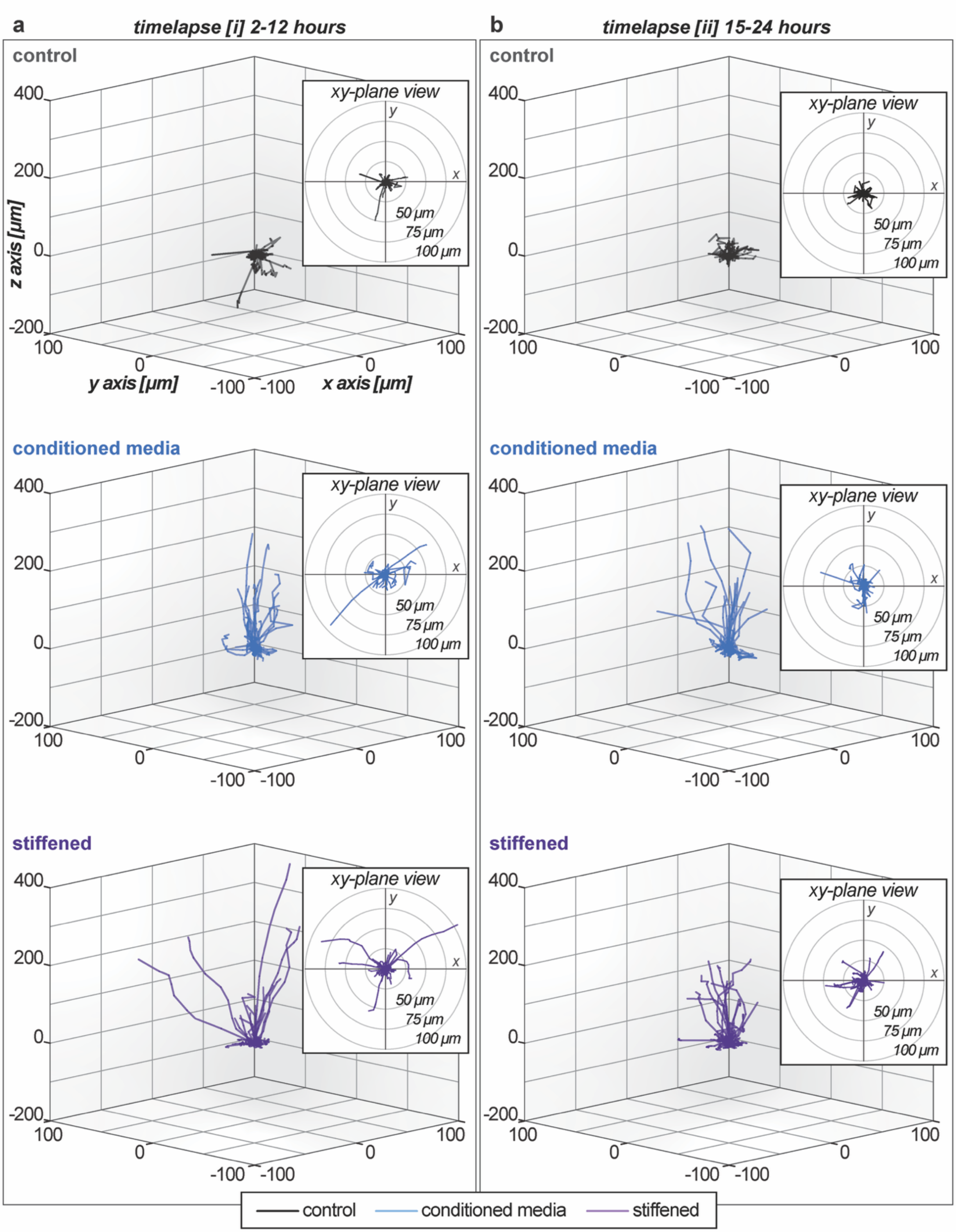

**Figure S16. Fibroblast motility: cell tracks separated by condition at timelapses *[i]* and *[ii]*.** 3D traces of cell tracks over the a) 2–12-hour and b) 15–24-hour timelapses (time from introduction of stimuli). Each identified cell object (individual cell or cluster of multiple cells) was tracked in three dimensions over the indicated time frame to determine measurements of motility, with insets of the *xy*-plane view (2D top-down). (n = 3 independent samples with measurements from ≥47 ([*i*]) or ≥52 ([*ii*]) cell objects per condition.)

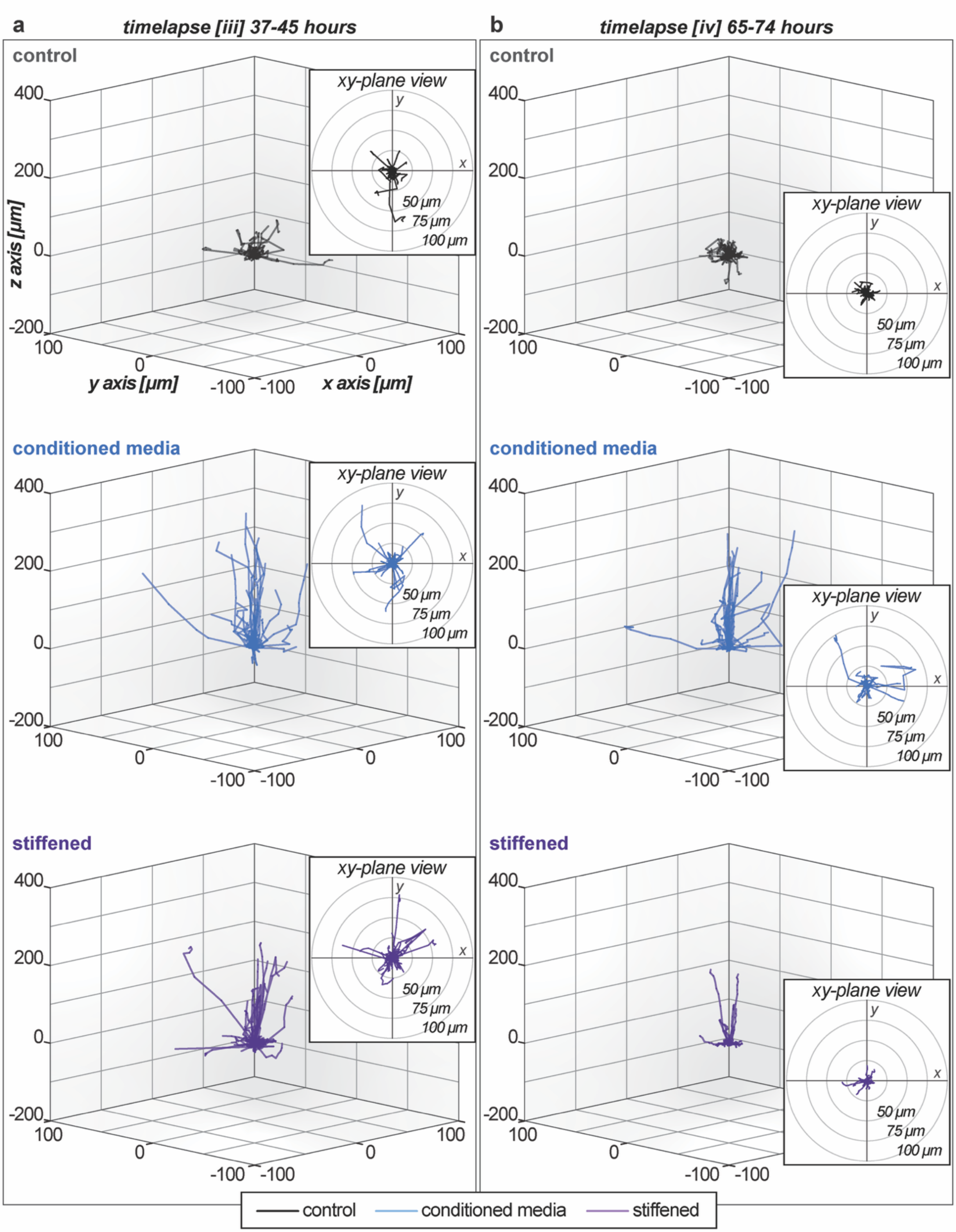

**Figure S17. Fibroblast motility: cell tracks separated by condition at timelapses *[iii]* and *[iv]*.** 3D traces of cell tracks over the a) 37–45-hour and b) 65–74-hour timelapses (time from introduction of stimuli). Each identified cell object (individual cell or cluster of multiple cells) was tracked in three dimensions over the indicated time frame to determine measurements of motility, with insets of the *xy*-plane view (2D top-down). (n = 3 independent samples with measurements from ≥51 ([*iii*]) or ≥36 ([*iv*]) cell objects per condition.)

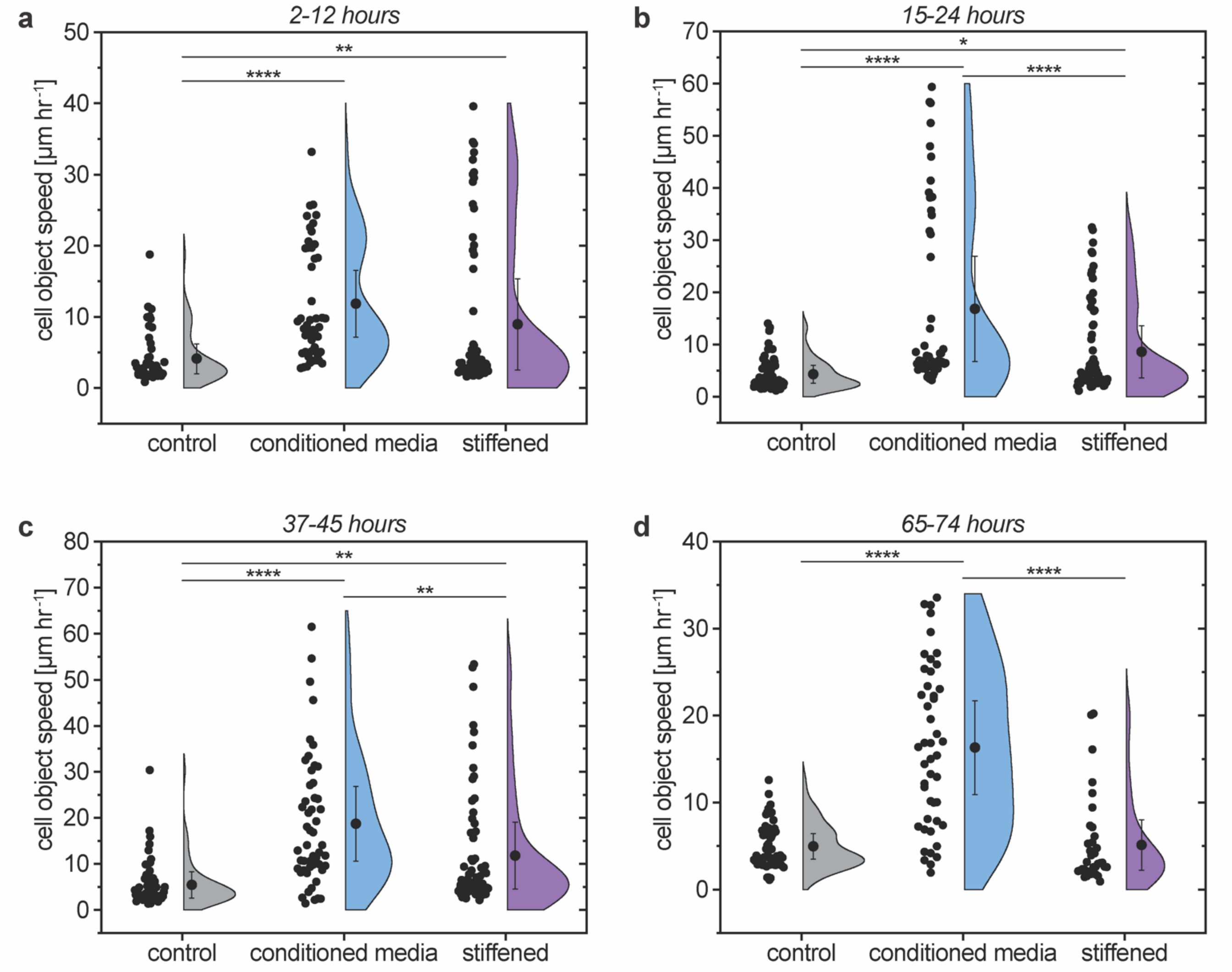

**Figure S18. Fibroblast motility speed and Tukey's post-hoc multiple comparisons at each timepoint.** Speed for each cell object tracked over the a) 2–12-hour, b) 15–24-hour, c) 37–45-hour, and d) 65–74-hour timelapses (time from introduction of stimuli). Each identified cell object (individual cell or cluster of multiple cells) was tracked in three dimensions and cell object speed was determined and are presented here as half violin plots with individual data points for each tracked object. (Mean ± SE; n = 3 independent samples with measurements from ≥47 (a), ≥52 (b), ≥51 (c), or ≥36 (d) cell objects per condition; Significance determined via one-way ANOVA followed by Tukey's post-hoc test: * indicates a significant difference, where * $p < 0.05$, ** $p < 0.01$, *** $p < 0.001$, **** $p < 0.0001$.) Statistical analyses within each condition over time are available in **Table S5**.

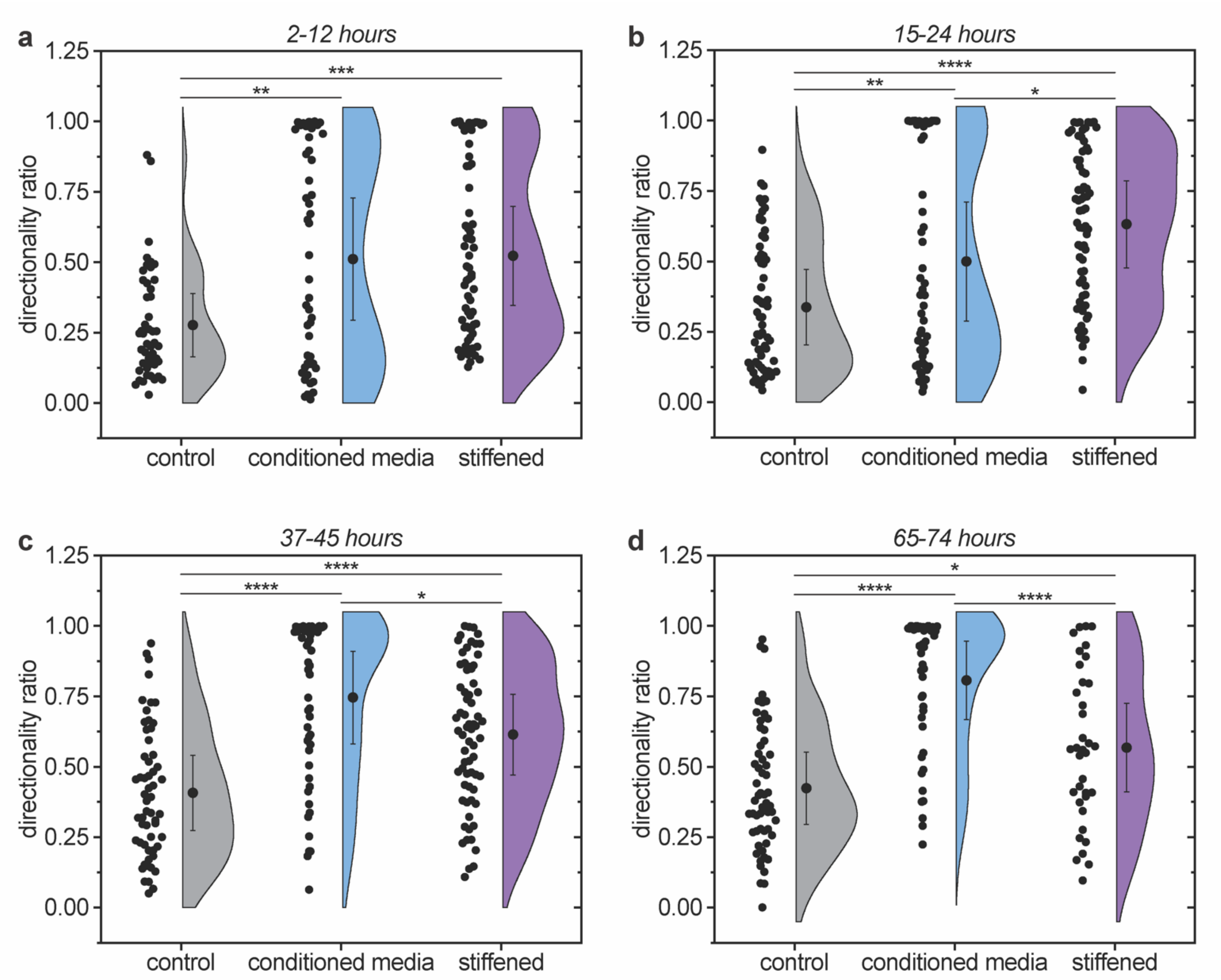

**Figure S19. Fibroblast directional persistence and Tukey's post-hoc multiple comparisons at each timepoint.** Directional persistence (measured by directionality ratio) for each cell object tracked over the a) 2–12-hour, b) 15–24-hour, c) 37–45-hour, and d) 65–74-hour timelapses (time from introduction of stimuli). Each identified cell object (individual cell or cluster of multiple cells) was tracked in three dimensions and the directionality ratio (ratio of displacement, *d*, to total distance traveled, *D*) of each cell object was calculated as a measurement of directional persistence. Values closer to 0 are indicative of meandering cell movement (low directional persistence), whereas values closer to 1 are indicative of more directional cell movement (high directional persistence). Directionality ratio results are presented here as half violin plots with individual data points for each tracked object. (Mean ± SE; n = 3 independent samples with measurements from ≥47 (a), ≥52 (b), ≥51 (c), or ≥36 (d) cell objects per condition; Significance determined via one-way ANOVA followed by Tukey's post-hoc test: * indicates a significant difference, where * $p < 0.05$, ** $p < 0.01$, *** $p < 0.001$, **** $p < 0.0001$.) Statistical analyses within each condition over time are available in **Table S6**.

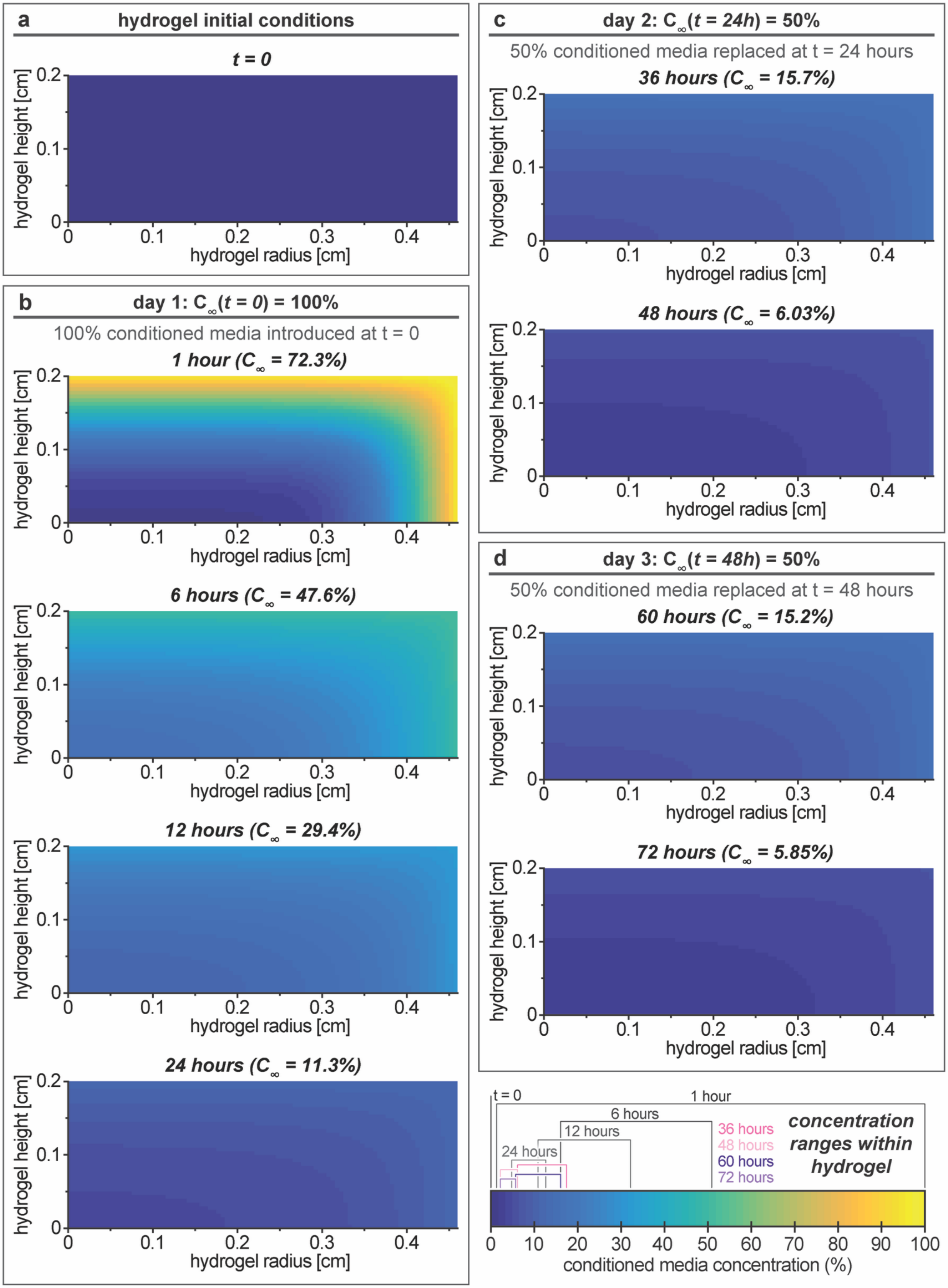
a
hydrogel initial conditions
t = 0
b
day 1: C∞(t = 0) = 100%
100% conditioned media introduced at t = 0
1 hour (C∞ = 72.3%)
6 hours (C∞ = 47.6%)
12 hours (C∞ = 29.4%)
24 hours (C∞ = 11.3%)
c
day 2: C∞(t = 24h) = 50%
50% conditioned media replaced at t = 24 hours
36 hours (C∞ = 15.7%)
48 hours (C∞ = 6.03%)
d
day 3: C∞(t = 48h) = 50%
50% conditioned media replaced at t = 48 hours
60 hours (C∞ = 15.2%)
72 hours (C∞ = 5.85%)
hydrogel height [cm]
hydrogel radius [cm]
t = 0
1 hour
6 hours
12 hours
24 hours
36 hours
48 hours
60 hours
72 hours
concentration ranges within hydrogel
conditioned media concentration (%)

**Figure S20. Conditioned media reaction-diffusion modeling results.** Results represent a slice of the hydrogel, from the axis of symmetry ($r = 0$) to the outer radius of the hydrogel ($r = R$) and from the bottom of the hydrogel ($z = 0$) to the top ($z = H$). a) Initial conditions. b) Day 1 incubation with 100% conditioned media. c) Day 2 incubation with 50% conditioned media. d) Day 3 incubation with 50% conditioned media. Concentration ranges within the hydrogel are indicated for each timepoint on the heatmap legend. Note: $C_\infty$ for each incubation decreases over time according to the protein flux into the hydrogel and the rate of degradation.

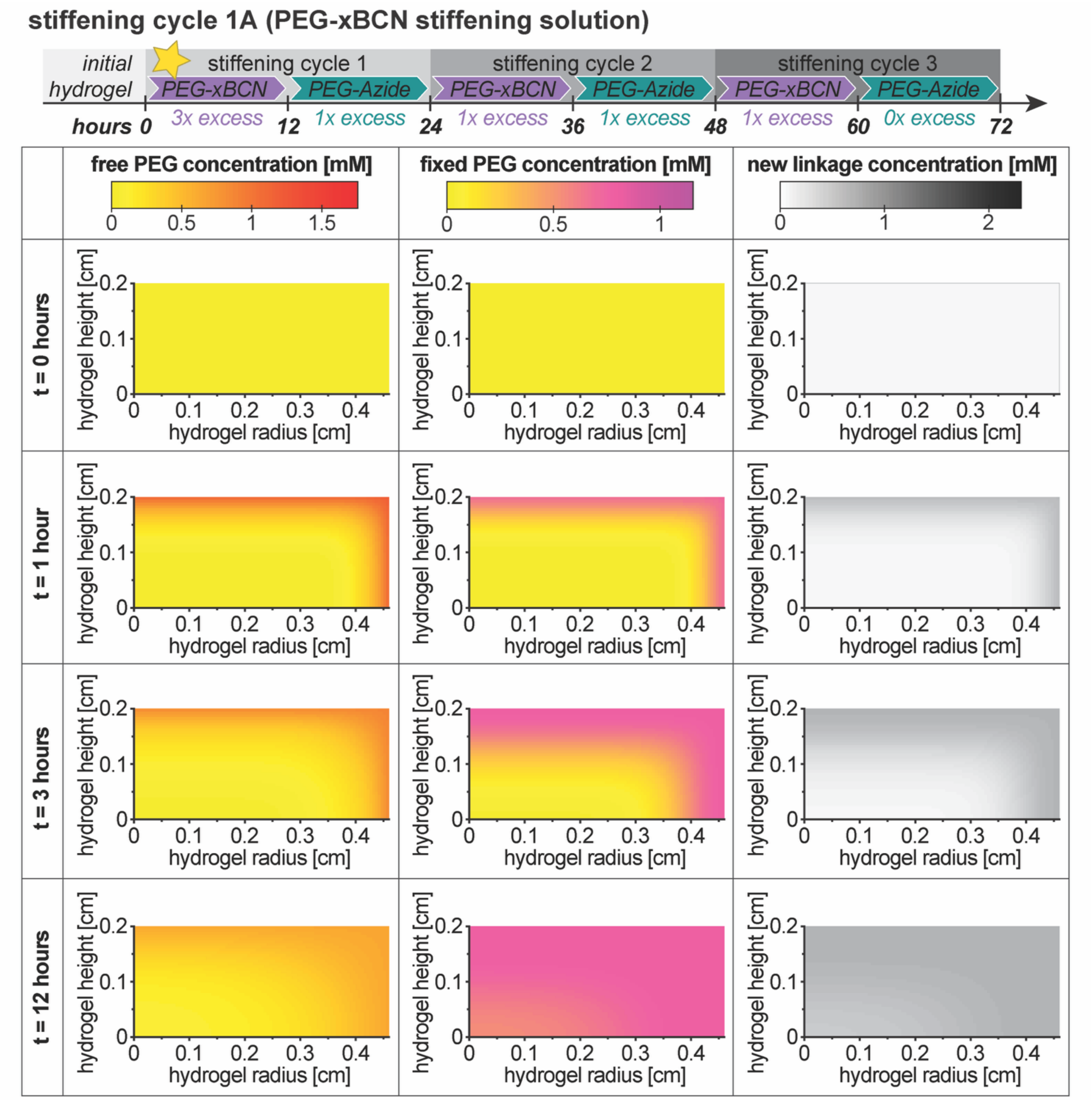

**Figure S21. Stiffening cycle 1A modeling results.** Hydrogel concentration results for free (diffusing) PEG-xBCN, PEG-xBCN that has been crosslinked and fixed to the hydrogel, and newly formed linkages within the stiffening cycle 1A incubation. Results shown for select timepoints throughout the 12-hour incubation. The number of "available" functional groups per PEG molecule ($f = 1$) was assigned according to the stoichiometric excess of PEG functional groups in the hydrogel at equilibrium. Although the new linkage concentration ranges between 0.529-0.770 mM (inside-to-outside of gel), each PEG group will theoretically be tethered to the hydrogel by a single linkage and the PEG groups will not contribute any new crosslinks at this time.

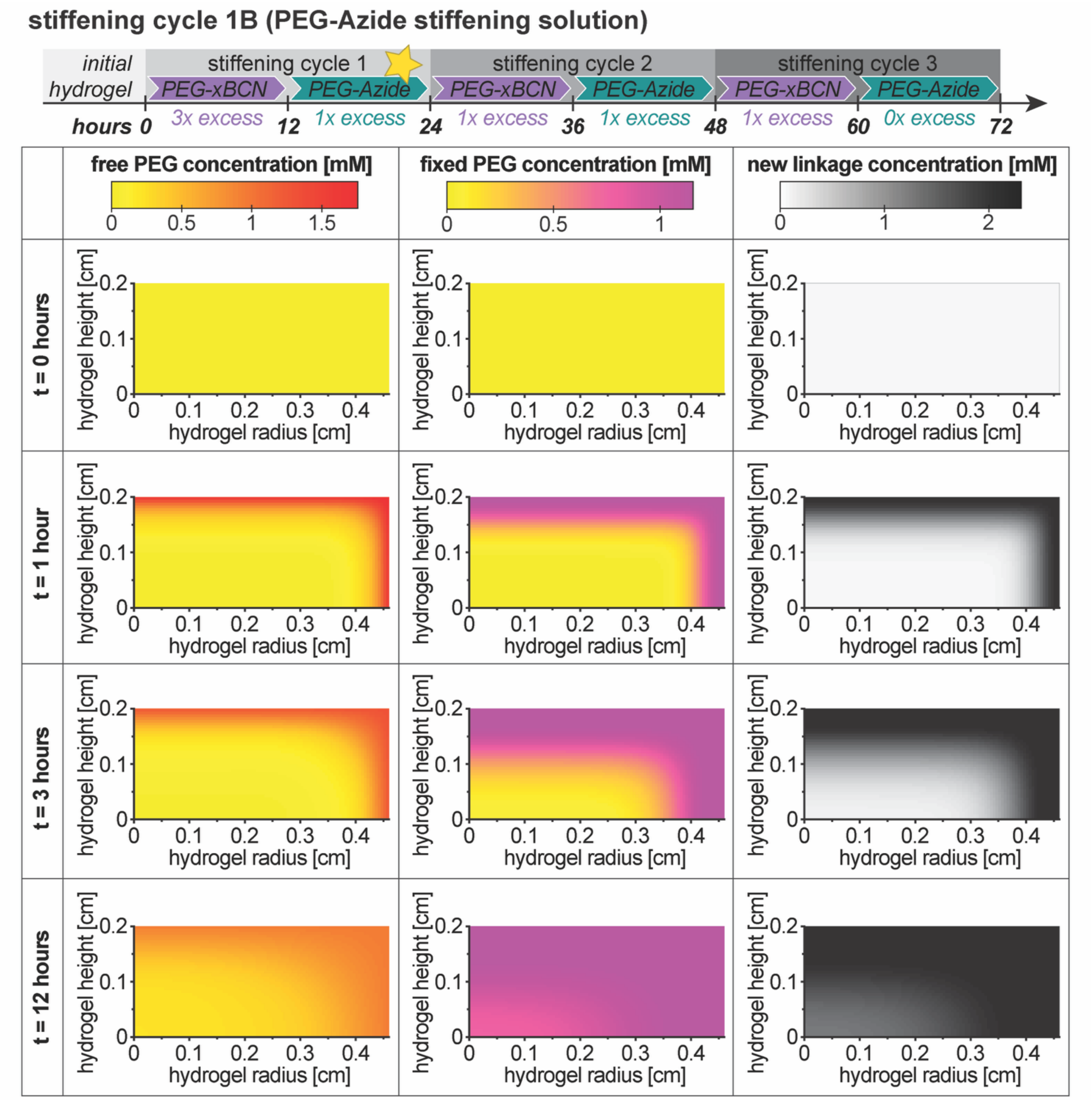

**Figure S22. Stiffening cycle 1B modeling results.** Hydrogel concentration results for free (diffusing) PEG-Azide, PEG-Azide that has been crosslinked and fixed to the hydrogel, and newly formed linkages within the stiffening cycle 1B incubation. Results shown for select timepoints throughout the 12-hour incubation. The number of "available" functional groups per PEG molecule ($f = 2$) was assigned according to the stoichiometric excess of PEG functional groups in the hydrogel at equilibrium. Within this 'incubation', the new linkage concentration ranges between 1.51-2.31 mM (inside-to-outside of gel). The PEG groups from incubations 1A and 1B will be contributing new crosslinks to the hydrogel, for a predicted cumulative new crosslink density ($\rho_x$) range of 1.28-1.93 mM (inside-to-outside of gel).

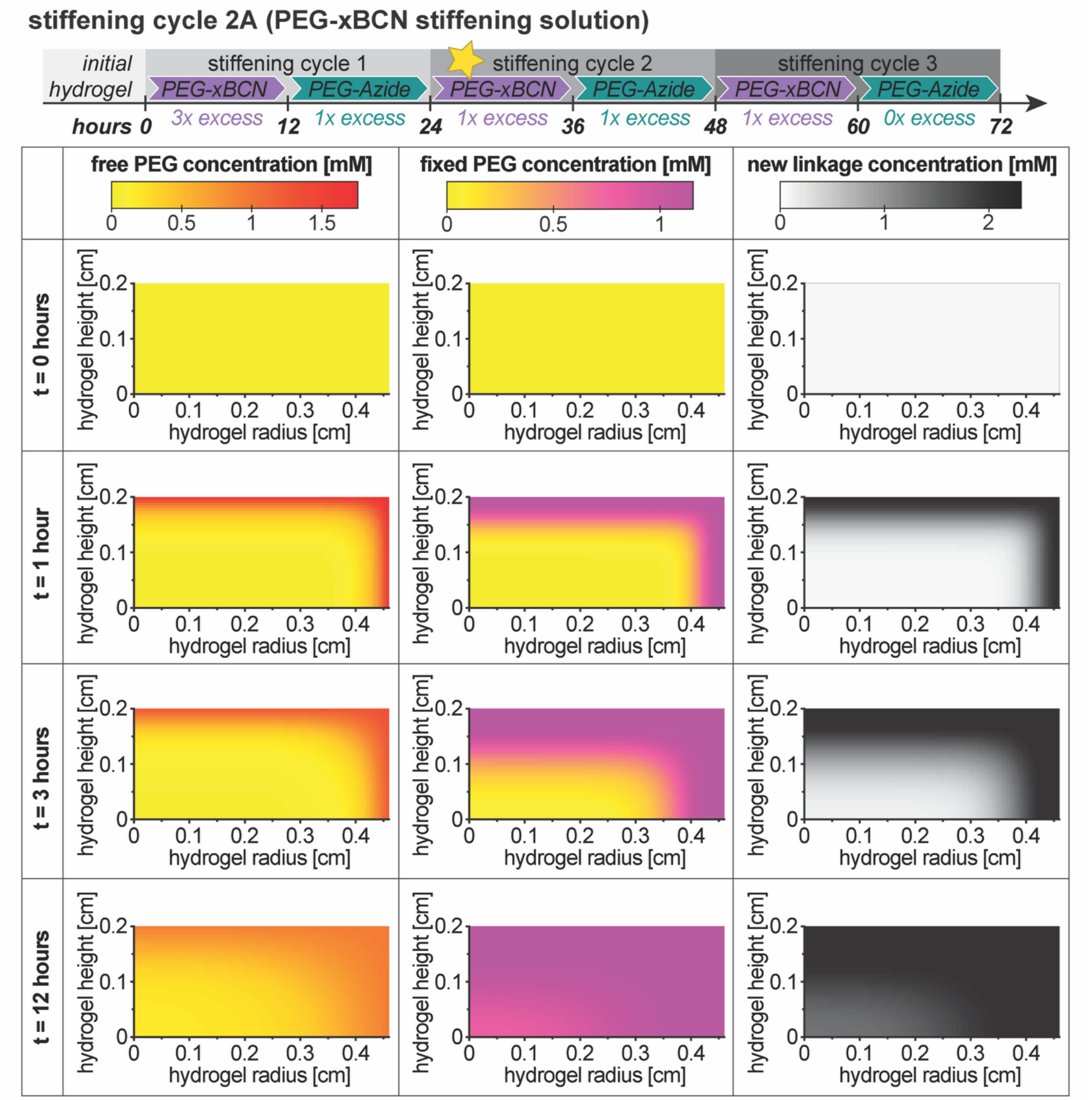

**Figure S23. Stiffening cycle 2A modeling results.** Hydrogel concentration results for free (diffusing) PEG-xBCN, PEG-xBCN that has been crosslinked and fixed to the hydrogel, and newly formed linkages within the stiffening cycle 2A incubation. Results shown for select timepoints throughout the 12-hour incubation. The number of "available" functional groups per PEG molecule ($f = 2$) was assigned according to the stoichiometric excess of PEG functional groups in the hydrogel at equilibrium. Within this 'incubation', the new linkage concentration ranges between 1.60-2.31 mM (inside-to-outside of gel). The PEG groups from incubations 1A through 2A will be contributing new crosslinks to the hydrogel, for a predicted cumulative new crosslink density ($\rho_x$) range of 2.08-3.08 mM (inside-to-outside of gel).

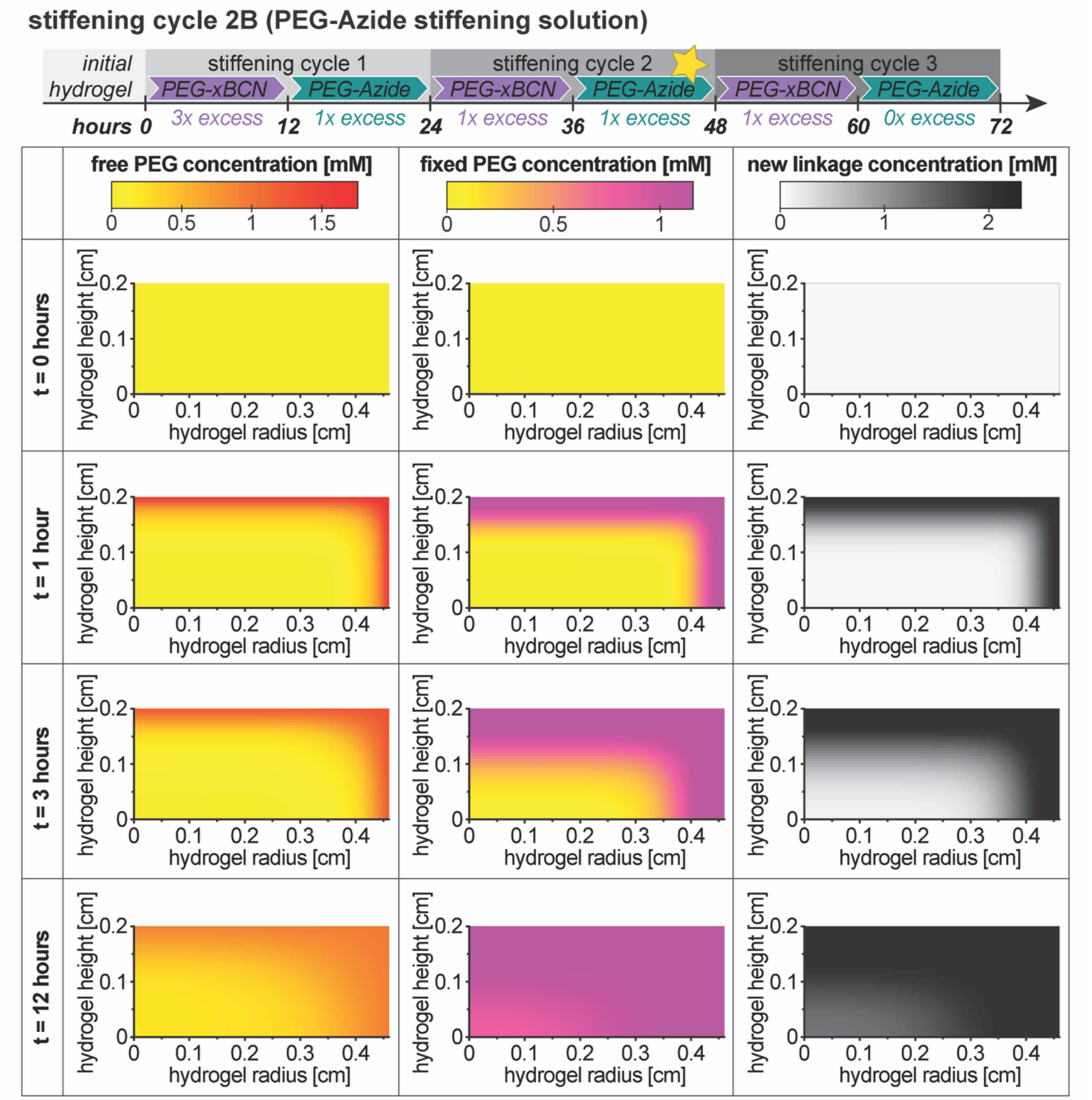

**Figure S24. Stiffening cycle 2B modeling results.** Hydrogel concentration results for free (diffusing) PEG-Azide, PEG-Azide that has been crosslinked and fixed to the hydrogel, and newly formed linkages within the stiffening cycle 2B incubation. Results shown for select timepoints throughout the 12-hour incubation. The number of "available" functional groups per PEG molecule ($f$ = 2) was assigned according to the stoichiometric excess of PEG functional groups in the hydrogel at equilibrium. Within this 'incubation', the new linkage concentration ranges between 1.56-2.31 mM (inside-to-outside of gel). The PEG groups from incubations 1A through 2B will be contributing new crosslinks to the hydrogel, for a predicted cumulative new crosslink density ($\rho_x$) range of 2.86-4.24 mM (inside-to-outside of gel).

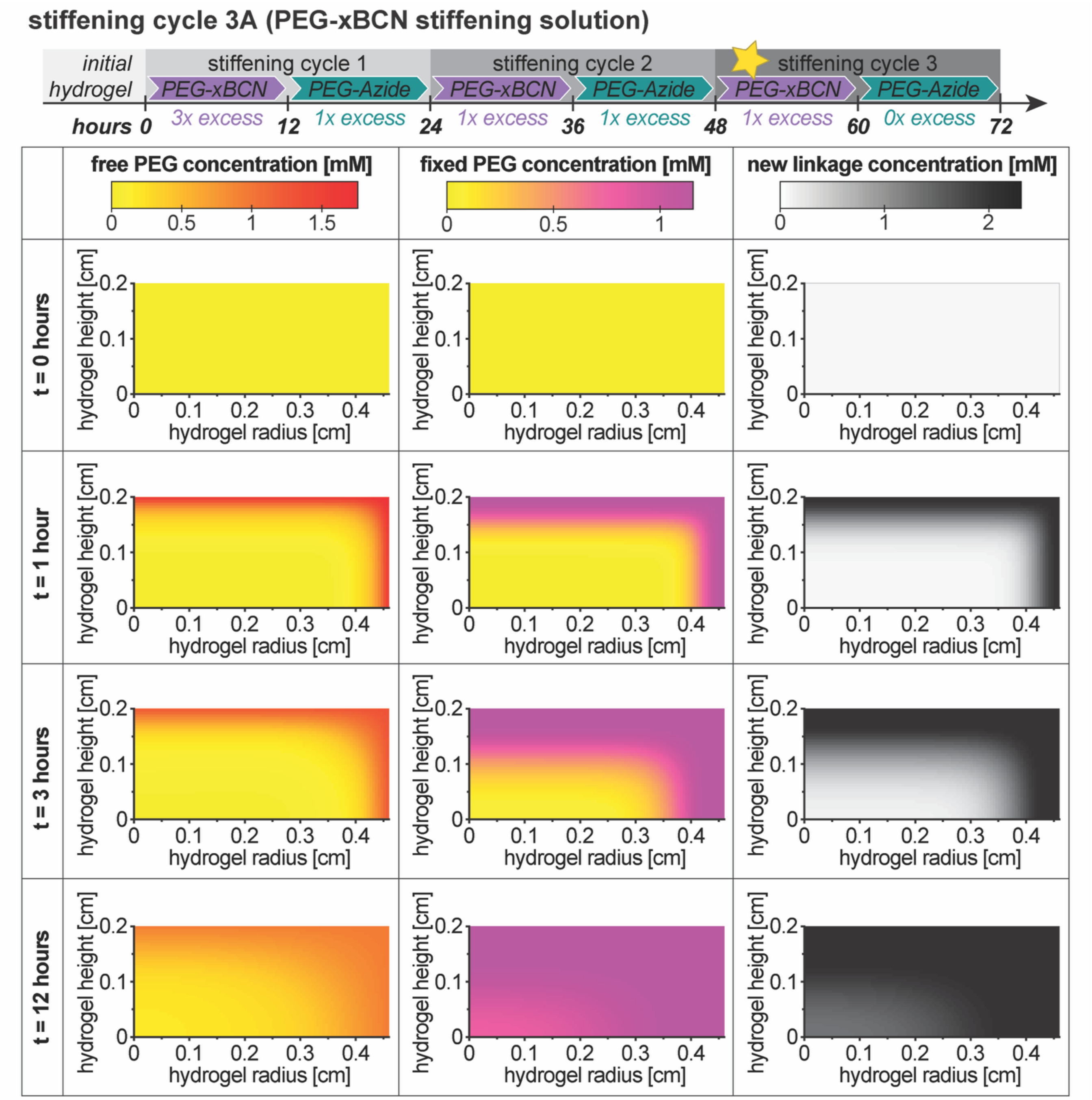

**Figure S25. Stiffening cycle 3A modeling results.** Hydrogel concentration results for free (diffusing) PEG-xBCN, PEG-xBCN that has been crosslinked and fixed to the hydrogel, and newly formed linkages within the stiffening cycle 3A incubation. Results shown for select timepoints throughout the 12-hour incubation. The number of "available" functional groups per PEG molecule ($f = 2$) was assigned according to the stoichiometric excess of PEG functional groups in the hydrogel at equilibrium. Within this 'incubation', the new linkage concentration ranges between 1.58-2.31 mM (inside-to-outside of gel). The PEG groups from incubations 1A through 3A will be contributing new crosslinks to the hydrogel, for a predicted cumulative new crosslink density ($\rho_x$) range of 3.65-5.39 mM (inside-to-outside of gel).

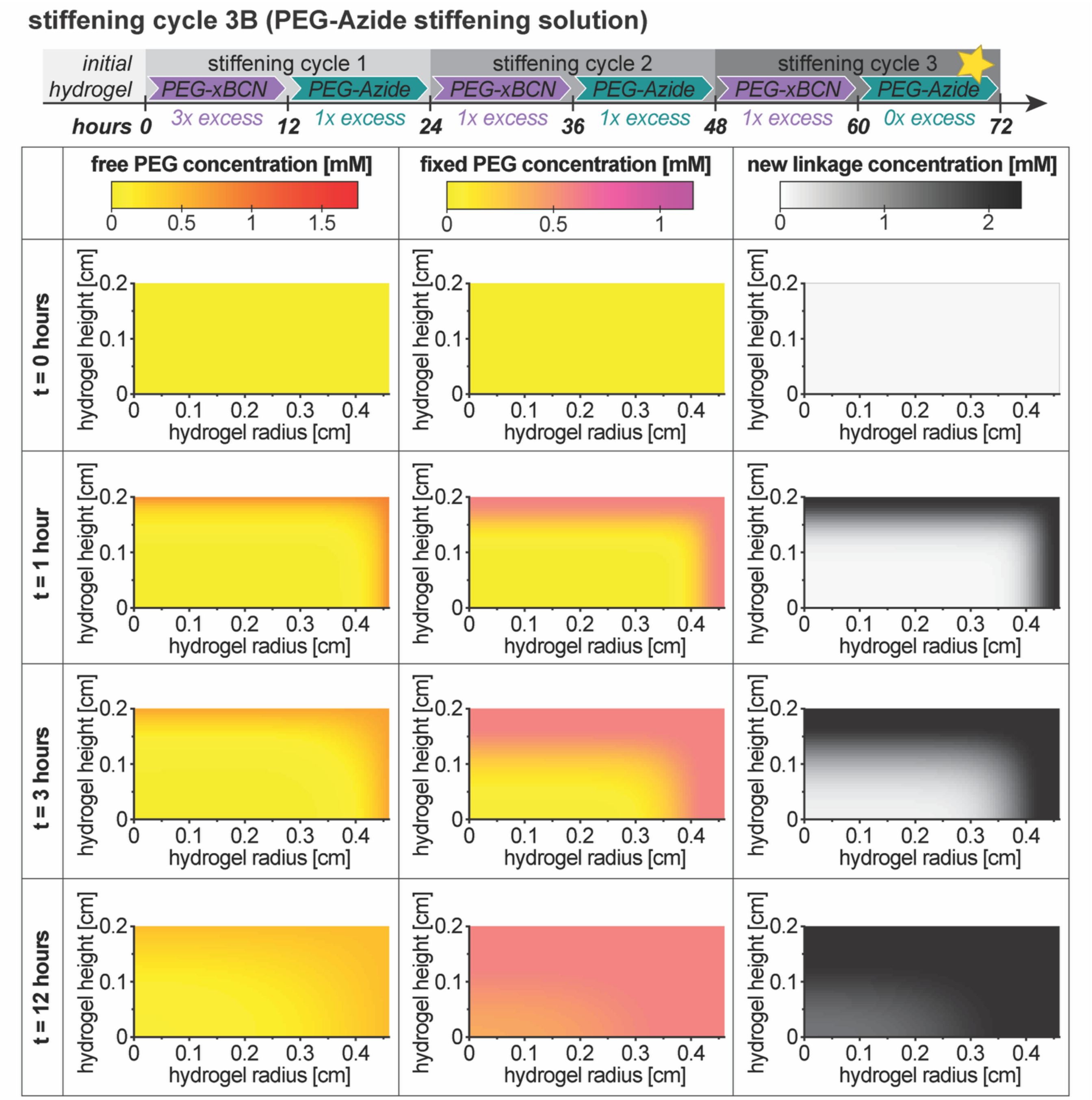

**Figure S26. Stiffening cycle 3B modeling results.** Hydrogel concentration results for free (diffusing) PEG-Azide, PEG-Azide that has been crosslinked and fixed to the hydrogel, and newly formed linkages within the stiffening cycle 3B incubation. Results shown for select timepoints throughout the 12-hour incubation. The number of "available" functional groups per PEG molecule ($f = 4$) was assigned according to the stoichiometric excess of PEG functional groups in the hydrogel at equilibrium. Within this 'incubation', the new linkage concentration ranges between 1.57-2.31 mM (inside-to-outside of gel). The PEG groups from incubations 1A through 3B will be contributing new crosslinks to the hydrogel, for a predicted cumulative new crosslink density ($\rho_x$) range of 4.04-5.97 mM (inside-to-outside of gel).

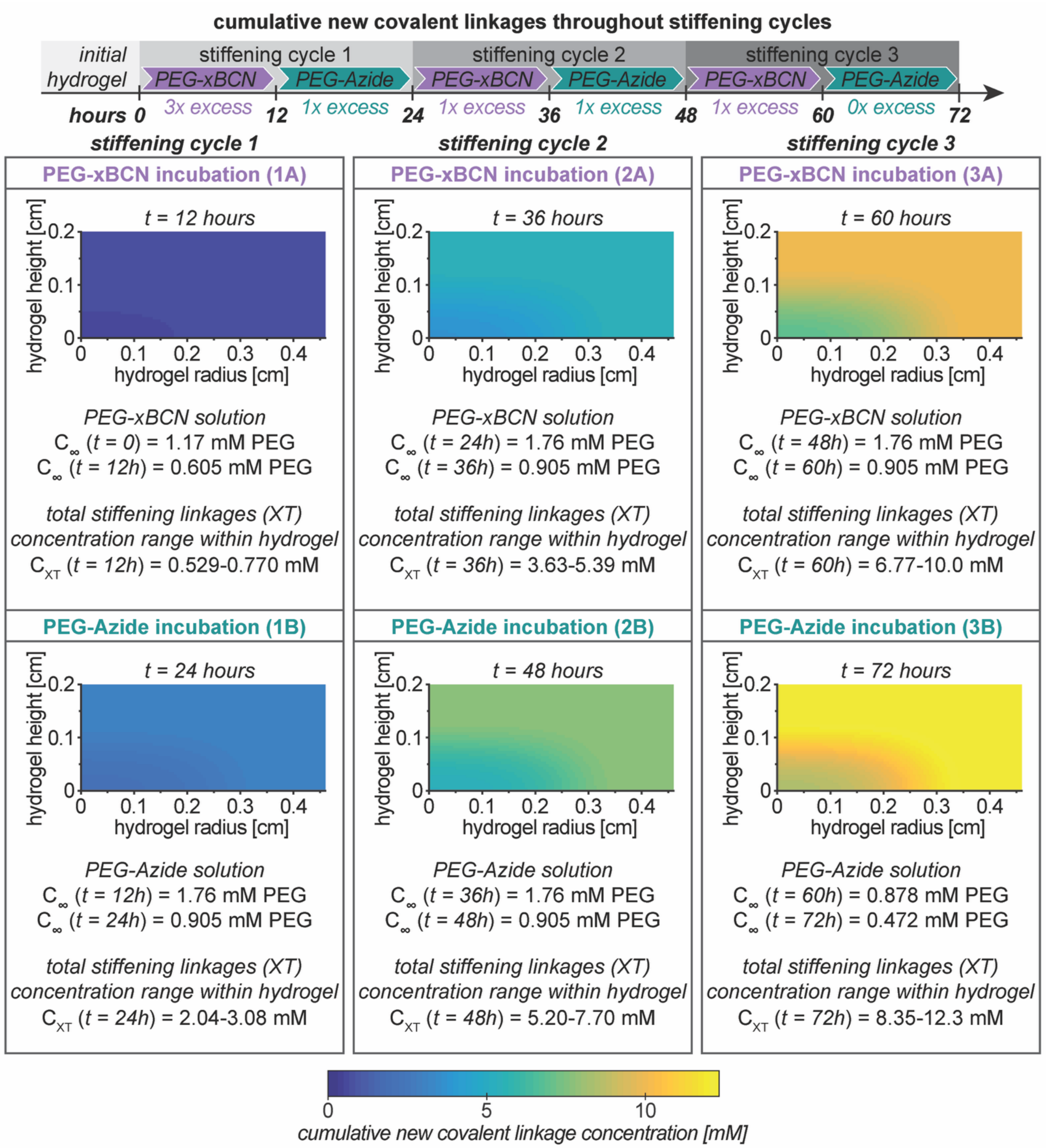

**Figure S27. Model results: cumulative new covalent linkages over the course of stiffening.** Total concentration of linkages formed during the stiffening process (with concentration ranges throughout the hydrogel indicated for each timepoint). PEG solution concentrations are also indicated for the start and end of each incubation, where $C_\infty$ decreases throughout each incubation as the PEG diffuses into the hydrogel. Results shown for the end of each incubation.

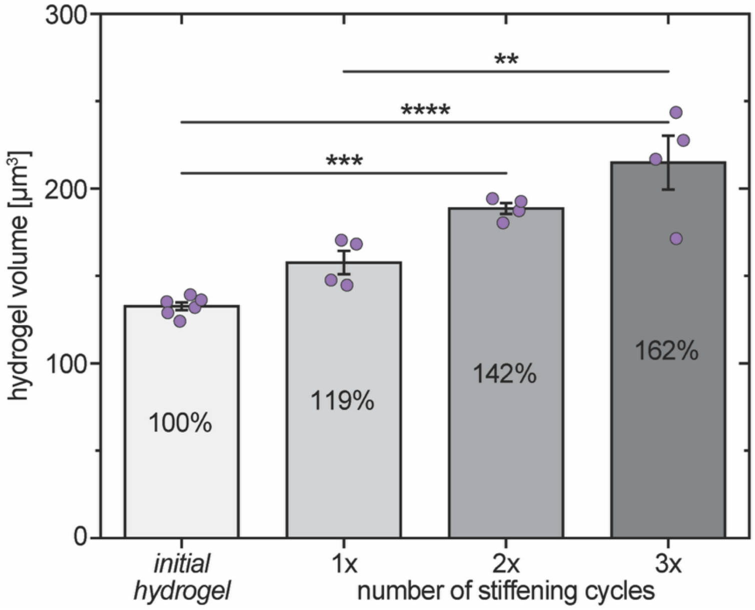

**Figure S28. Hydrogel volume over the course of stiffening.** Hydrogel volumes were measured prior to stiffening (after swelling) and after 1, 2, or 3 cycles of stiffening. Percentages represent the volume after each cycle relative to the initial hydrogel. All incubations and measurements were conducted at 37 °C. (Mean ± SE with individual datapoints; n ≥ 4 independent samples for each condition; Significance determined via one-way ANOVA followed by Tukey's post-hoc test: * $p < 0.05$, ** $p < 0.01$, *** $p < 0.001$, **** $p < 0.0001$.)

#### 5. Supplemental Tables

**Table S1. Initial hydrogel component concentrations.**

| **Component Information** | | | **Stock Solutions** | **Hydrogel at Preparation** | |
|---|---|---|---|---|---|
| *Component Identity* | *Functional Group(s)* | *Component Functionality* | *Functional Group (mM)* | *Functional Group (mM)* | *Component (mM)* |
| PEG (10 wt%) | Thiol | 4 | 55 | 20 | 5 |
| Linker Peptide | Alloc | 2 | 71.1 | 13 | 6.5 |
| mfCMP | Alloc | 1 | 5 | 5 | 5 |
| | Azide | 1 | 5 | 5 | |
| RGDS | Alloc | 1 | 43.3 | 2 | 2 |
| LAP | - | - | 40 | - | 2.2 |

**Table S2. Stiffening solution concentrations.**

| | **Hydrogel Before Reaction** | | **Stiffening Monomer and Functional Group Concentrations in Solution** | | | **Hydrogel After Reaction** | |
|---|---|---|---|---|---|---|---|
| | Available Functional Group | [Available Functional Group] | Stiffening Polymer Identity | [Stiffening Solution] (250 µL*) | [Stiffening Solution] (380 µL**) | Available Functional Group | [Available Functional Group] |
| **Cycle 1A** | Azide | 0.77 mM | PEG-xBCN | 4.68 mM [BCN] | 3.08 mM [BCN] | BCN | 2.31 mM |
| **Cycle 1B** | BCN | 2.31 mM | PEG-Azide | 7.02 mM [Azide] | 4.62 mM [Azide] | Azide | 2.31 mM |
| **Cycle 2A** | Azide | 2.31 mM | PEG-xBCN | 7.02 mM [BCN] | 4.62 mM [BCN] | BCN | 2.31 mM |
| **Cycle 2B** | BCN | 2.31 mM | PEG-Azide | 7.02 mM [Azide] | 4.62 mM [Azide] | Azide | 2.31 mM |
| **Cycle 3A** | Azide | 2.31 mM | PEG-xBCN | 7.02 mM [BCN] | 4.62 mM [BCN] | BCN | 2.31 mM |
| **Cycle 3B** | BCN | 2.31 mM | PEG-Azide | 3.51 mM [Azide] | 2.31 mM [Azide] | Azide | 0 mM*** |

* Volume of added stiffening solution = 250 µL

** Volume of equilibrium swollen hydrogel (130 µL) + added solution (250 µL) = 380 µL; Assumes that the stiffening monomer diffuses equally between hydrogel and surrounding solution

*** Azide and Alkyne groups should be at stoichiometric ratios after the final stiffening step

Note: the number of "available" functional groups per PEG monomer ($f$) was calculated for each cycle as $f = 4 \times a / b$, where

$a$ = **Hydrogel Before Reaction**
→ [Available Functional Group]

$b$ = **Stiffening Monomer and Functional Group Concentrations in Solution**
→ [Stiffening Solution] (380 µL)

**Table S3. Statistical analysis of metabolic activity over time (within each condition).**

| Control | | Conditioned Media | | Stiffened | |
|---|---|---|---|---|---|
| ANOVA p = 0.928 | | ANOVA ***p = 0.0000725*** | | ANOVA ***p = 0.0262*** | |
| *Day Comparison* | *p =* | *Day Comparison* | *p =* | *Day Comparison* | *p =* |
| -5 vs. -1 | 0.99994 | -5 vs. -1 | 0.890 | -5 vs. -1 | 0.960 |
| -5 vs. 3 | 0.930 | ***-5 vs. 3*** | ***0.000634*** | -5 vs. 3 | 0.341 |
| -5 vs. 5 | 0.998 | ***-5 vs. 5*** | ***0.000780*** | -5 vs. 5 | 0.224 |
| -1 vs. 3 | 0.945 | ***-1 vs. 3*** | ***0.000317*** | -1 vs. 3 | 0.582 |
| -1 vs. 5 | 0.9993 | ***-1 vs. 5*** | ***0.000385*** | -1 vs. 5 | 0.115 |
| 3 vs. 5 | 0.972 | 3 vs. 5 | 0.997 | ***3 vs. 5*** | ***0.0185*** |

**Table S4. Statistical analysis of αSMA expression over time (within each condition).**

| Control | | Conditioned Media | | Stiffened | |
|---|---|---|---|---|---|
| ANOVA ***p = 0.0073182*** | | ANOVA ***p = 2.30E-12*** | | ANOVA ***p = 2.02E-42*** | |
| *Day Comparison* | *p =* | *Day Comparison* | *p =* | *Day Comparison* | *p =* |
| ***-5 vs. -1*** | ***0.0290*** | ***-5 vs. -1*** | ***0.0000193*** | -5 vs. -1 | 0.938 |
| -5 vs. 3 | 0.9999998 | -5 vs. 3 | 0.115 | ***-5 vs. 3*** | ***3.77E-09*** |
| -5 vs. 5 | 0.999 | -5 vs. 5 | 0.992 | ***-5 vs. 5*** | ***3.77E-09*** |
| ***-1 vs. 3*** | ***0.0239*** | ***-1 vs. 3*** | ***3.77E-09*** | ***-1 vs. 3*** | ***3.77E-09*** |
| ***-1 vs. 5*** | ***0.0207*** | ***-1 vs. 5*** | ***4.32E-06*** | ***-1 vs. 5*** | ***3.77E-09*** |
| 3 vs. 5 | 0.99898169 | ***3 vs. 5*** | ***0.0214*** | ***3 vs. 5*** | ***0.0000784*** |

**Table S5. Statistical analysis of fibroblast speed over time (within each condition).**

| Control | | Conditioned Media | | Stiffened | |
|---|---|---|---|---|---|
| ANOVA p = 0.186 | | ANOVA p = 0.0645 | | ANOVA ***p = 0.0161*** | |
| *Time Comparison (h)* | *p =* | *Time Comparison (h)* | *p =* | *Time Comparison (h)* | *p =* |
| 2-12 vs. 15-24 | 0.995 | 2-12 vs. 15-24 | 0.220 | 2-12 vs. 15-24 | 0.997 |
| 2-12 vs. 37-45 | 0.240 | 2-12 vs. 37-45 | ***0.0423*** | 2-12 vs. 37-45 | 0.359 |
| 2-12 vs. 65-74 | 0.625 | 2-12 vs. 65-74 | 0.341 | 2-12 vs. 65-74 | 0.264 |
| 15-24 vs. 37-45 | 0.284 | 15-24 vs. 37-45 | 0.881 | 15-24 vs. 37-45 | 0.246 |
| 15-24 vs. 65-74 | 0.722 | 15-24 vs. 65-74 | 0.997 | 15-24 vs. 65-74 | 0.339 |
| 37-45 vs. 65-74 | 0.892 | 37-45 vs. 65-74 | 0.796 | 37-45 vs. 65-74 | ***0.00779*** |

**Table S6. Statistical analysis of fibroblast directional persistence over time (within each condition).**

| Control | | Conditioned Media | | Stiffened | |
|---|---|---|---|---|---|
| ANOVA ***p = 0.00274*** | | ANOVA ***p = 0.000000403*** | | ANOVA p = 0.0909 | |
| *Time Comparison (h)* | *p =* | *Time Comparison (h)* | *p =* | *Time Comparison (h)* | *p =* |
| 2-12 vs. 15-24 | 0.477 | 2-12 vs. 15-24 | 0.998 | 2-12 vs. 15-24 | 0.0851 |
| 2-12 vs. 37-45 | ***0.0152*** | 2-12 vs. 37-45 | ***0.00171*** | 2-12 vs. 37-45 | 0.205 |
| 2-12 vs. 65-74 | ***0.00414*** | 2-12 vs. 65-74 | ***0.0000533*** | 2-12 vs. 65-74 | 0.851 |
| 15-24 vs. 37-45 | 0.306 | 15-24 vs. 37-45 | ***0.000604*** | 15-24 vs. 37-45 | 0.981 |
| 15-24 vs. 65-74 | 0.136 | 15-24 vs. 65-74 | ***0.0000143*** | 15-24 vs. 65-74 | 0.659 |
| 37-45 vs. 65-74 | 0.978 | 37-45 vs. 65-74 | 0.787 | 37-45 vs. 65-74 | 0.843 |

**Table S7. Model parameter values.**

| *Hydrogel Parameters* | | | | | |
|---|---|---|---|---|---|
| Parameter | Symbol | MATLAB Variable | Units | Value | Source |
| Radius | $R$ | `R` | cm | 0.46 | † |
| Height | $H$ | `H` | cm | 0.20 | † |
| Solution volume | $V_S$ | `V` | $cm^3$ | 0.25 | † |
| Discretized radius length | $\Delta r$ | `delta_r` | cm | 0.005 | ‡ |
| Discretized height length | $\Delta z$ | `delta_z` | cm | 0.005 | ‡ |
| ***Conditioned Media Parameters (using PDGF as the model protein)*** | | | | | |
| Parameter | Symbol | MATLAB Variable | Units | Value | Source |
| Sum limit over m | $M$ | `SumLimitM` | - | 100 | ‡ |
| Sum limit over n | $N$ | `SumLimitN` | - | 100 | ‡ |
| Initial solution concentration | $C_0$ | `C_i` | % ($mmol\ cm^{-3}$) | 100, 50 | † |
| Diffusivity in water | $D_\infty$ | `D_water` | $cm^2\ s^{-1}$ | 9.38E-07 | § |
| Effective diffusivity | $D_{eff}$ | `D` | $cm^2\ s^{-1}$ | 7.99E-07 | § |
| PDGF half-life | $t_{1/2}$ | `t_half` | h | 2.4 | [34] |
| ***PEG Stiffening Parameters**** | | | | | |
| Parameter | Symbol | MATLAB Variable | Units | Value | Source |
| Sum limit over m (r nodes) | $M$ | `N` | - | 93 | ‡ |
| Sum limit over n (z nodes) | $N$ | `M` | - | 41 | ‡ |
| Diffusivity in water | $D_\infty$ | `D_water` | $cm^2\ s^{-1}$ | 4.81E-07 | § |
| Effective diffusivity | $D_{eff}$ | `D` | $cm^2\ s^{-1}$ | 2.80E-07 | § |
| Rate constant | $k$ | `k` | $cm^3\ mmol^{-1}\ s^{-1}$ | 0.57 | [24] |
| Discretized time step | $\Delta t$ | `delta_t` | s | 1 | ‡ |
| Initial solution concentration | $C_0$ | `C_i` | $mmol\ cm^{-3}$ | variable | † |
| Initial azide concentration | $C_{A,0}$ | `C_A0` | $mmol\ cm^{-3}$ | 7.7E-04 | † |

† Values determined according to experimental design and measurements
‡ Values selected according to brief sensitivity analysis for this system
§ Values calculated in Supplemental Calculations
* Note: in MATLAB code, i and N apply to the *r*-direction, while j and M apply to the *z*-direction
